# Varying molecular interactions explain crowder-dependent enzyme function of a viral protease

**DOI:** 10.1101/2022.12.16.520723

**Authors:** Natalia Ostrowska, Michael Feig, Joanna Trylska

**Affiliations:** Centre of New Technologies, University of Warsaw, Warsaw, Poland; Department of Biochemistry and Molecular Biology, Michigan State University, East Lansing, MI, USA

**Keywords:** Macromolecular crowding, synthetic crowders, Ficoll, polyethylene glycol (PEG), viral protease NS3/4A, molecular dynamics simulations

## Abstract

Biochemical processes in cells, including enzyme-catalyzed reactions, occur in crowded conditions with various background macromolecules occupying up to 40% of cytoplasm’s volume. Viral enzymes in the host cell also encounter such crowded conditions as they often function at the endoplasmic reticulum membranes. We focus on an enzyme encoded by the hepatitis C virus, the NS3/4A protease, which is crucial for viral replication. We have previously found experimentally that synthetic crowders, polyethylene glycol (PEG) and branched polysucrose (Ficoll), differently affect the kinetic parameters of peptide hydrolysis catalyzed by NS3/4A. To gain understanding of the reasons for such behavior, we perform atomistic molecular dynamics simulations of NS3/4A in the presence of either PEG or Ficoll crowders and with and without the peptide substrates. We find that both crowder types make nanosecond long contacts with the protease and slow down its diffusion. However, they also affect the enzyme structural dynamics; crowders induce functionally relevant helical structures in the disordered parts of the protease cofactor, NS4A, with the PEG effect being more pronounced. Overall, PEG interactions with NS3/4A are slightly stronger but Ficoll forms more hydrogen bonds with NS3. The crowders also interact with substrates; we find that the substrate diffusion is reduced much more in the presence of PEG than Ficoll. However, contrary to NS3, the substrate interacts more strongly with Ficoll than with PEG crowders, with the substrate diffusion being similar to crowder diffusion. Importantly, crowders affect also the substrate-enzyme interactions. We observe that both PEG and Ficoll enhance the presence of substrates near the active site, especially near catalytic His57 but Ficoll crowders increase substrate binding more than PEG molecules. The presence of crowders also enhances the stability of Zn^2+^ ion coordination necessary for structural stability of NS3/4A enabling catalysis.

**AUTHOR SUMMARY:** Enzyme-catalyzed reactions in reality occur in the crowded environment of the cell. Therefore, viruses entering the host cells also encounter a crowded surrounding in which the viral enzymes are replicated. One such enzyme is the NS3/4A protease encoded by the hepatitis C virus. This enzyme is crucial for viral replication and is used as the therapeutic target for clinically approved drugs. To gain understanding of this enzyme function and explain our previous experiments on its *in vitro* activity, we performed atomistic molecular dynamics simulations in the presence of synthetic crowders (polyethylene glycol and polysucrose) mimicking the cellular crowd. Based on these simulations we describe in detail how and why these crowders affect the diffusion and structural dynamics of this enzyme and enzyme-substrate interactions. In fact, crowders enhance substrate binding, which may have vast consequences for its function in the host cell and drug-design.

## INTRODUCTION

The environment of cells is both crowded and confined by various macromolecules that are physically occupying up to 40% of the cytoplasmic volume [1, 2]. The concentrations of the molecules found in cells, like proteins, lipids, nucleic acids, cofactors, and metabolites, reach up to 400 g/L [3]. This means that all biochemical processes in cells, including enzyme-catalyzed reactions, occur under highly crowded conditions in the presence of various background macromolecules. Importantly, these heterogenous crowders not only reduce the available space, but they also interact with each other. These nonspecific interactions can be attractive or repulsive and may exert either stabilizing or destabilizing effects on the molecules that directly participate in a particular process [4]. Therefore, the crowded surrounding impacts many properties of solutes including diffusion, bimolecular association, folding, flexibility, stability, and ligand-receptor binding [1, 5]. All these may in effect change the catalytic activity of enzymes when compared to their activity in dilute solutions [6]. Typically, enzymes associate with and bind their substrates, and, after the catalysis, the reaction products dissociate. The entire process of diffusion, binding, and product release, which includes conformational changes of the reaction components, is subject to interactions with other macromolecules present in the cell.

For ease of interpreting the results, most laboratory experiments characterizing enzymatic reactions have been performed in dilute buffer solutions. However, to reflect *in vivo* conditions, more and more studies investigate the kinetics of enzymatic catalysis in crowded environments. Concentrated solutions of synthetic polymers are often used as crowding agents [6, 7], to facilitate such experiments and to allow generalizations beyond effects that may be otherwise particular to a specific protein crowder. Common artificial crowders are PEG (polyethylene glycol), Ficoll (highly-branched polysucrose), and dextran (a linear flexible polyglucose). All three types of crowders are available with different molecular weights and polymer lengths to vary crowder sizes. Ficoll, formed by the copolymerization of sucrose and epichlorohydrin, is especially attractive as it is highly soluble, and has a mostly spherical shape. In contrast to protein crowders, artificial crowders are generally considered to be non-interacting with proteins. As a result, such crowders are assumed to focus on the volume-exclusion effect of crowding. However, it is impossible to have a completely inert polymer, and the focus of this work is to better understand how these model crowders may specifically interact with proteins and how such interactions may potentially impact the interpretation of experiments characterizing enzyme function under crowded conditions.

Many experimental assays have shown that kinetic rates estimated in dilute solutions can differ by orders of magnitude from the in-cell values [6, 8]. For example, the PEG crowders were shown to suppress the activity of horseradish peroxidase (ten-fold increase in K_M_) but the change depended on the substrate type [9]. The ATPase hydrolytic activity of the eIF4A translation initiation factor was shown to enhance six-fold in the presence of PEG crowders [10]. PEG, dextran, and Ficoll crowders also enhanced the enzymatic activity of adenylate kinase (AK3L1) [11]. Dextran was shown to induce the activity of the ribonuclease T1 most probably because the crowder promoted the folding of this protein [12]. We have also found that the kinetic parameters of peptide hydrolysis catalyzed by trypsin [13] and HIV-1 protease [14] were affected by PEG and BSA crowders. Our experiments on another protease, NS3/4A, encoded by the hepatitis C virus (HCV), showed different effects of Ficoll and PEG, as well as BSA on the proteolytic reaction [15]. These observed differences are the main motivation for the work presented here. Since enzymes are important drug targets, it becomes crucial to understand how and why the reaction environment modulates the enzymatic rates and equilibria. In general, different factors contribute to the observed changes in crowding-induced catalytic activity such as increased concentration of the substrate due to decreased space, induced folding of the enzyme to its functional form, induced oligomerization of the enzyme to an inactive form, decreased diffusion of the reactive molecules, occupation of the substrate binding or active site by crowders, and the change of the conformation or flexibility of the binding or dimerization site.

Many of the above effects cannot be elucidated using experimental methods alone. Therefore, the impact of crowding on different solute properties and stages of enzymatic reactions has also been investigated using computational approaches [16–19]. However, apart from a few atomistic models [4, 20], most simulations applied simplified crowder models that primarily mimicked only the effect of volume exclusion due to the impenetrability of molecules, e.g., [21–23]. One reason for using such simplified crowder models is computational efficiency, but the choice of such crowders also reflects the assumption that the artificial crowders mainly introduce the volume exclusion effect, and a more detailed representation of molecular-level interactions with crowders may not be needed.

We focus here on the NS3/4A protein complex which is a bifunctional heterodimeric enzyme composed of the serine protease and RNA helicase domains that can function independently [24]. The NS3/4A activity to process the HCV polyprotein is necessary for viral replication. HCV is responsible for a chronic illness affecting nearly 200 million people and this enzyme is the main target in HCV therapy. However, the emerging drug-resistant genotypes and subtypes of this virus necessitate the design of new inhibitors. The NS3 protease domain is a 180-residue protein, with the chymotrypsin-like fold dominated by β-sheets (**Figure 1A**). The active site contains a catalytic Ser139-His57-Asp82 triad and the structure is stabilized by a Zinc ion, which is at the opposite side of the active site. Zinc is coordinated by three cysteines (97, 99 and 145) and His149 (**Figure 1B**) that are conserved in the HCV strains [24] and mutations of any of the cysteines into alanines significantly decreased protease activity [25]. For efficient catalysis, the NS3 protease domain requires a 54 amino acid long peptide co-factor, called NS4A, but only its central part, embedded in the NS3 core, is sufficient for the *in vitro* activity of the protease. The N- and C-terminal parts of NS4A, which are not bound with NS3, are believed to be intrinsically disordered in solution. In the host cell, the NS3/4A complex associates with the endoplasmic reticulum (ER) membrane where the HCV RNA is replicated. The disordered N-terminal tail of the NS4A cofactor can be induced to form a helix that inserts into the ER membrane to anchor the NS3/4A complex [26, 27].

**Figure 1.**
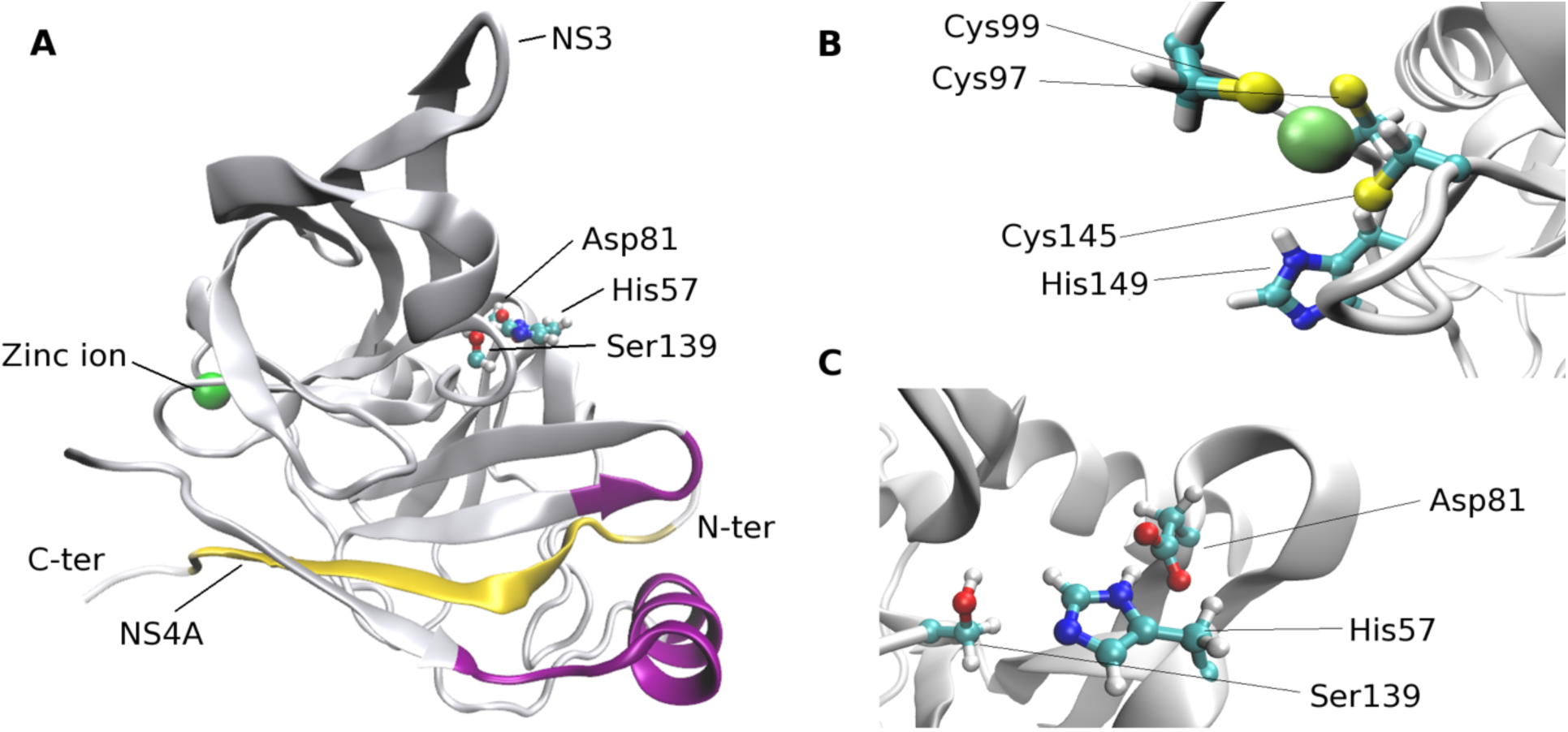
Ribbon model of the NS3/4A protease domain based on the crystal structure with PDB ID: 4JMY [28] (A). The β-sheet of the NS4A cofactor is in yellow and NS3 membrane contact sites are in magenta. The zinc coordinating residues (B) and the catalytic triad (C) are also shown.

Using fluorescence spectroscopy, we have found that PEG 6000 and Ficoll 400 differently affect (in a concentration-dependent manner) the initial and maximum velocities as well as the turnover number for the peptide hydrolysis catalyzed by the NS3/4A protease [15]. The kinetic parameters decreased in the presence of PEG and increased with Ficoll as compared to dilute buffer solutions. Ficoll also increased by 40% the inhibition constant Ki of telaprevir, the clinically-approved drug targeting the NS3/4A enzyme [15]. We have previously explained how PEG crowders may promote folding of the NS3/4A unstructured N-terminal fragment possibly enabling membrane anchoring [27]. However, it remains unclear how the two different crowders may impact enzyme function in an opposite manner. We hypothesize that there are specific interactions between the crowders, NS3/4A, and the substrate that differ between PEG and Ficoll, and the goal of this work is to not just explain the experimental data for NS3/4A but gain a broader understanding of how PEG and Ficoll may interact with proteins at a molecular level.

To probe these questions, we use extensive atomistic molecular dynamics simulations of NS3/4A with and without peptide substrates in the absence or presence of PEG and Ficoll crowders. We analyze how the crowders impact NS3/4A structure and dynamics, diffusion of the enzyme and the substrate, and how the crowders may specifically impact substrate binding near the active site. Based on this analysis we explain how PEG and Ficoll may differently impact the kinetic parameters of the NS3/4A-catalyzed reaction and provide more general insights into how PEG and Ficoll crowders interact with peptides and proteins.

## METHODS

### Preparation of solute structures

As in our previous studies [15, 27], the NS3/4A crystal structure (PDB ID: 4JMY [28]) was used as the starting point for the MD simulations. This structure contains the coordinates of the full protease domain of NS3, residues 21-32 of the NS4A cofactor, the zinc ion critical for maintaining the active structure of the protease [28], and a fragment of the peptide substrate. To match the 1b genotype of HCV, the D30E, L36V, G66A, A87K, M94L, S147F, V150A, I170V, A181S, and S182P mutations were introduced using Chimera [30] with the Dunbrack rotamer library [31]. The missing N- and C-terminal residues of the NS4A cofactor were added with MODELLER [32], ver. 9.19., using a template-free modeling method. Five models of complete NS4A were generated, and, based on the Dope [33] scoring function, the lowest-scoring conformation was selected. Residues were assigned standard protonation states at pH 7, with histidine neutral and protonated at N*δ*1 atom. Zinc-coordinating cysteines were modified with a specific cysteine-zinc patch to reflect deprotonation and polarized charges. The net charge of the NS3/4A-zinc complex was +2e. The peptide substrates with the sequence Ac-DEDEEAASK-NH_2_, corresponding to the sequence of the peptide used in experimental assays [15], were built with CHIMERA [30]. The substrate models were subjected to 2,000 steps of minimization and 20 copies of 200 ns MD simulations in NAMD [34]. Next, the substrate trajectories were clustered with ProDy [35] and the five most probable conformations were used as starting structures surrounding NS3/4A. The NS3/4A protease and substrates were parameterized using the CHARMM36m force field [36] which is suitable for simulating proteins with unstructured fragments.

### Preparation of crowder models

All-atom 28-mer PEG crowder models (**Figure 2**) were prepared as described elsewhere [15, 27]. Starting conformations were taken from MD simulations of a single PEG polymer in explicit water performed by Lee et al. [37]. The CHARMM36 CgenFF force field was used with a modification of the torsion angles based on the work of Leonard *et al.* [38].

**Figure 2.**
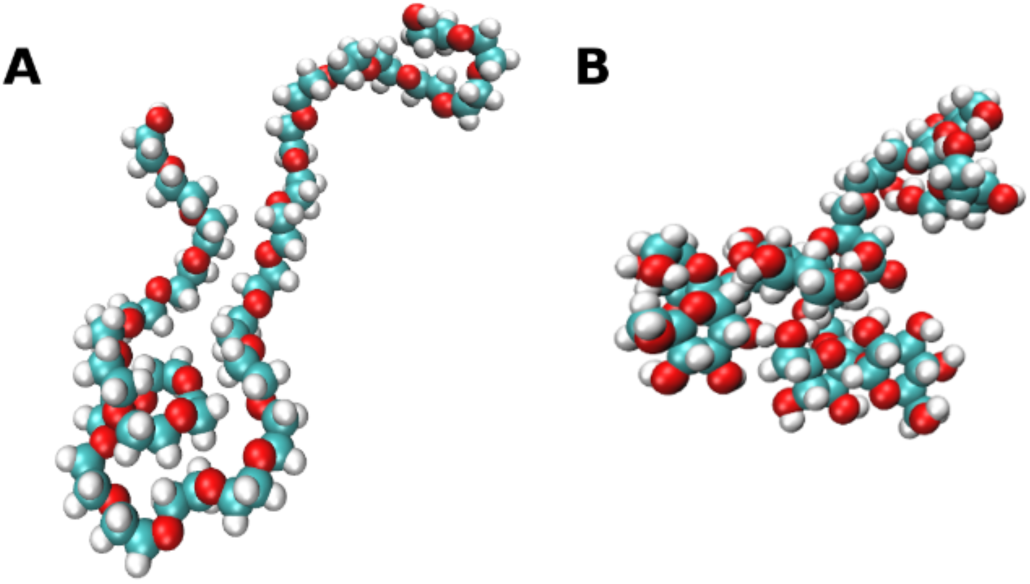
The all-atom model of PEG (A) and Ficoll (B) crowders used in MD simulations.

All-atom Ficoll models were built to match the mass of the PEG crowders (*cf.* **Supplementary Methods** for additional details) so that crowders with similar sizes can be compared. Our Ficoll molecule consists of four sucroses and three glycerol linkers, forming the smallest possible branched Ficoll-like polysucrose (**Figure 2**). Sucrose and glycerol parameters were assigned based on the CHARMM36 force field parameters for carbohydrates [39, 40]. Additional patches were constructed for sucrose-glycerol connections, with parameters derived by analogy to already parameterized polysaccharides in the CHARMM force field, e.g., using parameters for isomaltulose and melezitose (*cf.* **Supplementary Methods**). To obtain the Ficoll starting conformation for the simulations of the crowded systems, one tetra-sucrose was energy-minimized and an MD simulation in explicit TIP3P water was carried out over 200 ns. The starting Ficoll crowder conformations were randomly chosen from the 20% of the MD snapshots with the lowest R_g_ values.

### Simulation systems

Six types of systems were simulated: NS3/4A protease with and without 10 peptide substrates, NS3/4A surrounded by 130 PEG polymers, with and without the substrates, NS3/4A surrounded by 110 Ficoll polymers, with and without the substrates (see **Table S1**). All simulations were performed with explicit TIP3P water molecules and 2 Cl^-^ ions to neutralize the Zn^2+^ ion coordinated by NS3. Na^+^ ions were added to neutralize the simulations with 10 substrates (each substrate has a net charge of -4e). Additional Na^+^ and Cl^-^ ions were added to achieve an ionic strength of ∼20 mM. The MMTSB Toolset [41] was used to solvate the systems and add ions. Initial snapshots of the simulation systems are shown in **Figure 3**.

**Figure 3.**
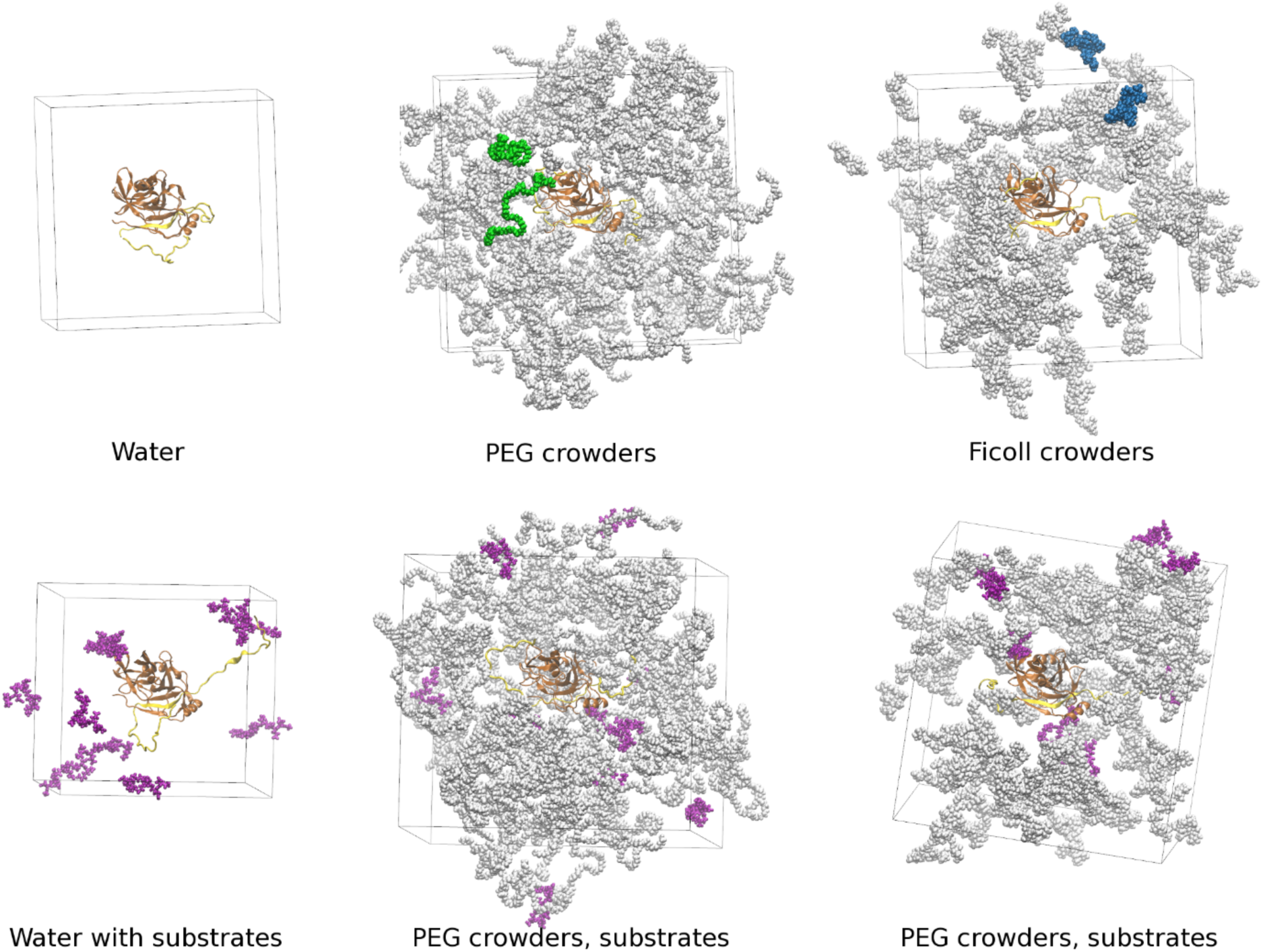
Snapshots of the systems simulated in this study. The NS3/4A enzyme is shown in brown. Substrates are shown in purple. Crowder molecules are shown in grey, but selected molecules are colored to indicate their size (PEG: green, Ficoll: blue). For clarity of the figure, water molecules and ions are not shown

Before solvating the systems, 130 copies of PEG or 110 copies of Ficoll molecules were added in random locations around NS3/4A. Three different sets of crowder positions were generated. The numbers of crowder molecules were chosen to match the mass concentration of crowders; the resulting volume fractions of crowders varied between systems, ranging from 11 to 15%. To avoid clashes, an additional step was carried out for the protease-PEG systems. They were minimized in vacuum before adding water molecules. Considering the high degree of crowding, substrates were added manually using the VMD [42] tools for the systems comprised of both crowders and substrates.

### Molecular dynamics simulations

The simulations were performed for the systems listed in **Table S1**. Each system was first energy-minimized over 3,000 steps using the steepest descent algorithm. Second, water molecules and ions were thermalized by increasing the temperature from 10 to 310 K in steps of 10 K. Each step was simulated for 25 ps, with 10 kcal/mol/Å^2^ harmonic constraints on the positions of the solute atoms. Third, the restraints on the solute were decreased in six 25-ps steps with harmonic force constants gradually decreasing from 10 to 5, 2, 1, 0.1, and 0 kcal/mol/Å^2^. These thermalization and equilibration steps were performed in the NVT ensemble, with a time step of 1 fs. Finally, an additional equilibration step of 25 ps was performed in the NPT ensemble with a time step of 2 fs using the SHAKE algorithm [43] to constrain bonds made by hydrogen atoms. For each system, a 500 ns production simulation was performed either using NAMD 2.11 [34] or OpenMM [44]. A temperature of 310 K and a pressure of 1 atm were maintained with the Langevin thermostat and Langevin piston methods [45]. Two friction coefficients in the Langevin thermostat were used: 1 ps^-1^, the value used in our previous simulations [15, 27], and 0.01 ps^-1^. The lower value gives more accurate estimates of kinetic properties such as diffusion [46, 47]. All trajectories were used to evaluate thermodynamic properties but only the trajectories obtained with a value of 0.01 ps^-1^ were used to analyze diffusive properties and contact life-times. To calculate the electrostatic interactions, the Ewald summation [48] method was used, with a cut-off of 12 Å. To avoid aggregation of the crowders, interactions between the solute and explicit water molecules were increased by a factor of 1.09 because it was found that this modification ensures realistic diffusion of the crowders [27, 47].

Multiple replicates of each system were simulated over at least 500 ns. The NS3/4A enzyme was stable in all simulations as indicated by Cα root-mean square deviation (RMSD) values generally between 1 and 2 Å throughout the trajectories (**Figure S4**). Replicate-averaged RMSD values increased significantly during the first tens of nanoseconds, but they changed slowly afterwards (**Figure S5**). Therefore, we decided to discard the first 50 ns as equilibration from all analyses. To compare the same simulation intervals, all trajectories were analyzed over 50-500 ns, even though some simulations were somewhat longer (**Table S1**).

### Trajectory Analysis

RMSD and RMSF values were calculated for the Cα atoms, after NS3/4A superposition onto the reference crystal structure as a starting point.

Translational diffusion was calculated from the mean-square displacements (*MSD*) of the centers of mass of the molecules. Diffusion coefficients, D_0_, were calculated from the slopes of linear fits to *MSD*(τ) = ⟨(**r**(t+ τ) − **r**(t))^2^⟩ according to the Einstein equation:

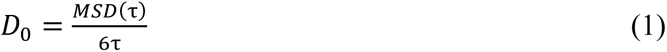

where **r**(t) and **r**(t+τ) are the vector positions of the center of mass of a molecule in time t and t+τ (τ is the time lag). D_0_ values were corrected for the artifacts resulting from using the periodic boundary conditions (PBC) [49] and reduced viscosity of the TIP3P water model (as the ratio of the shear viscosity of TIP3P equal to 0.308 cP and bulk water - 0.89 cP) giving:

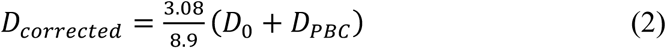

The correction *D_PBC_* was calculated as:

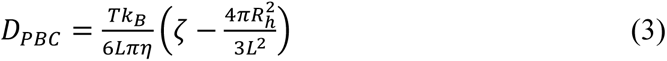

where T is the temperature set to 310 K, *k_B_* is the Boltzmann constant, ς is equal to 2.837, *R_h_* is the hydrodynamic radius of a molecule (calculated with HYDROPRO [50]), and *L* is the length of the simulation box. *η* = *η*_w_(1 + 2.5*φ*) describes the shear viscosity of the solvent as the viscosity of the TIP3P water scaled with a factor depending on φ, i.e., the volume fraction of crowders.

Rotational diffusion of the NS3/4A protease was determined using a method based on the rotational correlation functions [51]. First, a trajectory of random vectors was merged with a trajectory of centered protease. Second, the vectors were rotated along with the protease rotations, obtained by fitting the protease to its reference structure. To calculate the rotational correlation times for the protease, the average correlation function for the vectors was fitted to a double exponential function:

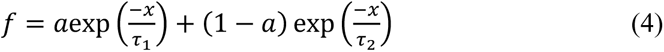

Obtaining the τ_1_ and τ_2_ times describing rotations in longer and shorter time-scale, with *a* being the weight corresponding to the shorter rotational correlation time. Rotational diffusion coefficients were related to calculated correlation times as:

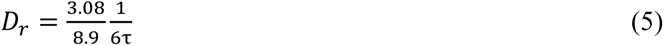

where the 3.08/8.9 is the scaling factor included to account for the reduced viscosity of the TIP3P water model (see above) and the overall t was calculated according to:

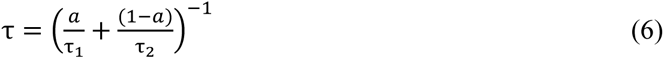

Correlation analysis was used to estimate the time scales of contacts between the molecules and of the dynamics of active site residues. For contact analysis, a contact function was used to describe whether a contact between the molecules present at time t0 is also present at time t_0_+Δt:

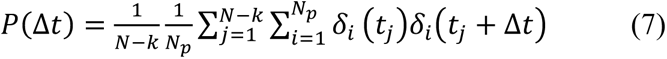

where *N* is the total number of trajectory frames, Δt is the *k*-th time interval and *N_p_* is the number of molecule pairs, and the summation runs over *j* trajectory frames and *i* pairs of molecules. The function *δ_i_(t)* takes the value of 1 when the distance between the *i*-th pair of molecules at time *t* is smaller than 5 Å, and 0 when the molecules are further away from each other. To obtain estimated times of contact survivals, the contact functions were fitted to double exponential functions according to Eq. 4.

The above analyses, along with radii of gyration and radial distribution functions, were conducted using the MMTSB Toolset [41], CHARMM version 46b2 [52], and in-house scripts, in particular to automate efficient data processing. The MMTSB Toolset was also used to cluster substrate binding poses. VMD was used to prepare the structural figures and analyze the hydrogen bonds, with the maximum donor-acceptor distance set to 3.5 Å and the donor-hydrogen-acceptor angle criteria set to 120 degrees. Secondary structures were analyzed using the Stride algorithm [53]. Plots were generated using scripted Gnuplot, version 5.2 [54].

## RESULTS AND DISCUSSION

The analysis of the simulations is organized by first describing the structural stability, conformational dynamics, and diffusive properties of NS3/4A as a function of crowders. We then describe how different crowders interact with NS3/4A and finally how crowders affect the substrates and substrate-enzyme interactions. Finally, we analyze the interaction of water molecules with NS3/4A and the crowders. All of the results are then interpreted in the context of the experimentally observed functional differences of the enzyme in the presence of different crowders.

### Structural stability of NS3/4A in the presence of crowders

The structure of the NS3/4A enzyme remained stable throughout the simulations with average Cα RMSD values of about 1.6 Å for NS3 and the 13 residues of NS4A resolved in the crystal structure (**Figure S4** and **Table S2**). RMSD values appeared to be slightly larger in the presence of Ficoll or PEG crowders compared to water; in the presence of substrates, RMSD values were smaller with Ficoll and larger with PEG. However, in all cases, the differences in RMSD values were on the order of the standard errors and therefore not considered significant.

Average radius of gyration values *R_g_* of 16.2 Å for NS3 (**Table S3**) were slightly larger than the value of 15.8 Å calculated from the experimental coordinates based on 4JMY. Again, there were only slight differences in the absence or presence of crowders, but it appears that the distribution of *R_g_* values is shifted to slightly larger values in the presence of Ficoll (**Figure S6A**) compared to dilute solvent or PEG crowders. However, in the presence of substrates, NS3 is slightly more compact with both PEG and Ficoll crowders (**Figure S6B**).

NS4A has flexible N- and C-termini which result in larger *R_g_* values and larger fluctuations (**Figure S6C**). Average R_g_ values of NS4A appear to be smaller with Ficoll and larger with PEG compared to the system without crowders (**Table S3**); with substrates the opposite trend is found, i.e., NS4A is less compact in the presence of Ficoll and more compact with PEG (**Figure S6D**). However, all of the differences are well within the standard errors and therefore not considered significant.

The presence of crowders is generally believed to favor more compact conformations, especially for more dynamic structural elements. However, for the system studied here, the effect of the crowders on the NS3/4A structure is very modest if any, including for the extended NS4A termini.

### Conformational dynamics of NS3/4A in the presence of crowders

Since NS3 remained overall stable, no major conformational changes were observed. The remaining conformational dynamics characterized via root-mean square fluctuations (RMSF) around the trajectory-averaged structures (**Figure S7**) shows fluctuations for most residues below 1.5 Å, with increased fluctuations of up to about 3 Å for more dynamic regions on the surface of NS3 (**Figure S8**) corresponding to loops and solvent-exposed secondary structure elements. The overall pattern of fluctuations is similar with and without crowders, but the simulation results suggest some differences (**Figure S7**). Without substrates, Ficoll crowders appear to induce greater conformational dynamics at the beginning of the N-terminal helix, around residue 12, at the loop around residue 89, and the β-turn around residue 121, while reducing conformational dynamics in the β-strand linker near residue 73, in residues 95-100, and the β-turn around residue 147. The regions 97-99 and 145-149 include the residues that coordinate the zinc ion (Cys97, Cys99, Cys145, and His149) suggesting that Ficoll may stabilize the zinc binding cavity (**Figure 1B**). PEG crowders affect conformational dynamics less, they also increase fluctuations in the loop around residue 89, but different from Ficoll, they decrease fluctuations in the β-turn around 121 and also slightly in the zinc binding region. This suggests that both crowders perturb the conformational sampling of NS3, Ficoll more than PEG, but with the detailed effects of the two types of crowders being qualitatively different. Previous work has suggested that altered conformational fluctuations in the presence of crowders may correspond to different overall stabilities of the folded state in the presence of the crowders [55]. Some of these effects are retained when substrates are introduced into the systems (**Figure S7B**), but together with substrates, PEG has a greater tendency to lead to increased fluctuations, for example in the entire N-terminal helix (residues 12-23), near 89, and in the β-turn around residue 146, whereas Ficoll with substrates results in more similar fluctuations than without Ficoll. The NS3 N-terminal helix (**Figure 1A**, magenta) is essential for membrane association of the NS3/4A complex [26] and both PEG and Ficoll seem to increase its conformational dynamics.

The cofactor NS4A has flexible N- and C-termini giving rise to greater RMSF values (**Figure S9**). Crowders somewhat reduce the fluctuations, but without substrates the effect is only statistically significant for the N-terminus. When substrates are present, the fluctuations in the NS4A termini are reduced more in the presence of crowders, with no clear differences between PEG and Ficoll (**Figure S9B**).

We previously described the ability of PEG crowders to induce helical structures in the N-terminus of PEG [27]. Such helix formation prepares for membrane anchoring of NS3 and thus we have previously concluded that cellular crowding may assist with this process [27]. Based on the simulations presented here, we again find a greater tendency towards helix formation in the N-terminus of NS4A in the presence of PEG crowders (**Figures S10, Table S4**). Without substrates, Ficoll does not appear to induce helical conformations to the same extent. However, when substrates are present, both PEG and Ficoll induce significantly more helical structures in NS4A compared to dilute solutions, and PEG induces more helical structures than Ficoll (**Figure S11, Table S4**). Similar conclusions are also found when analyzing the φ/ψ peptide backbone torsion angle distributions in the N- and C-termini of NS4A (**Figures S12** and **S13**). This confirms an overall role of crowding in promoting functionally relevant secondary structure formation in NS4A, but also indicates that this observation may depend on the choice of the crowder molecule used to mimic cellular environments.

### Self-diffusion of NS3/4A in the presence of crowders

Translational and rotational diffusion was analyzed from the trajectories with the reduced friction coefficients as described in the Methods section. Translational diffusion coefficients were obtained from linear fits to the mean-square displacement curves shown in **Figure S14** followed by corrections for periodic boundary artifacts and reduced viscosity of the TIP3P water model. The results are given in **Table 1**. As expected, the presence of crowders significantly reduces the diffusion rates of NS3/4A by about one third from diffusion in dilute aqueous solvent. Interestingly, diffusion is reduced more in the presence of PEG than in the presence of Ficoll. When substrate molecules are present, diffusion is reduced further, both without and with crowders and, again, diffusion is slower with PEG crowders than with Ficoll crowders when substrate is present. The substrate molecules therefore act themselves as additional crowders with substrate and crowders apparently affecting translational diffusion of NS3/4A in an independent, additive manner.

**Table 1.** Translational diffusion of NS3/4A, crowders, and substrates Averages based on linear fits to MSD curves in **Figure S14** from 1 to 10 ns over replicate trajectories with a reduced friction coefficient (see **Table S1**) after corrections for periodic boundaries and TIP3P water model. Standard errors of the mean are given in parentheses.

| | Translational diffusion [$\text{\AA}^2/\text{ns}$] | | |
| --- | --- | --- | --- |
|  | NS3/4A | Crowder | Substrate |
| **Water** | 9.40 (0.02) |  |  |
| **PEG** | 6.02 (0.13) | 14.83 (0.09) |  |
| **Ficoll** | 6.75 (0.22) | 20.30 (0.10) |  |
| **Substrate** | 8.02 (0.26) |  | 20.54 (0.39) |
| **PEG/Substrate** | 5.29 (0.39) | 13.92 (0.08) | 13.73 (0.45) |
| **Ficoll/Substrate** | 5.96 (0.02) | 18.69 (0.10) | 16.25 (0.51) |

Rotational diffusion coefficients were obtained from double-exponential fits to rotational correlation functions shown in **Figure S15**. The resulting rotational time scales are given in **Table S5**. Overall rotational diffusion rates, corrected for the faster viscosity of TIP3P water, are reported in **Table 2**. Different from translational diffusion, the presence of crowders alone appears to affect rotational diffusion rates of NS3/4A less. Diffusion actually appears to be slightly higher with PEG, although this result is based on only two simulations (**Table S1**). Moreover, this is a result of slightly faster rotation on fast time scales (3.4 ns with PEG vs. 3.7 ns with water) whereas rotation on slower time scales is retarded with PEG (28.2 ns with PEG vs. 22.7 ns with water) according to the double-exponential fits (**Table S5**). With Ficoll crowders, rotational diffusion of NS3/4A is slowed down more. The presence of substrates reduces rotational diffusion slightly without crowders, but there is a more significant reduction when either PEG or Ficoll crowders are present along with substrates.

**Table 2.** Rotational diffusion of NS3/4A, crowders, and substrates Results from double-exponential fits to combined correlation functions from replicate trajectories with reduced friction coefficient (see **Table S1**), corrected for the TIP3P water model. Standard errors of the mean given in parentheses were estimated from variations between fits to individual correlation functions.

|  | Rotational diffusion [1/ns] |  |  |
| --- | --- | --- | --- |
|  | NS3/4A | Crowder | Substrate |
| **Water** | 0.0170 (0.003) |  |  |
| **PEG** | 0.0179 (0.003) | 0.281 (0.002) |  |
| **Ficoll** | 0.0137 (0.001) | 0.226 (0.001) |  |
| **Substrate** | 0.0164 (0.007) |  | 0.288 (0.011) |
| **PEG/Substrate** | 0.0091 (0.002) | 0.279 (0.002) | 0.256 (0.018) |
| **Ficoll/Substrate** | 0.0112 (0.005) | 0.219 (0.001) | 0.245 (0.086) |

The suggesting that translational and rotational diffusion of NS3/4A may be affected differently in the presence of crowders, led us to further look for evidence of anomalous diffusion, *i.e.*, a variation of diffusion rates over different time scales. This could result, for example from a cage effect where crowders do not hinder short-term motion and rotational motions in place but present molecular obstacles when diffusing over longer distances. Based on plots of log(MSD/t) vs. time (**Figure S16**) we found that the presence of the crowders does indeed give rise to anomalous diffusion since log(MSD/t) decreases significantly from a peak at around 50 ps towards longer time scales. This effect appears to be stronger with PEG than with Ficoll, both without and with substrate (**Figure S16**).

Diffusion of the crowders themselves is also significantly different. Translational diffusion of PEG is significantly slower than that of Ficoll (**Table 1**). This is easily explained by the larger average size of PEG (**Table S3**) compared to Ficoll, but PEG again appears to rotate faster than Ficoll (**Table 2**), potentially indicating differences in crowder-crowder interactions. Finally, substrate diffusion is reduced upon crowding and follows crowder diffusion rates, both for translational and rotational diffusion. For example, substrate diffusion is reduced significantly more in the presence of PEG than in the presence of Ficoll. This suggests that substrates may spend a significant time near crowders and effectively move together. This will be discussed in more detail below.

### Interaction of NS3 with crowders

The PEG and Ficoll crowders are present at high concentrations. Based on radial distribution functions (**Figure 4**), the crowders overall do not have a strong preference for interactions with NS3 since crowder atom densities are higher away from the surface of NS3 than near it. As described previously [27], a peak at 4 Å for NS3-PEG interactions indicates specific interactions when PEG is near NS3. This is largely due to interactions involving PEG’s oxygen atoms (**Figure 4B**). Ficoll mostly lacks distinct features in the RDF, although the number of oxygen atoms (in hydroxyls) that could be involved in interactions with the protein is larger in Ficoll than in PEG (**Figure 4B**). NS3-crowder interactions are changed slightly when substrate molecules are present as well (dashed lines in **Figure 4A/B**).

**Figure 4.**
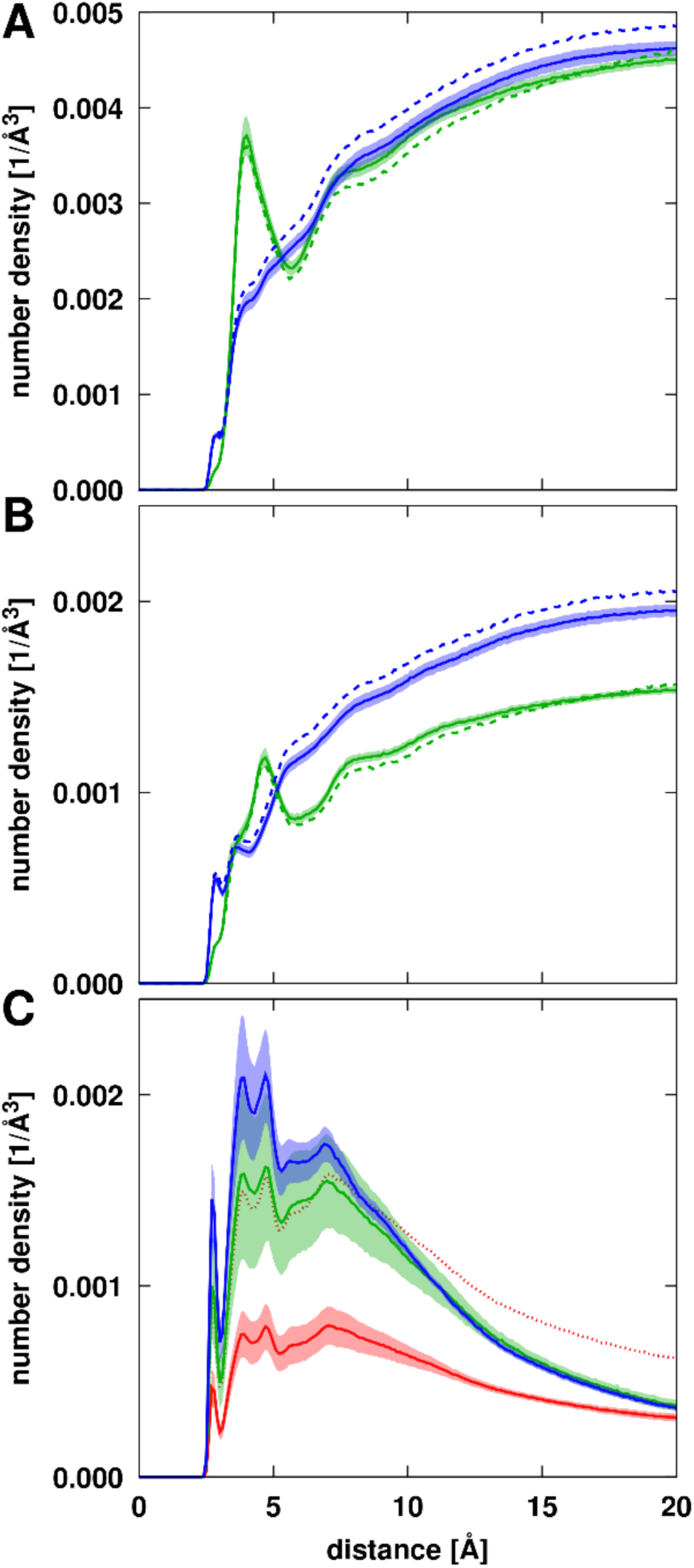
Radial distributions of crowder and substrate atoms around NS3 from simulations with PEG (green) or Ficoll (blue) crowders or in the absence of crowders (red). Distributions are shown for crowder heavy atoms (A), crowder oxygen atoms (B), and for substrate heavy atoms (C). Dashed lines in A and B indicate crowder distributions in the presence of substrates. Distances from the crowder or substrate atoms to the closest NS3 atom were counted and normalized by the total available volume at a given distance from the NS3 surface. For the simulations with only substrates (C), the distribution functions were scaled by a factor of 2 to account for the difference in substrate molarity between the systems without and with crowders (see **Table S1**). The dotted line shows the results obtained without scaling. Distribution functions were averaged over all trajectories for the systems with the crowders but without substrates. The shaded areas indicate standard errors.

To quantify NS3-crowder interactions, we calculated crowder contacts per amino acid residue as well as the average number of crowders in contact with NS3 (**Table 3**). Perhaps surprisingly, there were more than five PEGs and more than three Ficoll molecules in contact with NS3 at any given time. However, this is a consequence of the dense molecular environment with more than 100 crowder molecules (**Table S1**). Interestingly, the number of PEGs slightly decreases and Ficoll increases when substrate is added. Crowder contacts per residue, about 0.2-0.3, tell a similar story (**Table 3**). In addition to simply counting crowder contacts, we also specifically analyzed hydrogen bonding interactions as those are likely to be stronger and potentially longer-lasting contacts. The number of NS3-crowder hydrogen bonds are a small fraction of the total number of contacts, but interestingly, Ficoll formed significantly more hydrogen bonds with NS3 than PEG, even though PEG overall interacted more strongly with NS3 based on the radial distribution functions and total number of contacts. This may be explained by the fact that PEG only has ester oxygens acting as hydrogen bond acceptors, whereas Ficoll has more oxygens and its hydroxyl groups can act both as hydrogen bond acceptors and donors.

**Table 3.** Average number of contacts and hydrogen bonds between NS3, substrates, and crowders Contacts are defined by a minimum distance of less than 5 Å. Standard errors of the mean from variations between replicate trajectories are given in parentheses.

|  | **Contacts /**<br>**NS3 residue** | **H-bonds /**<br>**NS3 residue** | **Interacting**<br>**Molecules/**<br>**NS3** | **Contacts /**<br>**substrate**<br>**residue** | **Interacting**<br>**Molecules/**<br>**Substrate** |
| --- | --- | --- | --- | --- | --- |
| **PEG** | 0.287 (0.015) | 0.021 (0.03) | 5.37 (0.22) |  |  |
| **PEG w/substrate** | 0.279 (0.025) | 0.020 (0.004) | 5.10 (0.33) | 0.298 (0.135) | 0.92 (0.04) |
| **Ficoll** | 0.201 (0.010) | 0.032 (0.005) | 3.49 (0.09) |  |  |
| **Ficoll w/substrate** | 0.213 (0.009) | 0.048 (0.018) | 3.73 (0.06) | 0.949 (0.015) | 1.00 (0.01) |
| **Substrate** | 0.158 (0.026) | 0.046 (0.022) | 2.22 (0.27) |  | 0.13 (0.03) |
| **Substrate w/PEG** | 0.167 (0.045) | 0.050 (0.025) | 2.16 (0.35) |  | 0.10 (0.03) |
| **Substrate w/Ficoll** | 0.215 (0.027) | 0.071 (0.021) | 2.43 (0.14) |  | 0.09 (0.01) |

To further compare NS3-crowder interactions between PEG and Ficoll, we broke down interactions according to NS3 amino acid types (**Figure 5**). Both crowders interacted with all types of amino acids, with about half of the contacts with charged or polar residues. However, there are significant differences between PEG and Ficoll with respect to interactions with acidic residues (Ficoll interacts more strongly) and the classic hydrophobic residues (PEG interacts more strongly). When projecting crowder interactions onto the NS3 surface (**Figure 6**), we find broad coverage across most of the surface, but with preferences for certain parts of the structure. As may be expected, both crowders interacted more with the parts of the NS3 structure that are extruding most from the overall global shape, such as some of the solvent-exposed β-turns. In more detail, there are differences between the parts of NS3 in contact with PEG or Ficoll, presumably reflecting the differences in amino acid preferences. Many of the regions in frequent contact with the crowders correspond to the parts of the NS3 structure, where the presence of the crowders alters conformational fluctuations. This leads to the conclusion that crowder contacts directly modulate NS3 conformational dynamics where frequent contacts are formed.

**Figure 5.**
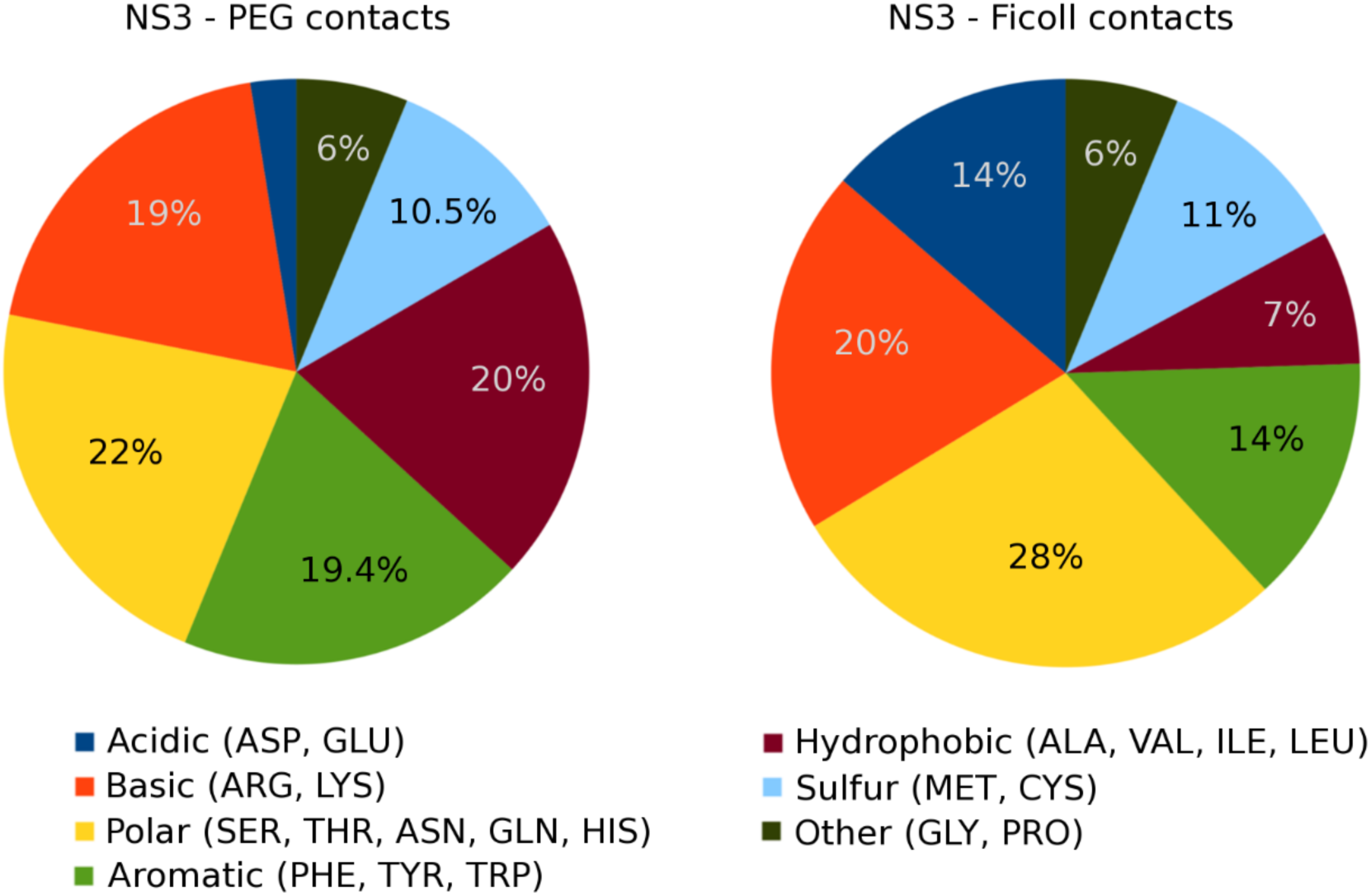
Relative distribution of contacts between PEG and Ficoll crowders and NS3 according to the NS3 amino acids that crowders are in contact with. A contact is defined based on a minimum distance between a crowder oxygen and the closest NS3 heavy atom of less than 5 Å.

**Figure 6.**
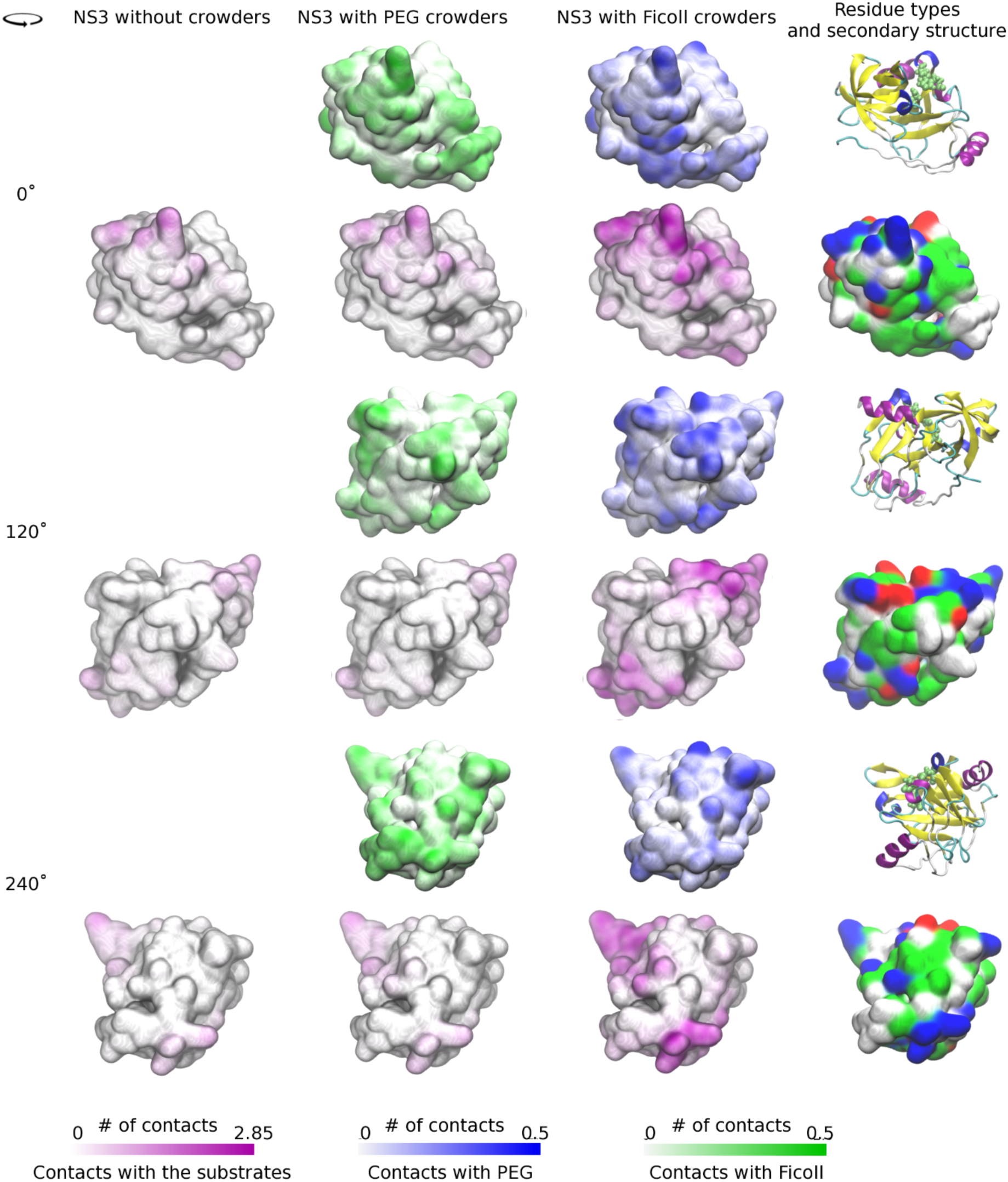
Surface representation of NS3 colored by frequency of crowder and substrate contacts per residue. Different surface colors indicate interactions with PEG (green), Ficoll (blue), and substrates (purple). Different views were generated by rotating the NS3 structure as indicated in the figure. A cartoon representation of NS3 in the same orientations as the surfaces, colored according to secondary structure elements and with van der Waals sphere representation of the catalytic residues, is shown for reference. For clarity, higher number of contacts up to a maximum value of 2.15 contacts per residue are shown at the fully saturated color levels for PEG and Ficoll contacts.

Finally, NS3-crowder contact life times were analyzed based on two-exponential fits to contact survival correlation functions (see Methods). We find that contacts persisted on two life times with roughly equal weight (**Table S6**). Short-lived contacts lasted for about 0.2-0.3 ns, with no statistically significant differences between PEG and Ficoll. Longer contacts lasted for about 6.5 ns with PEG and, significantly shorter, for 4.5 ns, with Ficoll. The observation that NS3 forms frequent contacts with multiple crowders and lasting on time scales relevant for diffusion corresponds to the overall slow-down in diffusion of NS3 in the presence of the crowders described above.

The overall picture from the analysis so far is that crowders do not favor interactions with the enzyme, but interactions lasting as long as several nanoseconds occur nevertheless simply due to the highly concentrated environment. Several crowders are in contact with NS3 at any given time and PEG interactions are slightly stronger, with more specific interactions and longer contact life times compared to Ficoll. However, Ficoll forms more hydrogen bonding interactions with NS3 than PEG and has a stronger preference for acidic residues than PEG. The different amino acid preferences lead to different surface interaction patterns explaining different effects of PEG and Ficoll on the internal dynamics of NS3. Further consequences on substrate binding and enzyme function will be discussed in the following.

### Interaction of substrates with NS3 and crowders

Peptide substrates were present in part of the simulations, without and with crowders. This allows us to analyze substrate dynamics and NS3-substrate interactions in the absence and presence of crowders as well as substrate-crowder interactions. In contrast to the crowders, the substrate peptides interact preferentially with NS3/4A (**Figure 4C**). We note that the concentration of substrates in terms of molarity is about two-fold higher in the control simulations without crowders than in the simulations with crowders (**Table S1**) in order to achieve a comparable number of substrates interacting with NS3/4A with and without crowders. To compare relative preferences of NS3-substrate interactions relative to bulk solvent with and without crowders, the distribution function without crowders was therefore scaled by a factor of two in **Figure 4C**. We find that the presence of crowders increases substrate interactions with NS3/4A and between crowders; Ficoll crowders increase substrate binding more than PEG crowders. On average, there are about two substrates bound to NS3 at any given time (**Table 3**) and substrate contacts per residue are almost the same as crowder contacts per residue. Substrate interactions projected onto the surface of NS3 (**Figure 6**) show strong preferences for certain parts of NS3, mostly near the active site. The substrate is highly acidic and substrate interactions appear to be guided by overall electrostatic attraction to surface patches of NS3/4A with a high concentration of basic amino acids (**Figure 6**). The substrates themselves sample mostly extended conformations with small percentages of helical structure (**Table S4**). Crowders do not significantly change the secondary structure sampling of the substrates (**Tables S4** and **Figure S17**). However, in the presence of the crowders, the substrates appear to be slightly more compact as average radii of gyration are reduced by 1-2% (**Table S3**).

Substrate contacts with crowders are characterized in **Figure 7**. As for NS3, the peptide substrates do not interact preferentially with the crowders. In contrast to NS3, the substrate interacts more strongly with Ficoll than with PEG. This is a result of Ficoll preferring interactions with acidic amino acids (**Figure 7B**) relative to PEG. As a result, there is, on average, almost 1 contact with Ficoll per substrate residue, but only 0.3 contacts per residue with PEG (**Table 3**). Nevertheless, the number of crowder molecules in contact anywhere with a given substrate is similar, around one crowder per substrate on average (**Table 3**). Substrate-crowder life times are somewhat shorter than NS3-crowder life times, with short contact times of 0.1 ns and long contact times of around 1.5 ns (**Table S7**). This suggests a picture where substrates mostly travel attached to a crowder. Substrates detach from crowders and exchange interactions frequently, but the contacts still remain long enough for substrate diffusion to be dominated by crowder diffusion as discussed above.

**Figure 7.**
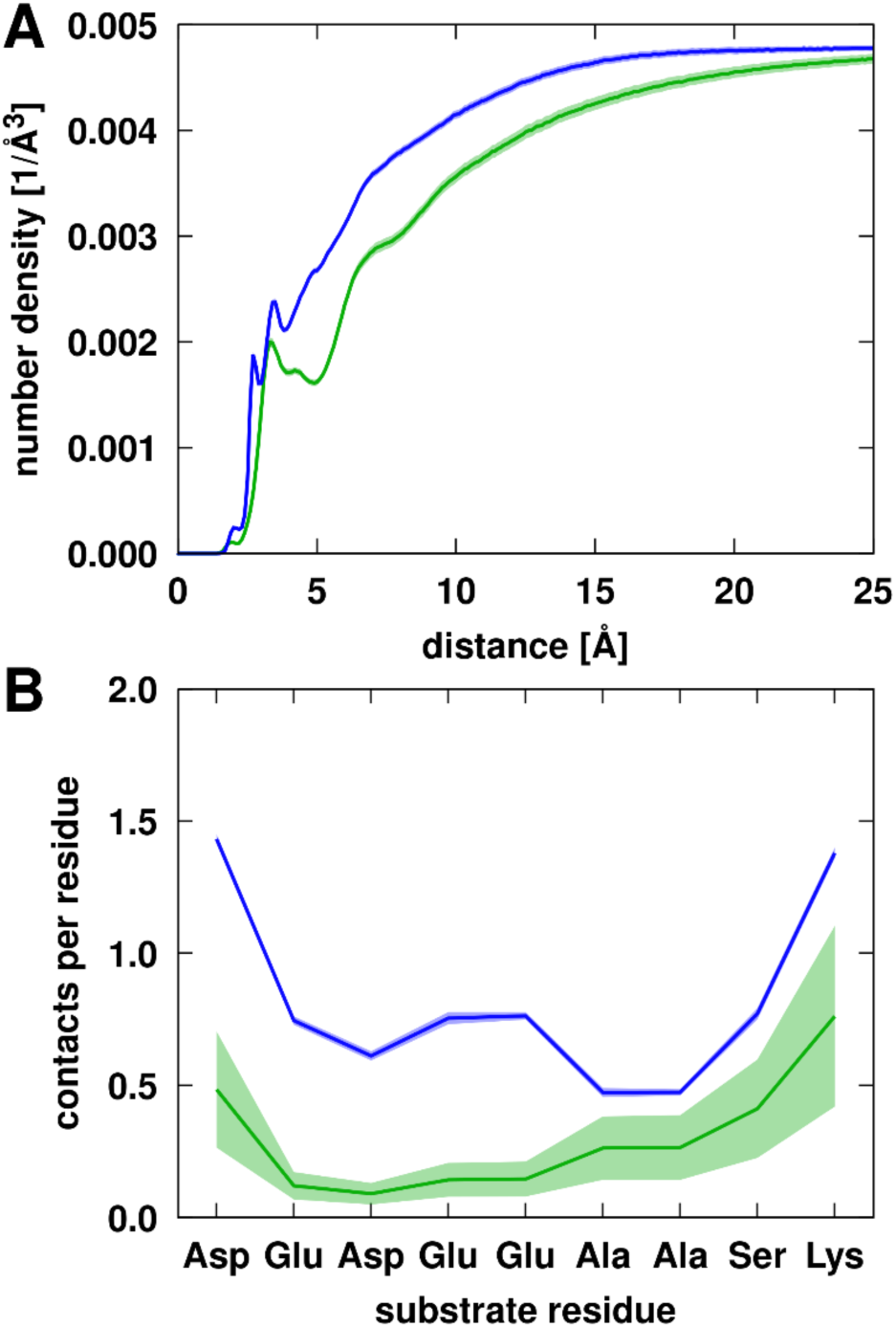
(A) Radial distribution of PEG (green) or Ficoll (blue) crowder heavy atoms around substrates. Distances from the crowder atoms to the closest substrate atom were counted and normalized by the total available volume at a given distance from the NS3 surface. (B) Contacts with PEG (green) or Ficoll (blue) per substrate residue along the substrate sequence. Trajectory averages are shown as solid lines with the shaded areas indicating standard errors.

### Crowding effects on NS3 active site

We now focus on the specific effects near the NS3 active site with the ultimate goal of explaining the experimental data that showed enhancement of NS3/4A activity in the presence of Ficoll crowders but reduced activity with PEG crowders [15]. According to radial distribution functions, the crowders themselves do not interact strongly with the active site residues, although Ficoll interacts more than PEG (**Figure 8**). At 10 Å distance, crowder atom densities range from less than 0.0005 Å^-3^ near Ser139 to 0.0012 Å^-3^ near His57 which is much less than the average crowder density of 0.0035 Å^-3^ with respect to any residue of NS3 (**Figure 4**). This may be expected since the active site residues lie at the bottom of a surface pocket (**Figure 1A/C**) where non-specific binding with the crowders is less likely.

**Figure 8.**
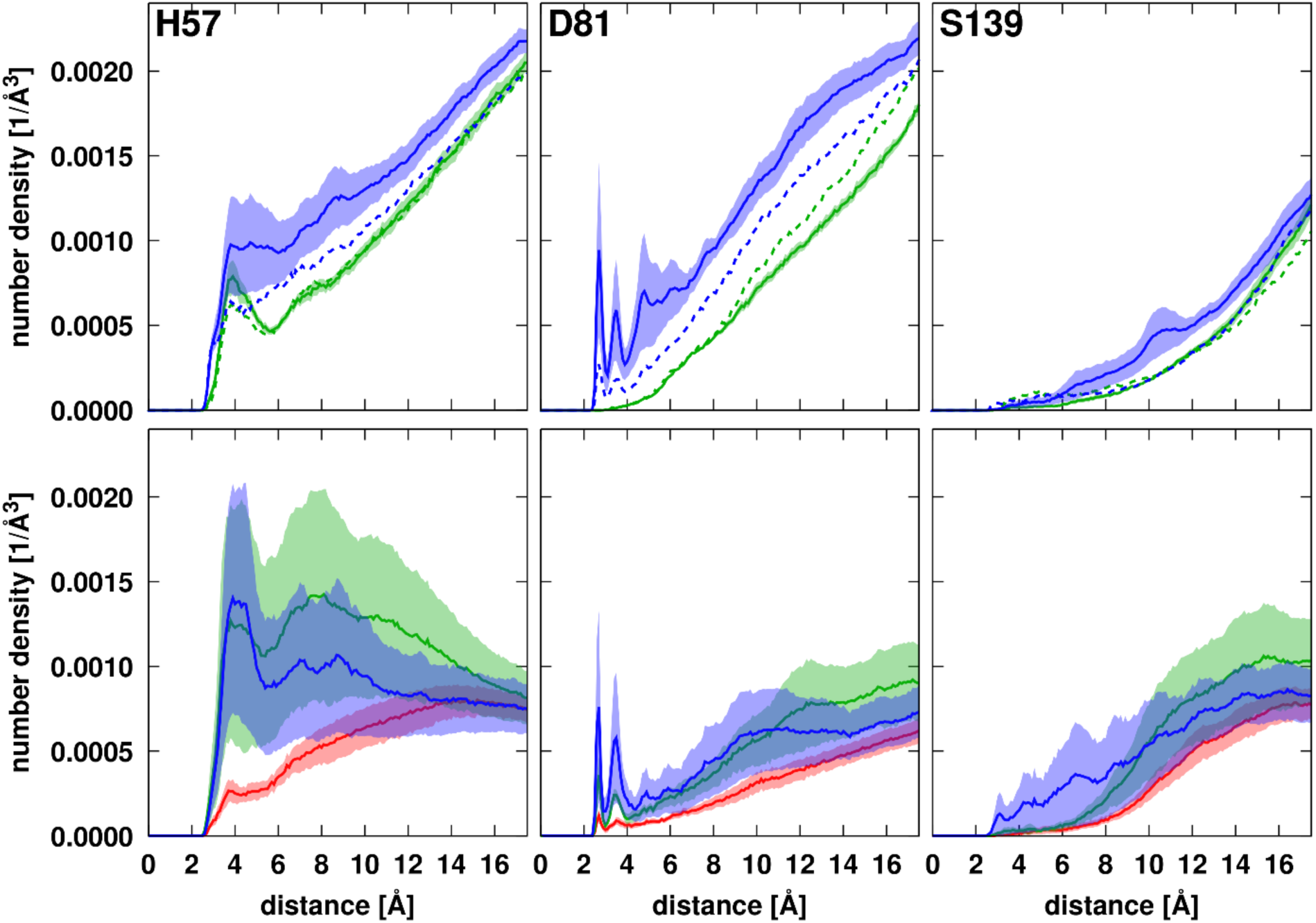
Radial distribution of PEG (green) or Ficoll (blue) crowder heavy atoms (top row) or substrate heavy atoms (bottom row) in simulations with PEG (green), Ficoll (blue) or without crowders (red) around NS3 active site residues. Distances from the crowder atoms to the closest substrate atom were counted and normalized by the total available volume at a given distance from the given NS3 residues. The distribution function of substrates in the simulations without crowders are not scaled; note that substrates are present at twice the overall concentration in those systems. Dashed lines in the top row panels indicate crowder distributions in simulations with substrate. Trajectory averages are shown as lines with the shaded areas indicating standard errors.

Substrate molecules are expected to bind near the active site for cleavage, but, without crowders, substrate atom densities near the active site are similar to crowder atom densities (**Figure 8**). However, both crowders significantly enhance the presence of substrates near the active site, especially near His57. At the same time, Ficoll crowder densities near the active site are reduced when substrate is present.

The geometry of the active site residues is important for enzyme function. Asp81 is expected to coordinate with His57, and His57 would attach the hydroxyl of Ser139 in the presence of substrate in the first step of the peptide cleavage reaction [56]. Therefore, we analyzed whether the presence of a crowder and/or a substrate at close distance to the active site would affect the conformational sampling and interactions between the active site residues.

We used RMSD of the active site residues (57, 81, 139) to compare the overall geometry of the active site relative to the crystal structure during the simulations (**Figure 9A/B**). As the structure of the enzyme fluctuates, there is a distribution of conformations ranging from structures very similar to the crystal structure (<1 Å) that are likely reflecting a conformation close to the catalytically competent state to structures that deviate more significantly (up to about 3 Å). The most likely structures seen in the simulations deviate by about 2.2 Å. When crowders are present, close contacts of PEG increase the sampling of crystal-like structures, whereas close Ficoll contacts actually decrease the sampling of such conformations (**Figure 9A**). When substrates are present as well, neither crowder enhances the sampling of crystal-like conformations when coming in contact with the active site (**Figure 9A**). The functionally more relevant scenario involves cases where substrate molecules approach the active site. Without crowders, crystal-like structures are not sampled as frequently as in pure water when substrate molecules come close to the active site (**Figure 9B**). The additional presence of PEG does not change that. However, the addition of Ficoll restores the more frequent sampling of crystal-like active site conformations when substrate approaches the active site seen in pure water (**Figure 9B**). This suggests that PEG and Ficoll crowders act differently in promoting catalytically relevant active site geometries along with substrate binding.

**Figure 9.**
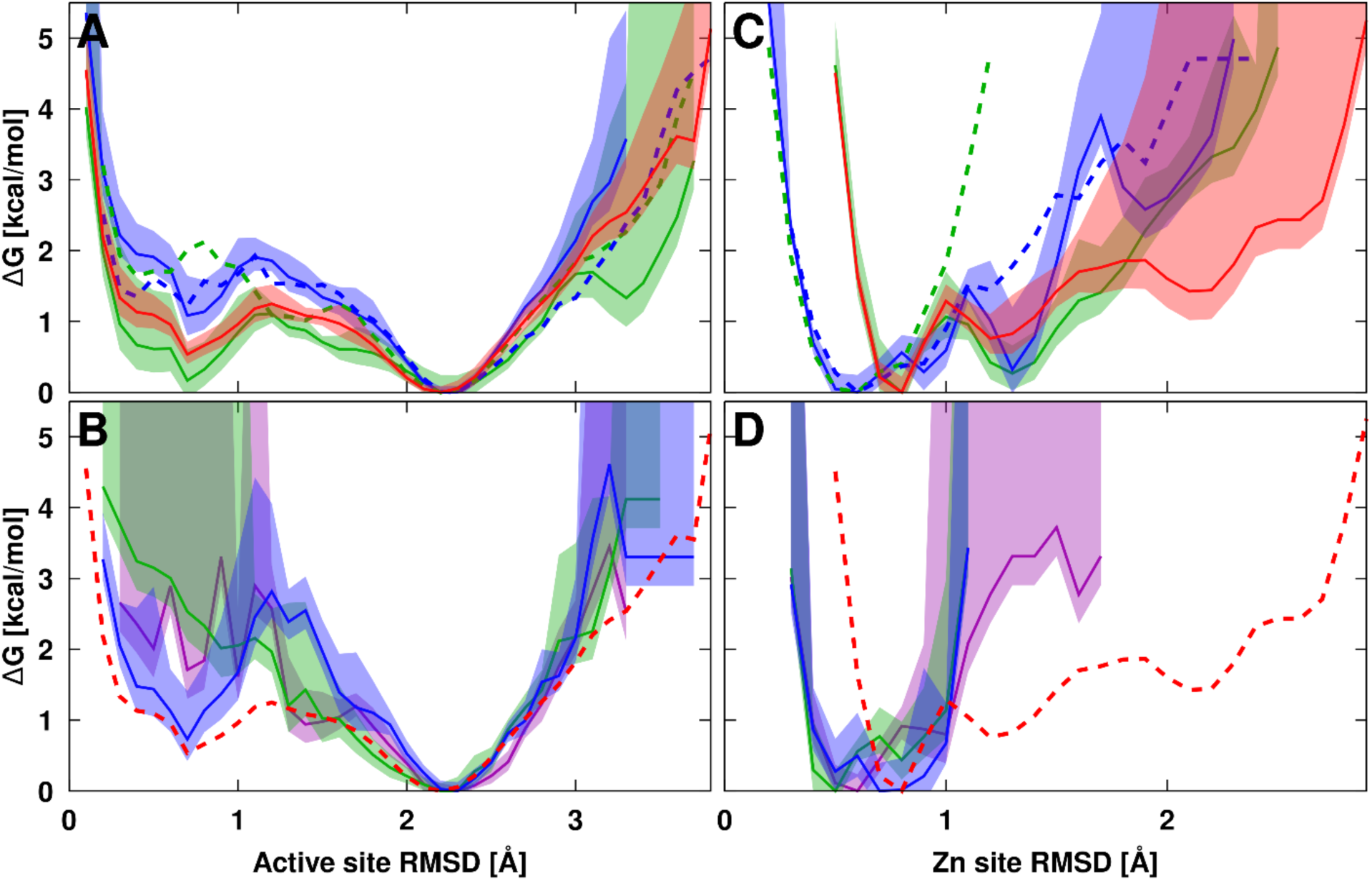
Relative free energies based on probability of sampling different values of root mean-square deviations (RMSD) of heavy atoms of active site residues (57, 81, 139) (panels A and B) or Zn_2+_-coordinating cysteines (97, 99, 145) (panels C and D). RMSD values were calculated with respect to experimental reference structure (PDB ID: 4JMY) after optimal superposition of the respective residues. Free energies were shifted so that the most likely distance corresponds to an energy of 0 kcal/mol. Results are shown for simulations in water (red), with only substrates (purple), and in the presence of PEG (green) or Ficoll (blue) crowders. The top panels (A, C) compare distributions in water with distributions when crowders are in contact (<5 Å). Solid lines show results without substrate, dashed lines show results when substrates are present along with the crowders. In the bottom panels (B, D), distributions are shown for cases where the substrate is in contact (<3 Å). For comparison, the dashed line shows the distribution in water. Trajectory averages are shown as lines with the shaded areas indicating standard errors.

To analyze the effect of crowders and substrate on the NS3 active site in more detail, we calculated the distribution of side chain χ_1_ torsion angles and inter-residue distances between active site residues. χ_1_ torsion angles as a function of the closest crowder or substrate distance from the active site are shown in **Figures S18-S20**. In addition, we compare the distribution of χ_1_ angles between dilute solvent without crowders or substrates with the distributions when a PEG, Ficoll, or substrate molecule is within 5 Å of any of the active site residues (**Figure S21**). The sampling of side chain orientations is broad, covering different rotameric states and not just the conformation found in the crystal structure (PDB ID: 4JMY) (**Figure S21**). This may be expected since the residues are solvent-exposed and the crystallographic conformation may be further constrained by the presence of a tightly bound inhibitory peptide. When crowders interact without substrates, the side chain orientations appear to change somewhat, for example the rotameric state with χ_1_=60° for His57 seen also in the crystal structure became less populated when Ficoll crowders interacted but more populated when PEG crowders were close. However, most of these effects disappear when substrates are present along with the crowders (**Figure S21**). On the other hand, close substrate contacts themselves also appear to modulate the torsional sampling, especially when a substrate is in very close direct contact (<3 Å). More specifically, χ_1_=60° for His57 and χ_1_=-150° for Asp81 become the preferred states similar to the crystal structures when substrate is in very close contact in the presence of Ficoll (**Figure S20**).

We further analyzed side chain center distances between active site residues. As the side chain orientations fluctuated, side chain distances also displayed significant dynamics (**Figure S22-S24**). Close crowder contacts without substrates changed the distribution of distances (**Figure S25**). Closer distances between all active site residues were favored when a PEG crowder was nearby. In contrast, Ficoll near the active site led to the opposite effect of active site residues moving further away from each other. However, as for the torsion angle sample, these effects mostly disappeared when substrates were present along with the crowders (**Figure S25**). Substrate contacts themselves had overall a more moderate effect, but with PEG crowders, side chains came moderately closer when substrates were close to the active site (**Figure S25**). With Ficoll, the effect was again more focused on very close substrate distances to the active site that promoted closer side chain interactions (**Figure S24**). This suggests that crowder-induced close substrate interactions may facilitate an induced-fit mechanism of substrate binding in the active site, especially with Ficoll crowders.

We proceeded to look at two key distances, His57:Nδ-H … Oδ1/2-Asp81 and His57-Nε … HOγ-Ser139, in more detail. These distances need to be close for the cleavage reaction to proceed. Crowder contacts alter the distance distributions in a similar fashion as the side chain distances (**Figure S29**), with most effects again disappearing when substrate is present. Substrate contacts by themselves without crowders do not strongly promote closer distances (**Figure S29**), but close His47-Ser139 hydrogen bond distances necessary to start the cleavage reaction become relatively more favorable than intermediate distances of 4-6 Å when Ficoll crowders are present (**Figures S28** and **S29**). In fact, with PEG crowders and substrate, sampling of close distances (<2.5 Å) of this hydrogen bond almost vanishes (**Figure S28**), whereas Ficoll enhances this interaction when a substrate is close to the active site.

To complement the conformational sampling analysis of active site geometries, we characterized side chain fluctuations from orientational correlation function decays (**Figure S30**). Double exponential fits to the correlation functions (**Table S8**) reveal fast motions on time scales of around 1 ns and slow motions on time scales between about 50 and 500 ns. The slower time scales presumably correspond to transitions between different rotamer states. We find that the presence of Ficoll crowders generally accelerated the slow motions over dilute solvent dynamics when substrates are not present, whereas PEG crowders slowed down those motions for His57 and Ser139 but not for Asp81. The presence of substrates by themselves, accelerated the side chain motions significantly, about twofold (**Table S8)**. With crowders and substrates, the effect on side chain dynamics became more complicated; His57 dynamics remained fast with PEG and it became even faster with Ficoll. Asp81 dynamics slowed down significantly with PEG, whereas Ser139 dynamics slowed down significantly with Ficoll. This may suggest that the crowders can influence the dynamics of functionally relevant residues to different degrees with possible consequences for enzyme catalysis.

### Crowding effects on substrate binding modes

Finally, we looked at the binding modes of substrates near the active site. Experimental structures of NS3/4A provide partial evidence of where the substrate is expected to bind NS3, mainly based on the binding mode of inhibitory peptides [57, 58]. According to these studies, the substrate binds in a mostly extended conformation with the N-terminal part of the substrate interacting with Asp168 of NS3 and pointing towards the NS3/4A active site (comprised of His57, Asp81, and Ser139). We analyzed the preferred binding modes from the simulations based on clustering analysis of substrates found near the active site (**Figure 10**). Substrates interacted in a variety of ways. Only some conformations were close to the exact conformation of the inhibitory peptide bound in the crystal structure (PDB ID: 4JMY). Without crowders, we find that bound substrates were generally located in the surface depressions above and to the left of the active site in **Figure 10B**, interacting with key residues known from previous studies to be involved in substrate binding (such as Asp168 at the N-terminal side of the substrate and Gln41, Ser42, Arg109, and Lys136 for the C-terminal part of the substrate near the cleavage site [58, 59]). However, some of the bound conformations appeared to be relatively far from the surface (**Figure 10F**). In the presence of Ficoll crowders, substrate binding appeared to become more focused towards the inhibitory peptide binding site in the crystal structure, and mostly involved linearly extended substrates across the active site (**Figure 10C/G**). In contrast, the presence of PEG appeared to have an opposite effect of more diffuse binding of the substrate pointing in different directions but without clear examples of the substrate following the binding mode along the cleft above the active site suggested by the inhibitory peptide bound in the crystal structure.

**Figure 10.**
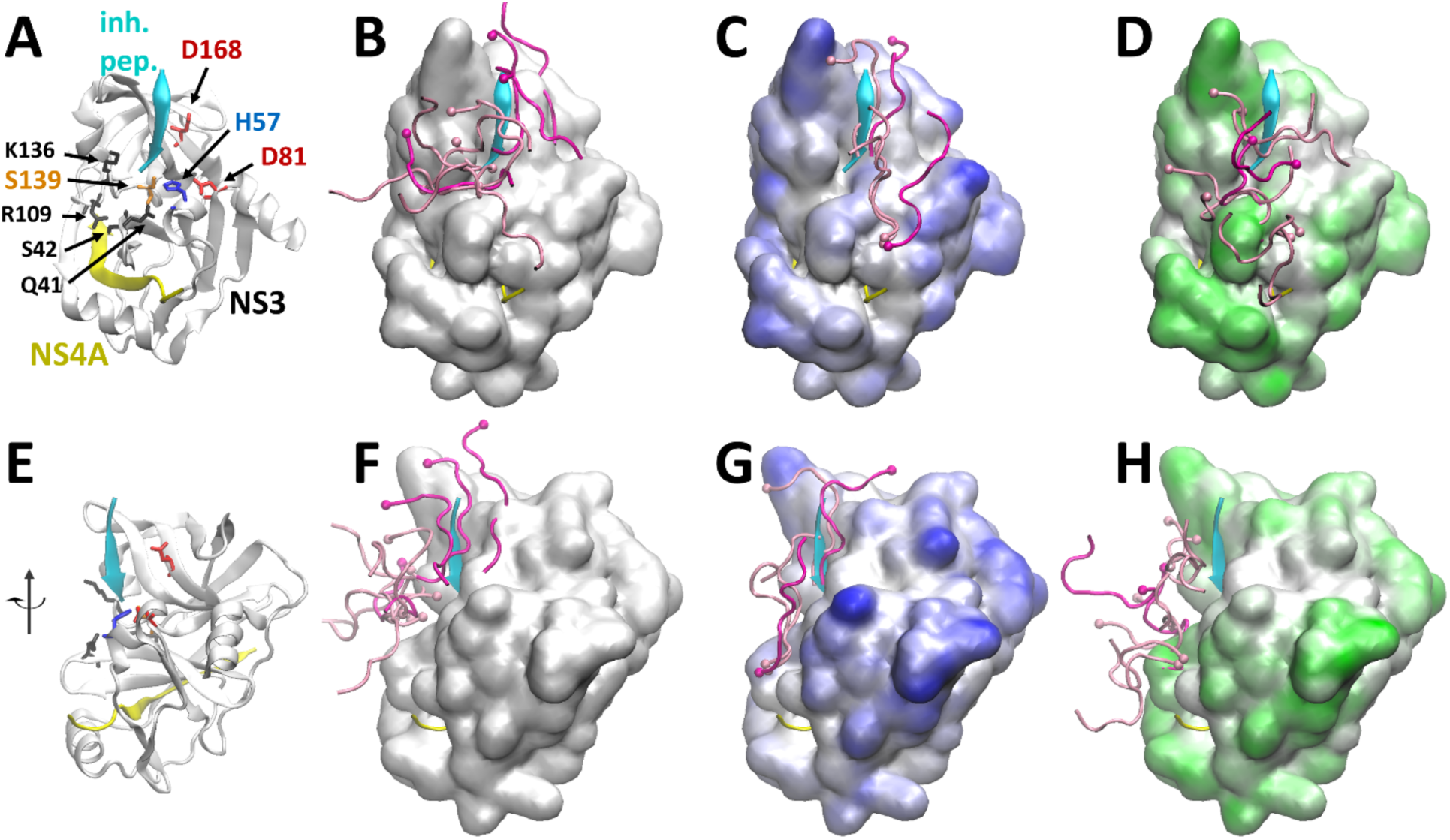
Substrate binding near NS3 active site from simulations without crowders (B/F), with Ficoll crowders (C/G) and with PEG crowders (D/H). Substrate conformations shown are representative structures closest to cluster centers for the largest clusters covering the top 25% (dark magenta) and the next 25% (light magenta) of all substrate conformations extracted for a given system. Surface representations of NS3 are colored based on crowder contact frequencies, with Ficoll (C/G, blue) and with PEG (D/H, green). A cartoon representation of NS3/4A is shown for reference with labels identifying key residues in the active site and those involved in substrate binding (A/E). The bottom row of figures shows the protein after rotation to the left by about 45°.

Along with the differences in substrate binding modes, **Figure 9** also shows the preferences of crowder molecules to interact with NS3 residues. It can be seen that Ficoll interacts more prominently on the right side of the active site (near D81 and above), whereas PEG crowders interact much less in that area, presumably because of Ficoll’s preference for acidic residues. Based on this analysis, we speculate that Ficoll binding to NS3 guides substrates to more easily find the catalytically relevant binding mode whereas PEG binding to NS3 may actually have the opposite effect of discouraging catalytically competent substrate binding.

### Crowders and substrates effects on zinc binding site

In addition to the active site, the coordination of a Zn^2+^ ion by three conserved cysteines (97, 99, and 145) is also essential for NS3 function. Since the Zn^2+^-binding site is opposite to the active site (**Figure 1A**), it is not expected to be directly involved in catalysis. Instead, this motif is presumed to stabilize the NS3 structure [24]. To examine whether the crowders had an impact on Zn^2+^-coordination, we carried out a similar analysis as for crowding effects near the active site. An analysis of the Zn^2+^-coordinating cysteine conformations based on RMSD to the experimental reference structure (PDB ID: 4JMY) shows that the conformations sampled in the trajectories generally remain close to the crystal structure, indicating that Zn^2+^-coordination is maintained in all cases (**Figure 9C/D** and **Figures S31/S32**). The preferred conformation in water is about 0.8 Å RMSD away from the crystal structure (**Figure 9C**). In this conformation all cysteine centers of mass are about 3 Å from the Zn^2+^ ion. In the simulations with crowders, both PEG and Ficoll interact with NS3 near the Zn^2+^ motif (**Figure 6**). When crowders without substrates come into close contact to the Zn^2+^-binding site, the distributions remain mostly the same with PEG (**Figure 9C**), but in the presence of Ficoll significantly shifts the conformations to structures closer to the crystal structure (**Figure 9C**). In this conformation, Zn^2+^-Cys97 distances remain at 3.5 Å, but shorter distances of 2-2.5 Å are preferred for Zn^2+^-Cys99 and Zn^2+^-Cys145 distances (**Figure S31**).

At the same time, cysteine side chain torsion angles also more likely assume the conformations seen in the crystal structure (**Figure S32**). This results effectively in a tighter coordination of Zn^2+^. When substrates are present, close crowder contacts induce the tighter Zn^2+^-coordination with both types of crowders (**Figure 9C**). This suggests that crowders, especially Ficoll, may indirectly support NS3 function by stabilizing the Zn^2+^ motif that is essential for structural integrity of NS3. Substrate molecules also induce a shift to tighter Zn^2+^-coordination, with or without crowders (**Figure 9D** and **Figure S31**). The acidic substrate molecules were found to bind near the Zn^2+^ motif in addition to the active site environment (**Figure 6**), presumably attracted by the Zn^2+^ ion as well as basic residues nearby. The analysis here suggests that substrate binding by itself may also help with stabilizing the Zn^2+^ motif in addition to the crowder effects.

## CONCLUSIONS

Understanding enzyme function in the presence of crowders is important for understanding enzyme function *in vivo*. Proteins are the main type of crowder molecules encountered in biology but experiments often use artificial crowders to focus on generic crowder properties and avoid the complexity of biological environments. The present study was motivated by the observation that the enzyme kinetics of NS3/4A responded very differently to two different types of artificial crowders, PEG and Ficoll. Although K_M_ values in the absence and presence of crowders were found to be similar, catalytic rates decreased with PEG and significantly increased with Ficoll compared to the enzyme in buffer solution [15]. This suggested that these commonly used crowders may affect enzyme function via crowder-specific interactions rather than the more generic volume exclusion effect typically assumed to be dominant for these types of crowders [1]. The extensive simulations of NS3/4A in the presence of PEG and Ficoll as well as peptide substrates that are presented here offer molecular-level insights into how PEG and Ficoll may interact differently with NS3 and the substrates, and how they may lead to different enzyme behavior. As expected, neither PEG and Ficoll interact strongly with NS3. However, interactions do occur with enough frequency for NS3 to be in contact with several crowder molecules at any given time. Moreover, NS3-crowder interactions are relatively long-lived, up to several nanoseconds. An interesting finding is that PEG and Ficoll have different preferences for different amino acid types. Ficoll clearly prefers interactions with acidic residues, whereas PEG is more likely to interact with amino acids that have hydrophobic aliphatic side chains. As a result, crowders decorate different parts of the NS3 surface according to their amino acid preferences.

The crowders also interact with the substrate peptides, especially Ficoll because the substrates are highly acidic. In fact, based on the analysis presented here, substrates rarely diffuse on their own but are in contact with a crowder at any given time. This suggests that substrate diffusion is essentially slaved to crowder diffusion. The crowders considered here were relatively small, and therefore the effective substrate diffusion was reduced more moderately in the presence of crowders, but one expects this effect to be significantly more pronounced when much larger crowders such as PEG 6000 or Ficoll 400 are used as common in experiments. In that case, reduced substrate diffusion could become a rate-limiting step for enzyme reactions. On the other hand, we found that the presence of crowders enhanced substrate binding to NS3, perhaps by delivering substrates to NS3 and then hindering substrates from leaving once bound to NS3. Moreover, crowders appeared to modulate the substrate binding modes, essentially by binding to certain NS3 surface regions thereby focusing substrate binding on the remaining patches that are unoccupied by crowders. With PEG crowders this led to more diffuse binding, while Ficoll may focus substrate binding towards a catalytically competent pose. In addition, Ficoll crowders also appeared to facilitate the formation of catalysis-relevant conformations of active site residue when a substrate molecule was closely interacting. Moreover, Ficoll crowders were found to stabilize the Zn^2+^-binding motif essential for structural integrity of NS3. PEG crowders only showed this effect together with substrate molecules. Taken together, this may explain the experimental observation of increased NS3/4A activity in the presence of Ficoll compared to buffer and when PEG is present.

While the study highlights specific effects of PEG and Ficoll due to different interactions with proteins and peptides at the molecular level, it does not fully interpret the experimental data. Substrate binding near the active site was observed, but it is unclear whether any of the bound substrates fully reached a catalytically competent binding pose. Probably much more extensive simulations with enhanced sampling techniques could overcome this challenge. However, it is also not clear from experiment how exactly cleavable substrates would bind to the active enzyme. Another limitation is that the crowder models used here may compare to PEG 600 or 2000, but not the much larger PEG 6000 and Ficoll 400 crowders used in the experiments. Simulations with much larger crowders would be computationally prohibitive. Another complication is that the higher-order structure of Ficoll is not well-characterized. Finally, the simulations presented here only address classical aspects of biomolecular interactions and dynamics but do not directly describe the peptide cleavage reaction, which requires quantum-mechanical treatments [56].

While the work here focuses on the specifics of the NS3/4A protease and compares the effects of two types of artificial crowders, the more general conclusions are that even artificial crowders may significantly impact functional studies under crowding conditions as a result of molecule-specific interactions with proteins. We hope that the insights gained here will motivate further studies of enzymes under crowded conditions and guide the interpretation of results obtained with different crowding agents.

## ACKNOWLEDGMENTS

Funding was provided by the University of Warsaw “Excellence Initiative – Research University (2020 – 2026)” Tandems for Excellence project and by the National Institute of Health (NIGMS) grant R35 GM126948 to MF.

## AUTHOR CONTRIBUTIONS

**Natalia Ostrowska,** Conceptualization, Data Curation, Formal Analysis, Investigation, Methodology, Software, Validation, Visualization, Writing – Original Draft Preparation; **Michael Feig,** Conceptualization, Data Curation, Formal Analysis, Investigation, Methodology, Software, Validation, Visualization, Writing – Original Draft Preparation, Writing – Review & Editing; **Joanna Trylska,** Conceptualization, Funding Acquisition, Investigation, Project Administration, Resources, Supervision, Validation, Visualization, Writing – Original Draft Preparation, Writing – Review & Editing

## SUPPORTING INFORMATION

### S1 Text. Supplementary Methods

**Figure S1.** Chemical structure of the polysucrose molecule used as a model for Ficoll.

**Figure S2.** Building blocks used to parameterize Ficoll molecules.

**Figure S3.** Structures of isomaltulose and melezitose and CHARMM force field patches.

**Figure S4.** Time evolution of coordinate root mean square deviations of NS3/4A.

**Figure S5.** Time-averaged Cα root mean square deviations as a function of simulation time.

**Figure S6.** Radius of gyration distributions of NS3 and NS4A.

**Figure S7.** Root mean square fluctuations of NS3.

**Figure S8.** Root mean square fluctuations of NS3 in non-crowded and crowded environments projected onto the protease structure.

**Figure S9.** Root mean square fluctuations of NS4A.

**Figure S10.** Secondary structure evolution of the NS4A cofactor in the simulations with water and with either PEG or Ficoll crowders.

**Figure S11.** Secondary structure evolution of the NS4A cofactor in the simulations with substrates and with either PEG or Ficoll crowders.

**Figure S12.** Ramachandran map of backbone torsion angles φ and ψ for N-terminal residues (residues 2-19) of NS4A.

**Figure S13.** Ramachandran map of backbone torsion angles φ and ψ for C-terminal residues (residues 31-53) of NS4A.

**Figure S14.** Mean square displacement of centers of mass vs. time used for determining translation diffusion coefficients for NS3/4A.

**Figure S15.** Rotational correlation functions used for determining rotational diffusion coefficients for NS3/4A.

**Figure S16.** MSD/t vs. time on a log-log scale based on trajectory averaged mean square displacement curves.

**Figure S17.** Ramachandran map of backbone torsion angles φ and ψ for substrates.

**Figure S18.** Side chain χ_1_ torsion angle sampling of active site residues as a function of closest PEG crowder distance.

**Figure S19.** Side chain χ_1_ torsion angle sampling of active site residues as a function of Ficoll crowder distances.

**Figure S20.** Side chain χ_1_ torsion angle sampling of active site residues as a function of substrate distances.

**Figure S21.** Sampling of side chain χ_1_ torsion angles for active site residues in water and when crowders or substrates are in close contact.

**Figure S22.** Side chain center distances between active site residues as a function of PEG crowder distances.

**Figure S23.** Side chain center distances between active site residues as a function of Ficoll crowder distances.

**Figure S24.** Side chain center distances between active site residues as a function of substrate distances.

**Figure S25.** Sampling of side chain center distances between active site residues in water and when crowders or substrates are in close contact.

**Figure S26.** Catalysis-relevant distances between active site residues (His57:Nδ-H … Oδ1/2- Asp81 and His57-Nε … HOγ-Ser139) as a function of PEG crowder distances.

**Figure S27.** Catalysis-relevant distances between active site residues (His57:Nδ-H … Oδ1/2- Asp81 and His57-Nε … HOγ-Ser139) as a function of Ficoll crowder distances.

**Figure S28.** Catalysis-relevant distances between active site residues (His57:Nδ-H … Oδ1/2- Asp81 and His57-Nε … HOγ-Ser139) as a function of substrate distances.

**Figure S29.** Sampling of catalytically relevant active site distances in water and when crowders or substrates are in close contact.

**Figure S30.** Orientational correlation function for active site side chains (His57, Asp81, Ser139) in NS3.

**Figure S31.** Sampling of distances between the Zn^2+^ ion and the coordinating cysteine residues of NS3 in water and when crowders or substrates are in close contact.

**Figure S32.** Sampling of side chain torsion χ_1_ angles for Zn^2+^ coordinating cysteine residues of NS3 in water and when crowders or substrates are in close contact.

**Table S1.** Simulations of NS3/4A under different conditions analyzed in this study.

**Table S2:** Average Cα coordinate root mean square deviations of NS3/4A

**Table S3:** Average radius of gyration for NS3, NS4A, substrate, and crowders from heavy atoms

**Table S4:** Average secondary structure content for NS4A, and substrate

**Table S5:** Double-exponential fits to rotational correlation functions

**Table S6.** NS3-crowder contact life-times from two-exponential fits to contact survival decays

**Table S7.** Substrate-crowder contact life-times from two-exponential fits to contact survival decays

**Table S8:** Double-exponential fits to the combined orientational correlation functions of the His57, Asp81, and Ser139 active site side chains

## Notes

### Competing Interest Statement

The authors have declared no competing interest.

