## Supplemental Information for "Varying molecular interactions explain crowder-dependent enzyme function of a viral protease"

**Supplementary Methods**
**Figures S1-S32**
**Tables S1-S8**

### Supplementary Methods

##### Building the model of Ficoll crowders

Ficoll crowders were designed to match the size of PEG crowders used in our previous simulations of NS3/4A [1,2] in order to compare the effects of crowding agents of similar size but with different atom composition. We decided to build polysucrose crowders with comparable number of atoms as PEG crowders to achieve molecular weight equivalence rather than molecular volume equivalence. To that extent, we built a 204-atom polymer composed of four sucrose molecules connected with three glycerol linkers (**Figure S1**). Such a molecule matches the 206-atom 28-mer of PEG. This polysucrose crowder also forms a branched structure, a known property of Ficoll polymers. We note that commercially available Ficoll is much larger, with molecular weights of either 70 or 400 kDa.

Since Ficoll is a commercial product, its exact structure has remained unclear [3]. However, we know that Ficoll is synthesized via reaction of sucrose with epichlorohydrin that leads to the formation of a branched polymer glycerols linking sucroses [4]. While the probabilities of attaching glycerol to different hydroxyl groups of a sucrose molecule are unclear, we made the assumption that hydroxyl groups attached to primary carbons of sucrose (GLC:C6, FRU:C1 and FRU:C6) would be more likely to react with epichlorohydrin than hydroxyl groups of secondary carbons. Thus, out of six sucrose-glycerol linking sites in our model, in four cases the terminal groups of glycerol attach to one of the primary carbons, and in two cases - to the C2 carbon of glucose, located opposite the C6 primary carbon. The final structure of our Ficoll-like crowder model used in the simulations is shown in **Figure S1**.

##### Parameterization of the Ficoll crowders

Ficoll crowders were parameterized using the CHARMM force field for carbohydrates [5,6] (**Figure S2**). For sucrose monomers, we used the parameters for α-D-glucose and β-fructofuranose of the AGLC and BFRU residues, along with the SUCR patch, specifically designed to link AGLC and BFRU forming the sucrose molecules. For glycerol the standard MGL residue topology was used.

To create sucrose-glycerol linkages, individual topology patches (shown below) were written for each sucrose-glycerol ester bond, *i.e.*, oxygen and two surrounding carbon atoms. The patch names are related to the atom names of glycerol and saccharide carbons; *e.g.*, the patch GG23 describes the ester bond between the C2 carbon of glucose and C3 carbon of glycerol.

**CHARMM topology:**

```
PRES GG23          -0.27 ! apply to GLC2:C2 -- MGL3:C3
dele atom 1HO2
dele atom 2O3
dele atom 2HO3
GROU
ATOM 1C2  CC3161    0.09 !   like MELZ:2C3
ATOM 1O2  OC301    -0.36
ATOM 2C3  CC322     0.00 !   like IMAL:2C6
BOND 1O2  2C3

PRES GF11          -0.36 ! apply to MGL:C1 -- FRU:C1
dele atom 1HO1
dele atom 1O1
dele atom 2HO1
GROU
ATOM 1C1  CC322     0.00  !   like IMAL:2C6
ATOM 2O1  OC301    -0.36
ATOM 2C1  CC321     0.00   !   like IMAL:2C6
BOND 1C1  2O1

PRES GF31          -0.36  ! apply to MGL:C3--FRU:C1
dele atom 1HO3
dele atom 1O3
dele atom 2HO1
GROU
ATOM 1C3  CC322     0.00  !   like IMAL:2C6
ATOM 2O1  OC301    -0.36
ATOM 2C1  CC321     0.00  !   like IMAL:2C6
BOND 1C3  2O1

PRES FG61           -0.36 ! apply to FRU:C6 -- MGL:C1
dele atom 1HO6
dele atom 2O3
dele atom 2HO3
GROU
ATOM 1C6  CC321     0.00  !   like IMAL:2C6
ATOM 1O6  OC301    -0.36
ATOM 2C1  CC322     0.00  !   like IMAL:2C6
BOND 1O6  2C1
```

The sucrose-glycerol linkages were parameterized by analogy to the molecules already parameterized in the CHARMM force field. Considering the structural similarity between the glycerol and C1O1H-C2O2H-C3O3H part of glucose (**Figure S2**), we assigned the force field parameters based on the similarity to glycosidic bonds found in certain di- and trisaccharides.

All the ester bond oxygens were assigned the partial charge of -0.36 e and the C-O-C angle of 109.7$^{o}$. These are the standard values for glycosidic bonds linking glucose and fructose molecules.

Carbon atoms taking part in the ester bond formation were divided into two groups:

1. secondary carbons, bonding with two carbon atoms. This group includes the GLC:C2 atom, present in the GG23 patch. The C2 carbon was parameterized based on the 2C3 carbon in melezitose patch (MELZ), which is one of just a few cases in the CHARMM force field where the secondary carbon is involved in the glycosidic bond formation (**Figure S3**).

2. primary carbons, bonding with one carbon atom. This group includes both the primary carbon of glucose and fructose (GLC:C6, FRU:C1, FRU:C6) and terminal carbons of glycerol (GLY:C1, GLY:C3). Each of those carbons were assigned partial charges of 0.0 and the type CC321, analogous to the 2C6 carbon of isomaltulose (IMAL).

The patches described here include only the atoms for which the charges are different from free monomers. After adding the appropriate number of hydrogen atoms, whose partial charge always equal 0.09 e, the total charges for each patch sum up to zero.

The missing angle and dihedral parameters, as well as one set of bond parameters, were assigned individually based on the similarity to already parameterized di- and trisaccharides containing fructose and glucose.

**CHARMM parameters:**

```
BONDS
CC3151  OC301    360.00  1.415   ! CC3162 OC302

ANGLES
CC3161  OC301   CC312    50.00   109.20  !
CC322   OC301   CC321    95.00   109.70
CC3161  OC301   CC322    95.00   109.70  ! IMAL 1C1-1O1-2C6
CC3051  CC3151  OC301    45.00   110.50  ! FRU OC303-C3-C2
HCA1    CC3151  OC301    60.00   109.50  ! FRU OC303-C3-HCA1
OC301   CC3151  CC3151   45.00   110.50  ! OC303 CC3151 CC3151
CC3151  OC301   CC322    95.00   109.70  ! IMAL 1C1-1O1-2C6

DIHEDRALS
CC3161  CC3161  OC301   CC312    0.13  1  180.0 ! CC3161 CC3161 OC301
CC3162
CC3161  CC3161  OC301   CC312    0.25  2  180.0
CC3161  CC3161  OC301   CC312    0.06  3  180.0

HCA1    CC312   OC301   CC3161   0.284 3    0.0 ! HCA2 CC321 OC301 CC3152
HCA1    CC3161  OC301   CC312    0.284 3    0.0 ! HCA1 CC3152 OC301 CC321

OC301   CC312   CC322   OC301    1.7749 1 180.00 ! OC301 CC312 CC322 OC311
OC301   CC312   CC322   OC301    1.5713 2   0.00
OC301   CC312   CC322   OC301    1.8214 3   0.00

CC3051  CC321   OC301   CC322    0.64  1  180.0 ! CC3051 CC321 OC301 CC3051
CC3051  CC321   OC301   CC322    0.03  2  180.0
CC3051  CC321   OC301   CC322    0.61  3    0.0
CC322   OC301   CC321   HCA2     0.284 3    0.0 ! CC331 OC301 CC321 HCA2

CC321   OC301   CC322   HCA2     0.20  3    0.0 ! CC322 CC312 CC322 HCA2

CC322   CC312   CC322   OC301    0.35  1    0.0 ! CC322 CC312 CC322 OC311
CC322   CC312   CC322   OC301    0.69  2    0.0
CC322   CC312   CC322   OC301    2.79  3  180.0
OC3C51  CC3051  CC3151  OC301    0.32  1  180.0 ! OC3C51 CC3051 CC3151 OC303
OC3C51  CC3051  CC3151  OC301    0.65  2  180.0 !
OC3C51  CC3051  CC3151  OC301    2.62  3    0.0 !

CC322   OC301   CC3151  CC3051   0.07  1  180.0 ! CC3162 OC303 CC3151 CC3051
CC322   OC301   CC3151  CC3051   0.04  2    0.0 !
CC322   OC301   CC3151  CC3051   0.14  3    0.0 !
CC321   CC3051  CC3151  OC301    0.94  1    0.0 ! CC321 CC3051 CC3151 OC303
CC321   CC3051  CC3151  OC301    1.59  2  180.0 !
CC321   CC3051  CC3151  OC301    0.84  3    0.0 !
CC3153  CC3151  CC3151  OC301    0.01  1  180.0 ! CC3153 CC3151 CC3151 OC303
CC3153  CC3151  CC3151  OC301    0.72  2    0.0 !               "
```

```
CC3153  CC3151  CC3151  OC301    0.73  3    0.0 !

CC322   OC301   CC3151  HCA1     0.284 3    0.0 ! CC3162 OC303 CC3151 HCA1
OC302   CC3051  CC3151  OC301    0.12  1  180.0 ! OC302 CC3051 CC3151 OC303
OC302   CC3051  CC3151  OC301    1.87  2  180.0 !
OC302   CC3051  CC3151  OC301    1.64  3  180.0 !
OC301   CC3151  CC3151  HCA1     0.14  3    0.0 ! OC311 CC3151 CC3151 HCA1
OC301   CC3151  CC3151  OC311    2.87  1  180.0 ! OC311 CC3151 CC3151 OC311
OC301   CC3151  CC3151  OC311    0.03  2    0.0 !
OC301   CC3151  CC3151  OC311    0.23  3    0.0 !

CC3151  OC301   CC322   HCA2     0.284 3    0.0 ! CC3062 OC301 CC321 HCA2
CC3161  OC301   CC322   HCA2     0.284 3    0.0 ! CC3062 OC301 CC321 HCA2

CC3161  CC3161  OC301   CC322    0.13  1  180.0 ! CC3161 CC3161 OC301 CC3162
CC3161  CC3161  OC301   CC322    0.25  2  180.0 !
CC3161  CC3161  OC301   CC322    0.06  3  180.0

CC3162  CC3161  OC301   CC322    0.13  1  180.0 ! CC3161 CC3161 OC301 CC3162
CC3162  CC3161  OC301   CC322    0.25  2  180.0 !
CC3162  CC3161  OC301   CC322    0.06  3  180.0

HCA1    CC3161  OC301   CC322    0.284 3    0.0 ! HCA1 CC3161 OC301 CC3152
CC322   OC301   CC3151  CC3151   0.07  1  180.0 ! CC3162 OC303 CC3151 CC3151
CC322   OC301   CC3151  CC3151   0.04  2    0.0
CC322   OC301   CC3151  CC3151   0.14  3    0.0

OC301   CC3161  CC3162  OC302    0.59  1  180.0 ! OC301 CC3161 CC3162 OC301
OC301   CC3161  CC3162  OC302    1.16  2    0.0

CC322   CC312   OC301   CC3161   0.40  1  180.0 ! CC321 CC3051 OC302 CC3162
CC322   CC312   OC301   CC3161   0.48  2    0.0 !
CC322   CC312   OC301   CC3161   0.19  3    0.0 !

CC312   CC322   OC301   CC321    0.64  1  180.0 ! CC3051 CC321 OC301 CC3051
CC312   CC322   OC301   CC321    0.03  2  180.0
CC312   CC322   OC301   CC321    0.61  3    0.0

CC312   CC322   OC301   CC3151   0.64  1  180.0 ! CC3051 CC321 OC301 CC3051
CC312   CC322   OC301   CC3151   0.03  2  180.0
CC312   CC322   OC301   CC3151   0.61  3    0.0

CC312   CC322   OC301   CC3161   0.64  1  180.0 ! CC3051 CC321 OC301 CC3051
CC312   CC322   OC301   CC3161   0.03  2  180.0
CC312   CC322   OC301   CC3161   0.61  3    0.0

CC3153  CC321   OC301   CC322    0.64  1  180.0 ! CC3153 CC321 OC301 CC3162
CC3153  CC321   OC301   CC322    0.03  2  180.0
CC3153  CC321   OC301   CC322    0.61  3    0.0
```

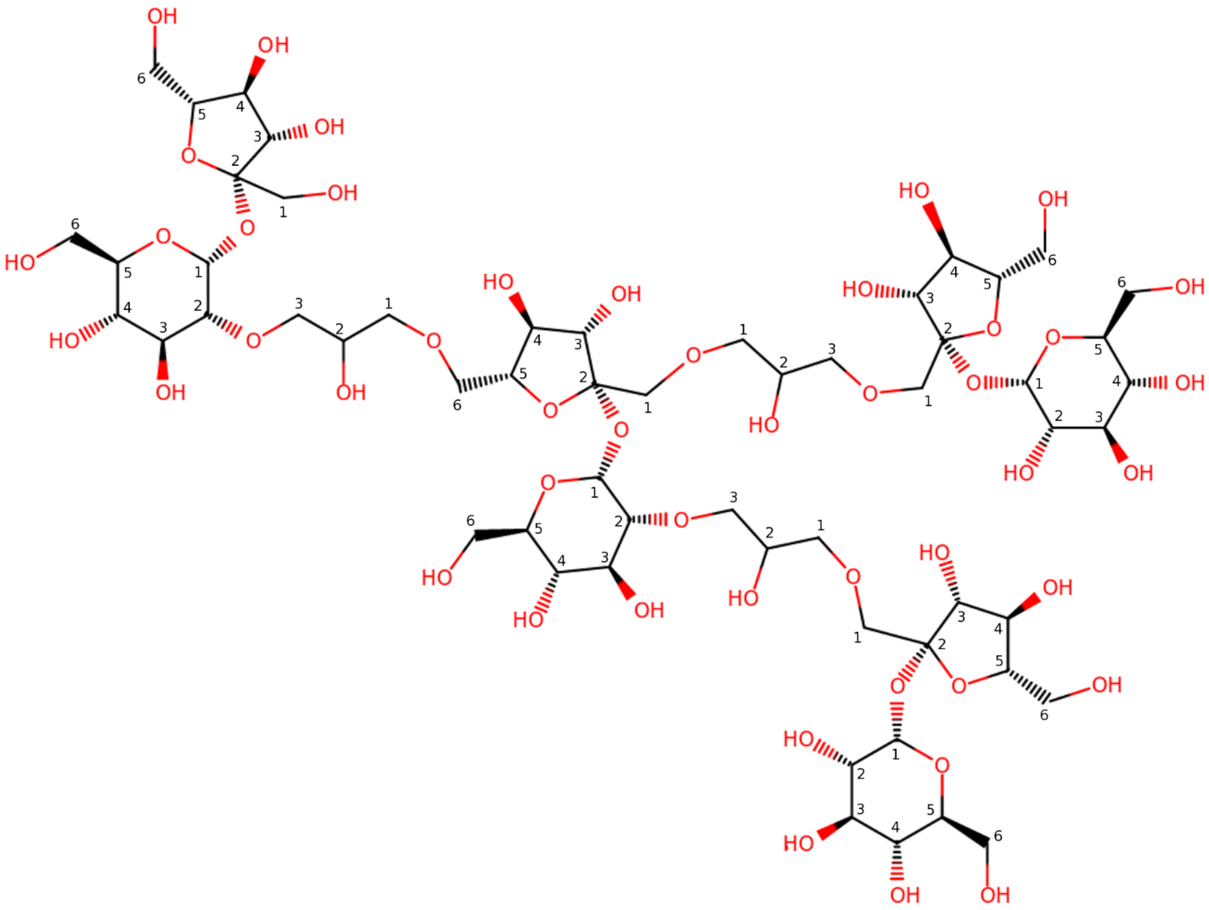

**Figure S1.** Chemical structure of the polysucrose molecule used as a model for Ficoll. The sucrose molecules are connected with glycerol linkers. Carbon atoms are annotated according to the numbers used in the CHARMM force field.

glucose, AGLC

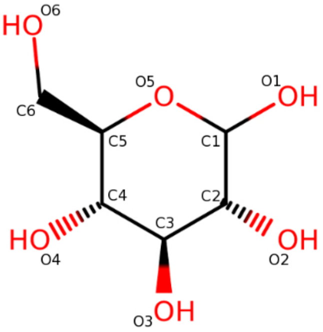


fructose, BFRU

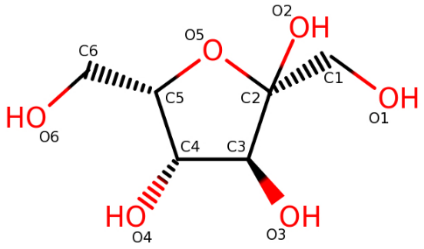


sucrose

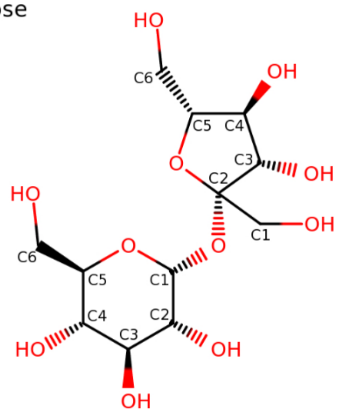


```
PRES SUCR          0.00 ! apply to AGLC,BFRU
!dele atom 1HO1
!dele atom 2O2
!dele atom 2HO2
GROU
ATOM 1C1  CC3162    0.29 !
ATOM 1O1  OC302    -0.36 !
ATOM 2C2  CC3051    0.38 !
ATOM 1H1  HCA1      0.09 !
ATOM 1C5  CC3163    0.11 !
ATOM 1H5  HCA1      0.09 !
ATOM 1O5  OC3C61   -0.40 !
ATOM 2O5  OC3C51   -0.40 !
ATOM 2C5  CC3153    0.11 !
ATOM 2H5  HCA1      0.09 !
BOND 1O1  2C2
```

**Figure S2.** Building blocks used to parameterize Ficoll molecules. Glucose and fructose are shown, together forming sucrose molecules. All structures are shown along with atom names used in the CHARMM topology.

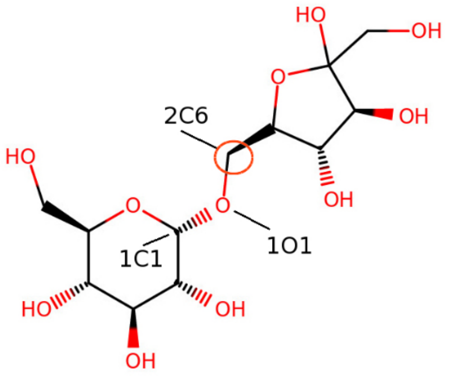


```
PRES IMAL          -0.07 ! pram apply to AGLC,BFRU
dele atom 1HO1
dele atom 2O6
dele atom 2HO6
ATOM 1C1  CC3162    0.29
ATOM 1O1  OC301    -0.36
ATOM 2C6  CC321     0.00
BOND 1O1  2C6
```

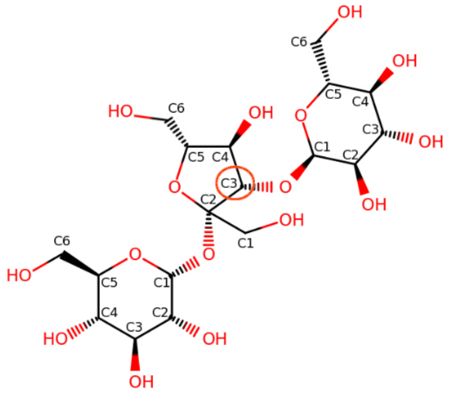


```
PRES MELZ           0.33 !  apply to AGLC,BFRU,AGLC
dele atom 1HO1
dele atom 2O2
dele atom 2HO2
dele atom 3HO1
dele atom 2O3
dele atom 2HO3
ATOM 1C1  CC3162    0.29 !
ATOM 1O1  OC302    -0.36 !
ATOM 2C2  CC3051    0.38 !
ATOM 3C1  CC3162    0.29 !
ATOM 3O1  OC303    -0.36 !
ATOM 2C3  CC3151    0.09 !
BOND 1O1  2C2  3O1  2C3
```

**Figure S3.** Structures of isomaltulose and melezitose and CHARMM force field patches. The patches are used to form these molecules from glucose and fructose monomers. The 2C6 and 2C3 atoms used to parameterize carbon atoms in the model of Ficoll are marked with orange circles.

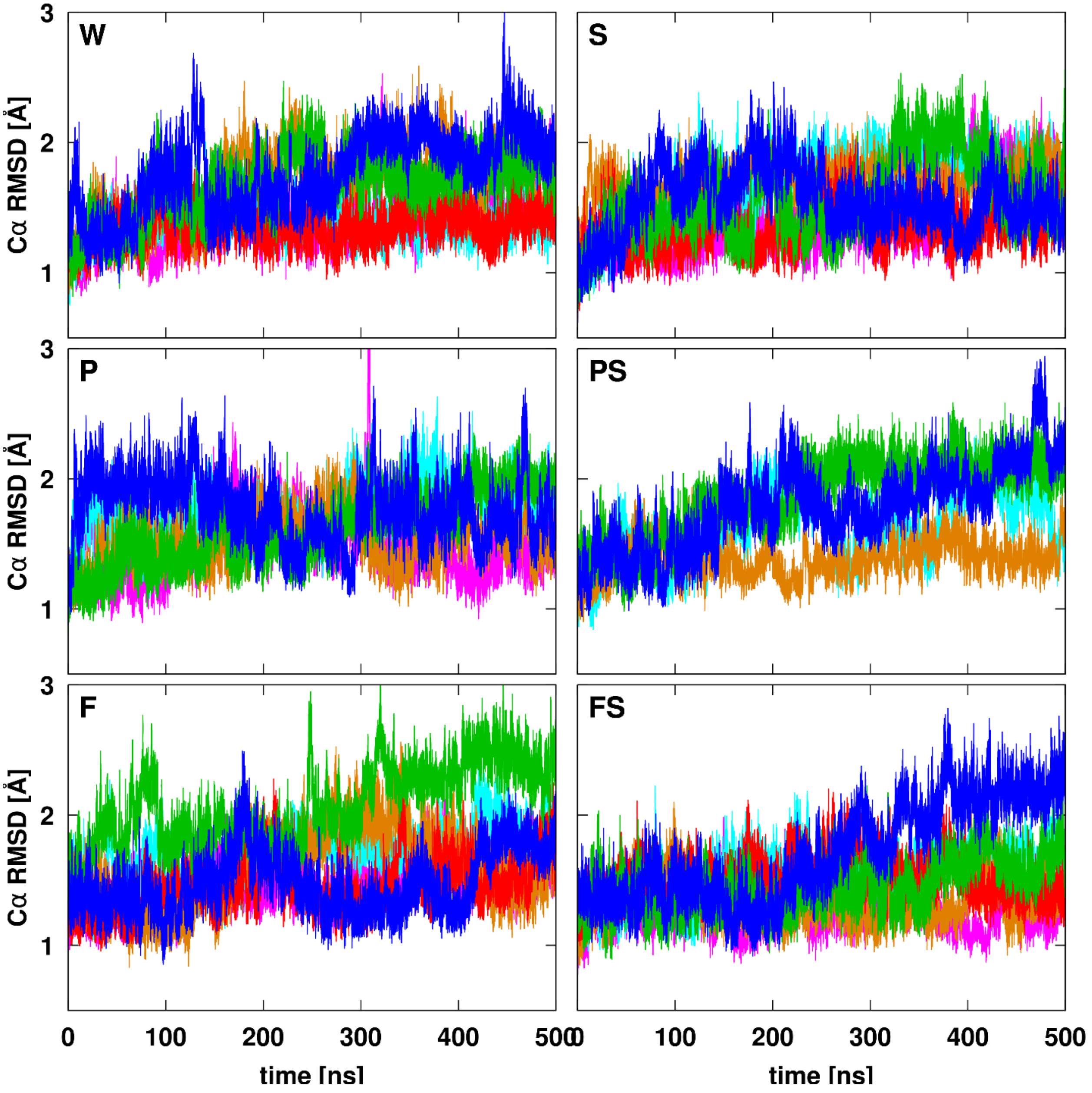


**Figure S4**. Time evolution of coordinate root mean square deviations of NS3/4A. RMSD values are shown in water (W), in the presence of PEG (P) and Ficoll (F) and with substrates (S, PS, FS). RMSD values are based on Cα coordinates after optimal superposition with respect to the experimental structure (PDB ID: 4JMY) for NS3 and the central 13 residues of NS4A for which structure information is available in the PDB. Different colors distinguish individual trajectories.

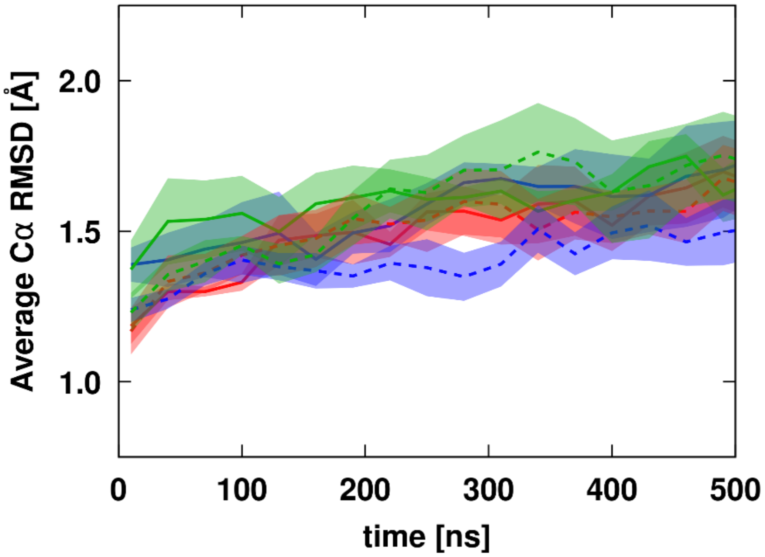


**Figure S5**. Time-averaged Cα root mean square deviations as a function of simulation time. Results are shown for NS3/4A in water (red), in the presence of PEG (green) or Ficoll (blue). Solid and dashed lines are from simulations in the absence and presence of substrates, respectively. Averages were calculated over 20 ns trajectory segments. Error bars indicate the standard errors of the mean from variations between replicate simulations.

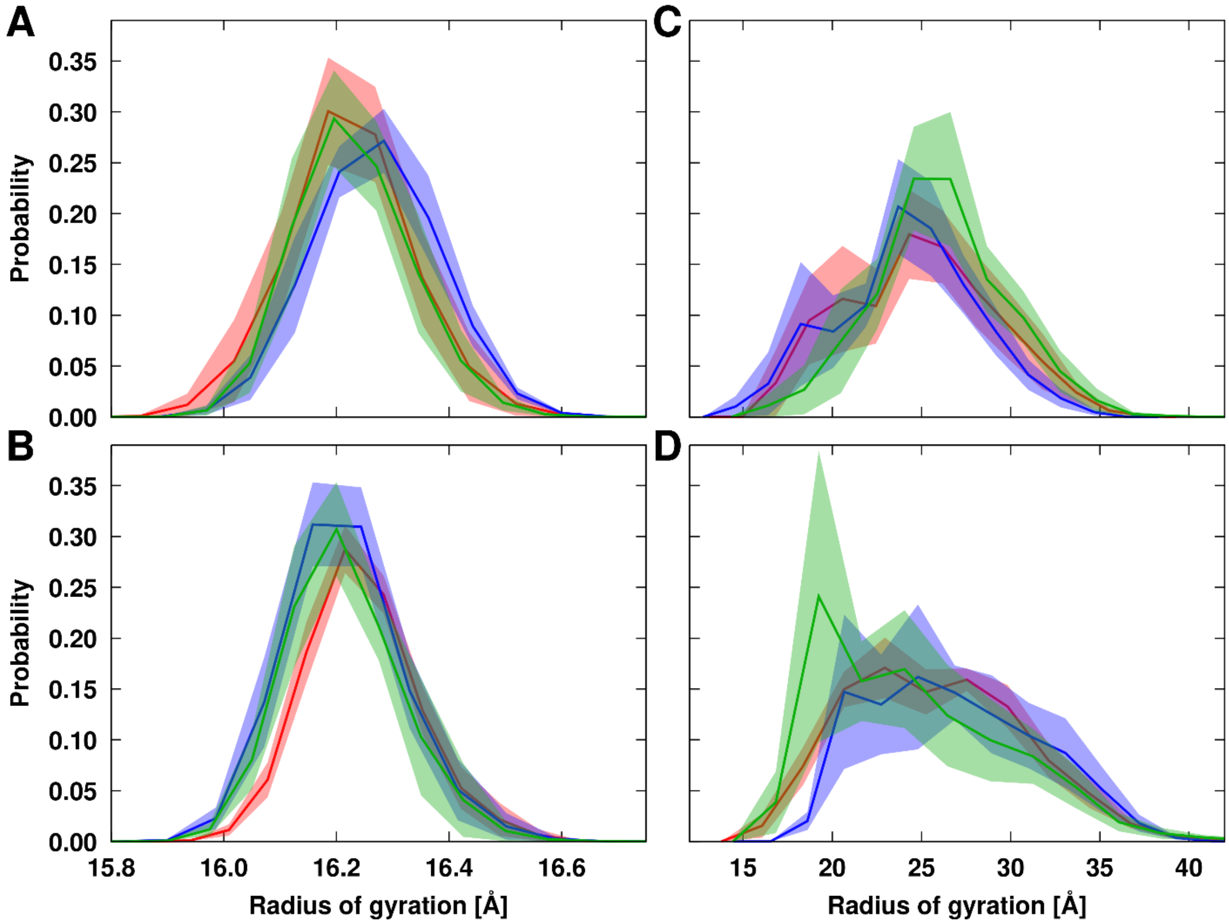


**Figure S6.** Radius of gyration distributions of NS3 and NS4A. Results are shown for NS3 (A,B) and NS4A (C,D) without (A,C) and with (B,D) substrates in water (red) or in the presence of PEG (green) or Ficoll (blue) crowders. Individual histograms were averaged over replicate trajectories. Error bars indicate standard errors for each bin.

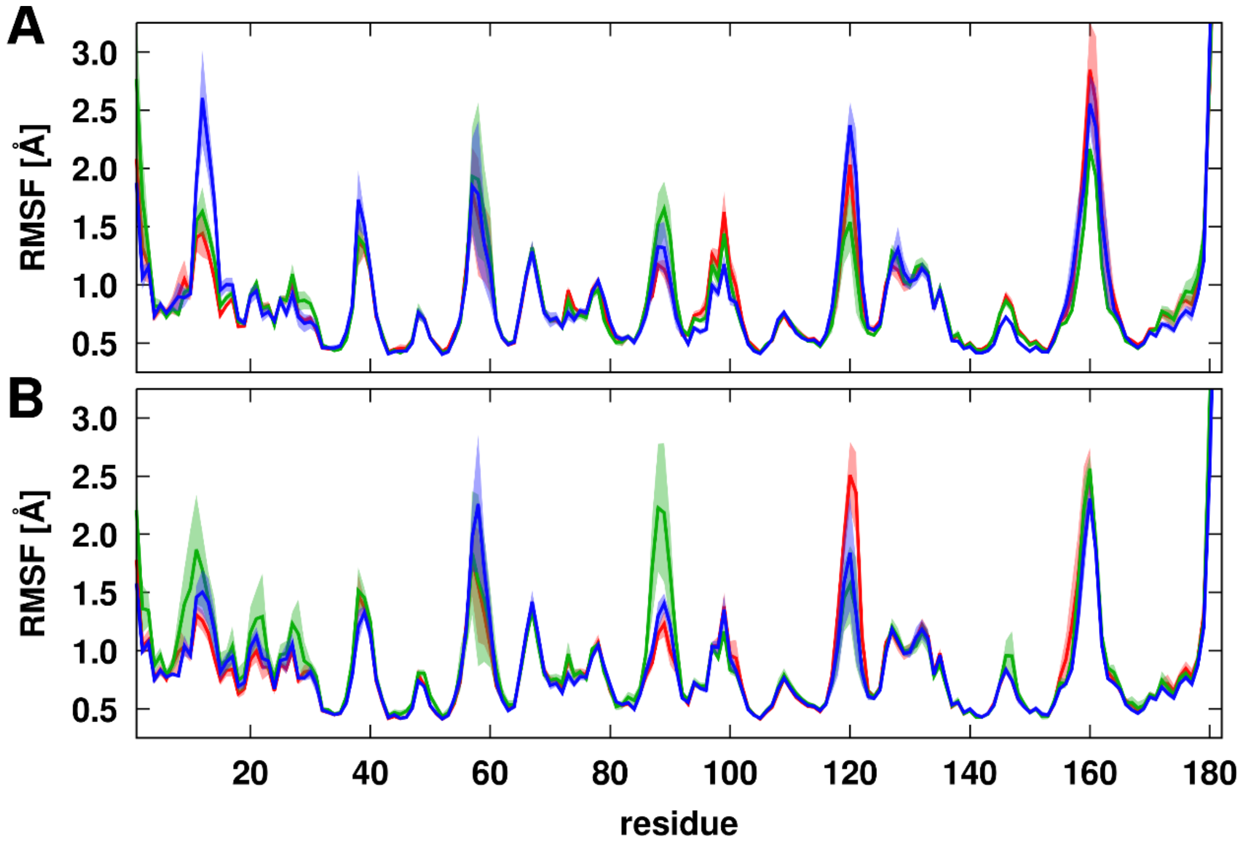


**Figure S7**. Root mean square fluctuations of NS3. Results are shown without (A) and with (B) substrates in water (red) or in the presence of PEG (green) or Ficoll (blue) crowders. RMSF was calculated from Cα atoms with respect to the average structures of NS3 for each trajectory after superposition onto a reference structure. Solid lines reflect trajectory averages, shaded areas indicate standard errors of the mean.

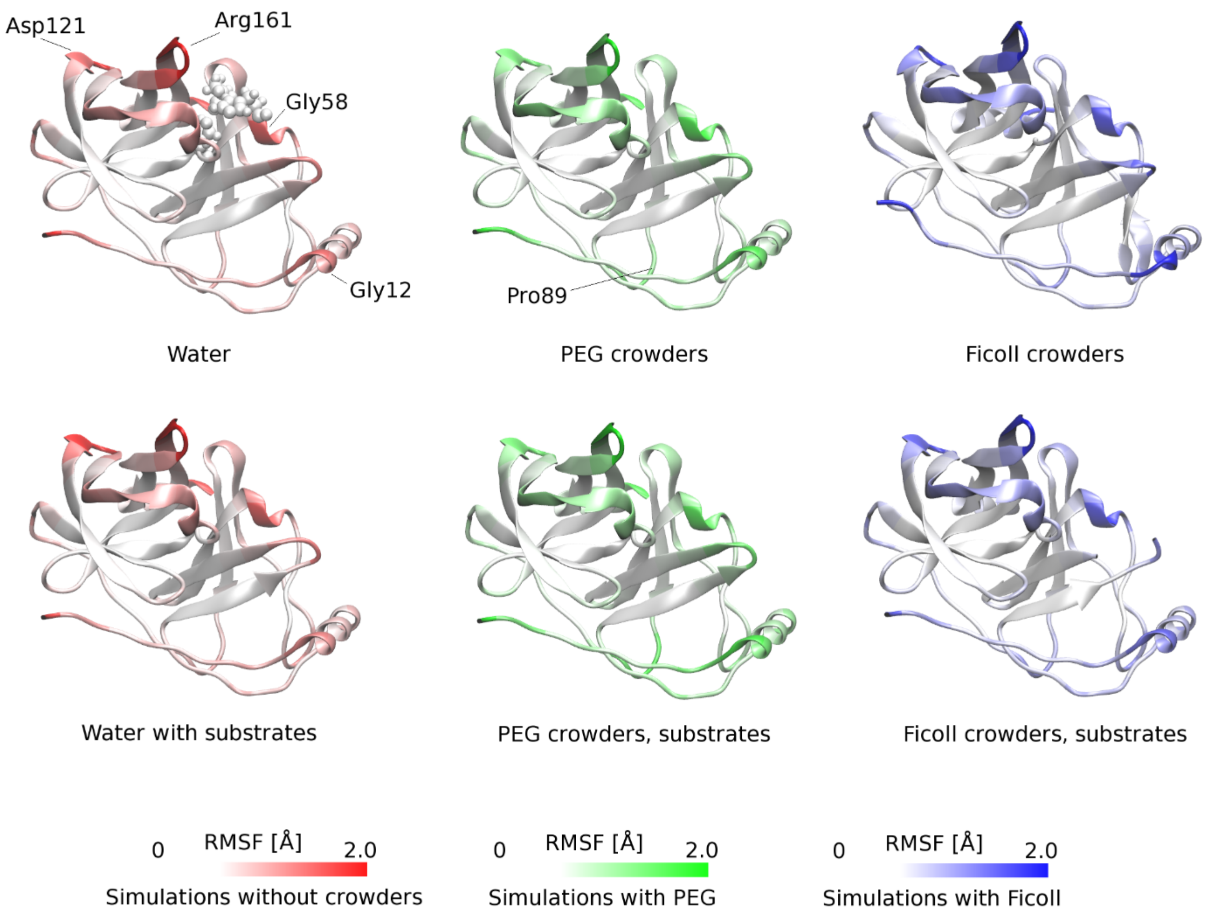


**Figure S8.** Root mean square fluctuations of NS3 in non-crowded and crowded environments projected onto the protease structure. RMSF values were calculated for the $C_\alpha$ atoms of each amino acid, with respect to the average structure for each trajectory, after superposition onto the crystallographic structure (PDB ID: 4JMY). For clarity, in all cases the higher end of the RMSF scale (dark colors) in the legends correspond to RMSF values between 2.0 and the maximum value of 5.23 Å.

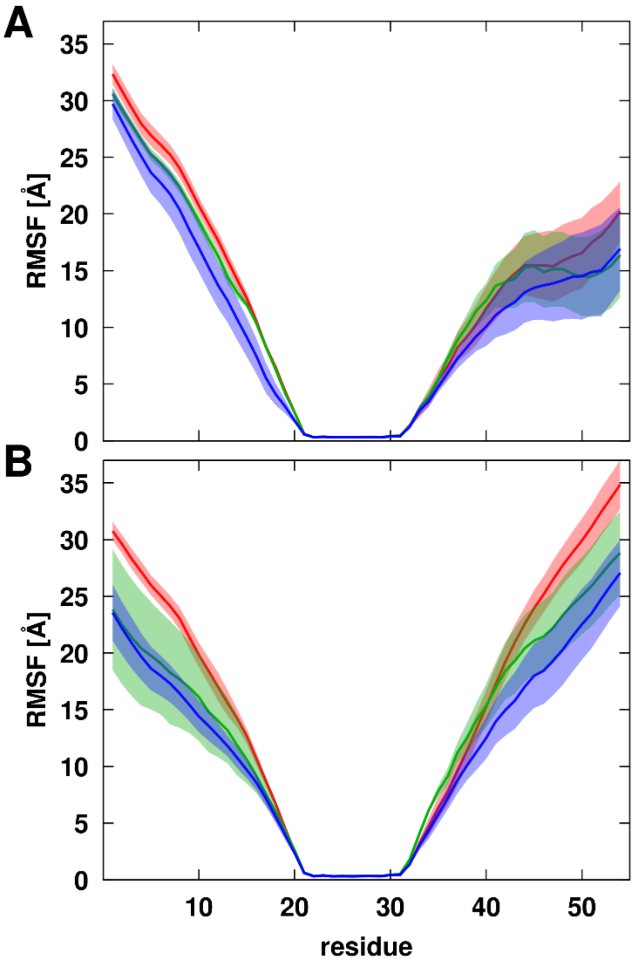


**Figure S9.** Root mean square fluctuations of NS4A. Results are shown without (A) and with (B) substrates in water (red) or in the presence of PEG (green) or Ficoll (blue) crowders. RMSF was calculated from Cα atoms with respect to the average structures of NS4A for each trajectory after superposition of the ordered central residues (21-31). Solid lines reflect trajectory averages, shaded areas indicate standard errors of the mean.

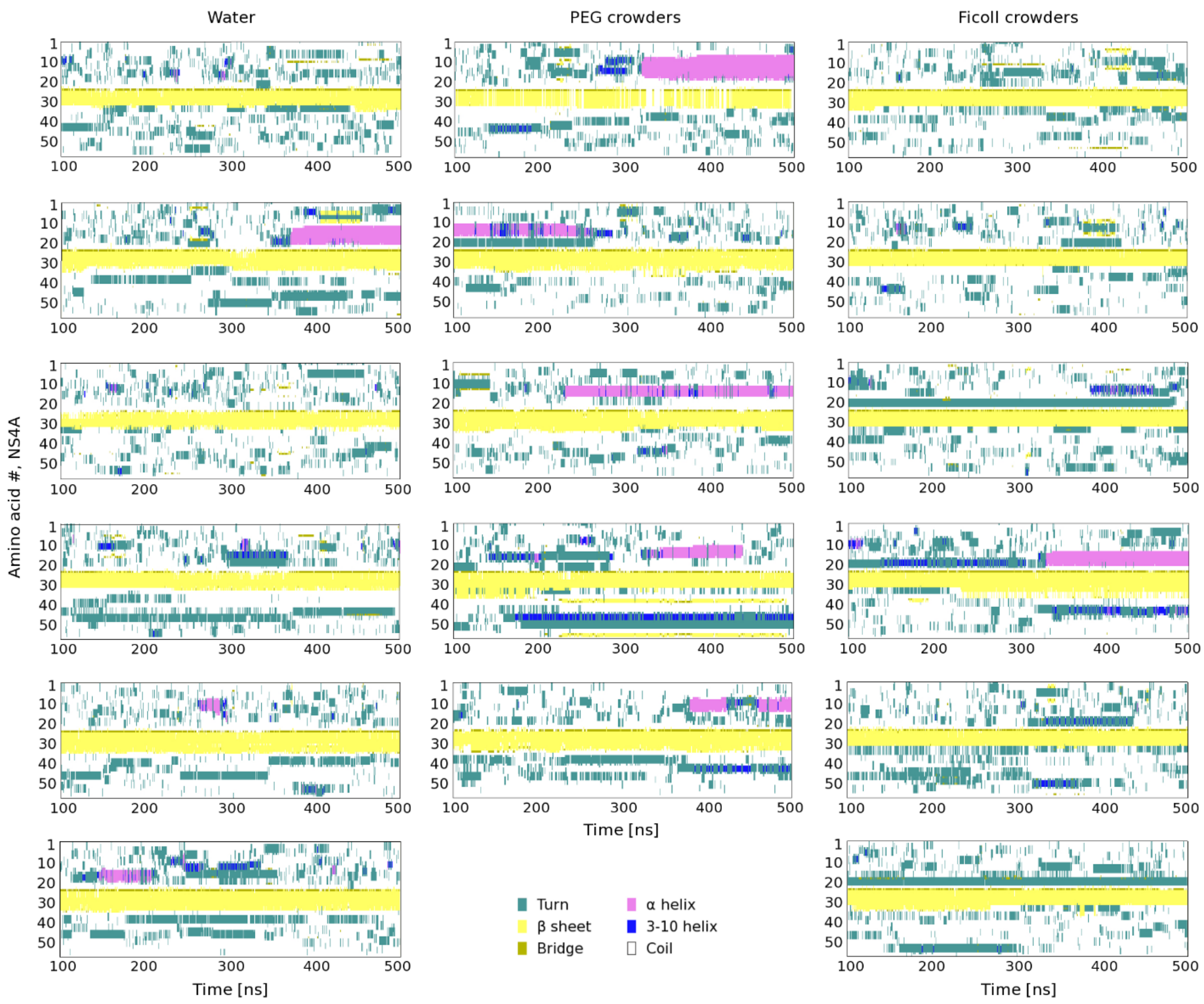


**Figure S10.** Secondary structure evolution of the NS4A cofactor in the simulations with water and with either PEG or Ficoll crowders. Helical conformations are marked in pink or blue, β-strands in yellow, and turns in cyan. Each graph shows data from one MD trajectory.

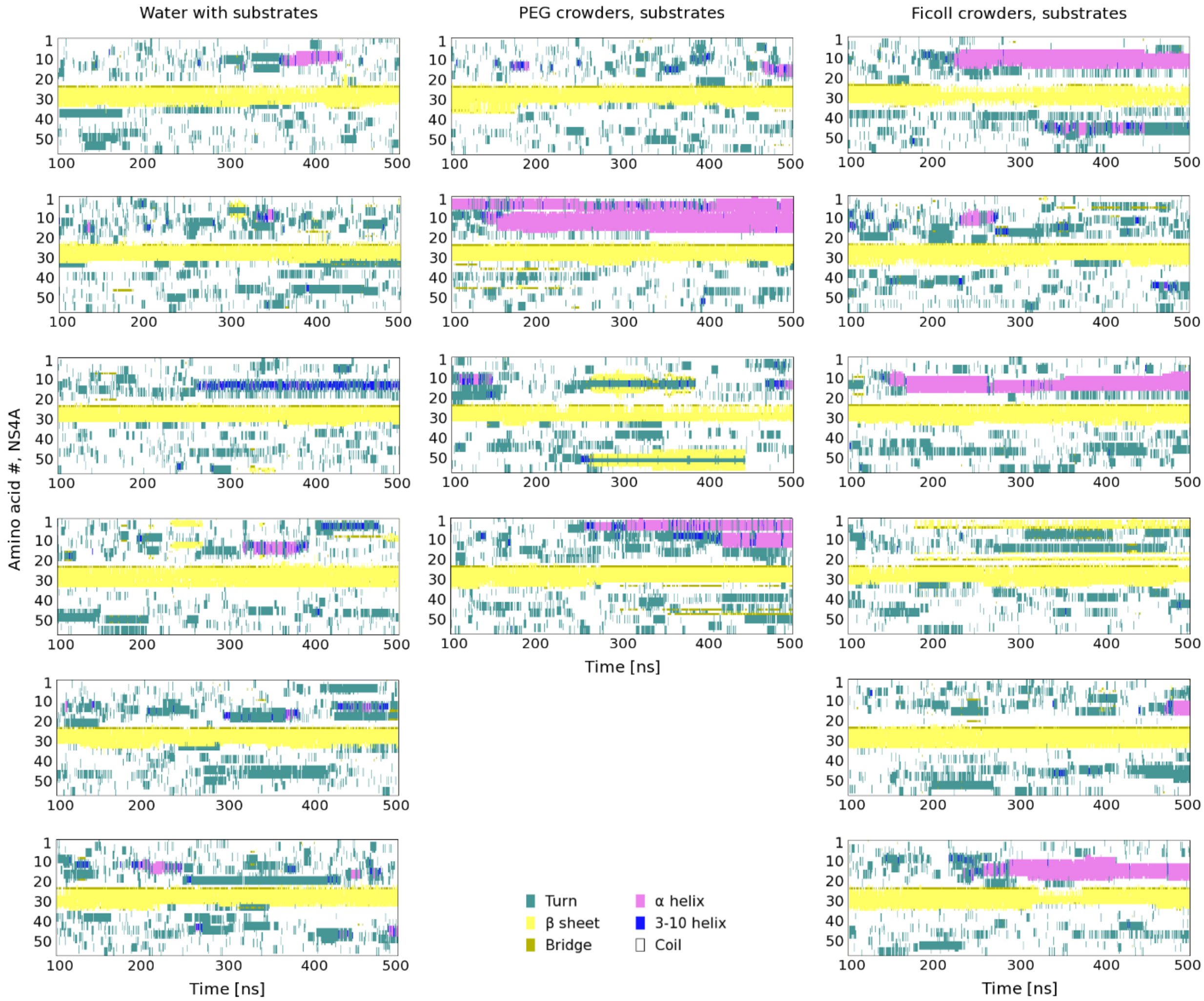


**Figure S11.** Secondary structure evolution of the NS4A cofactor in the simulations with substrates and with either PEG or Ficoll crowders. Helical conformations are marked in pink or blue, β-strands in yellow, and turns in cyan. Each graph shows data from one MD trajectory.

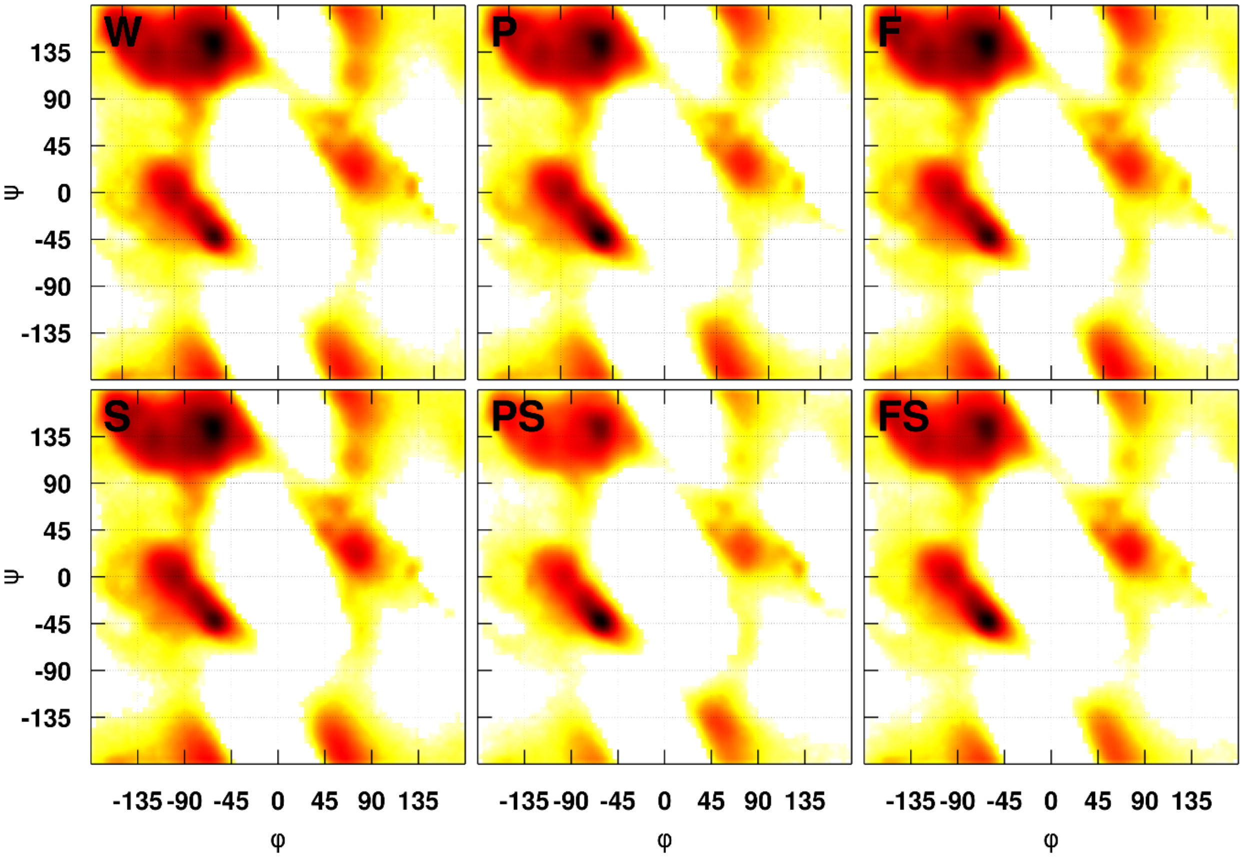


**Figure S12.** Ramachandran map of backbone torsion angles φ and ψ for N-terminal residues (residues 2-19) of NS4A. Results are shown from sampling in water (W), in the presence of PEG (P), in the presence of Ficoll (F), with substrates (S), with substrates in the presence of PEG (PS), and with substrates in the presence of Ficoll (FS).

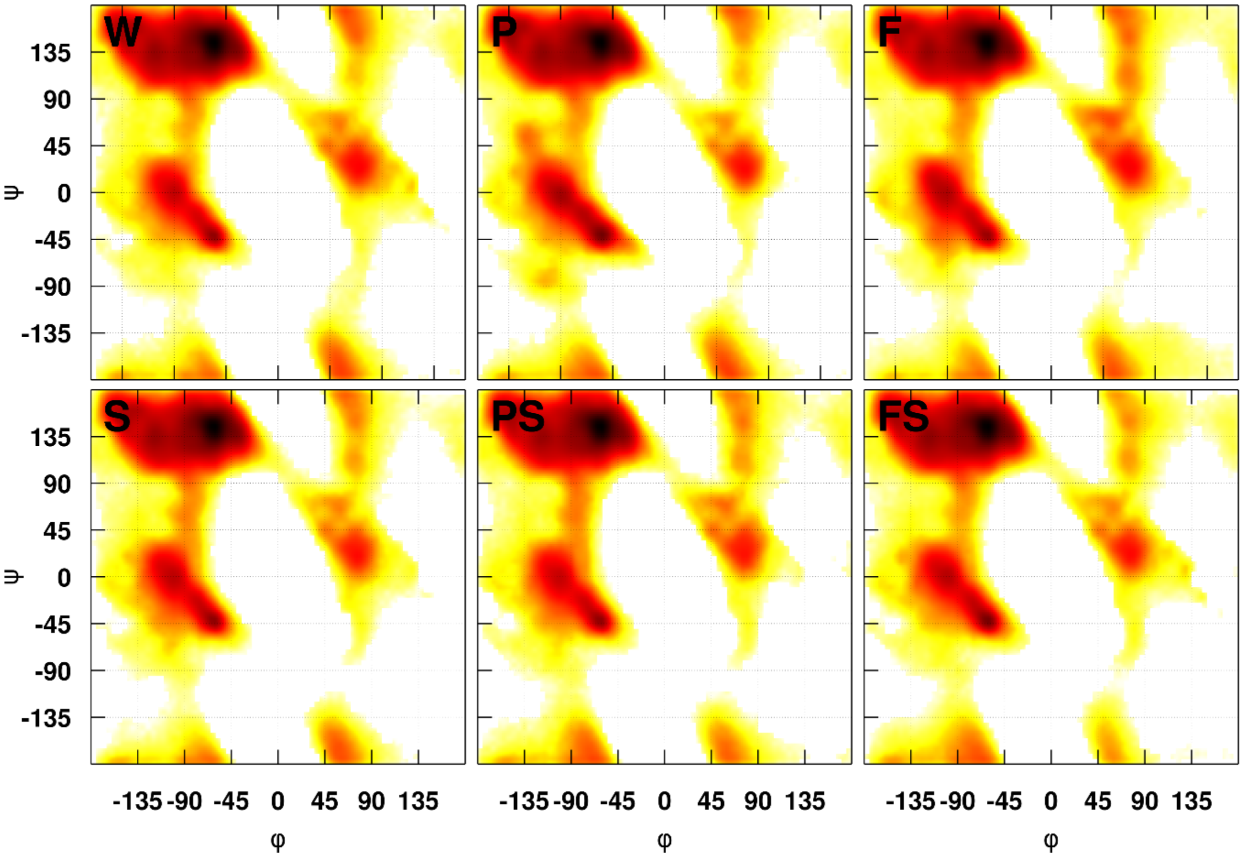


**Figure S13**. Ramachandran map of backbone torsion angles φ and ψ for C-terminal residues (residues 31-53) of NS4A. Results are shown from sampling in water (W), in the presence of PEG (P), in the presence of Ficoll (F), with substrates (S), with substrates in the presence of PEG (PS), and with substrates in the presence of Ficoll (FS).

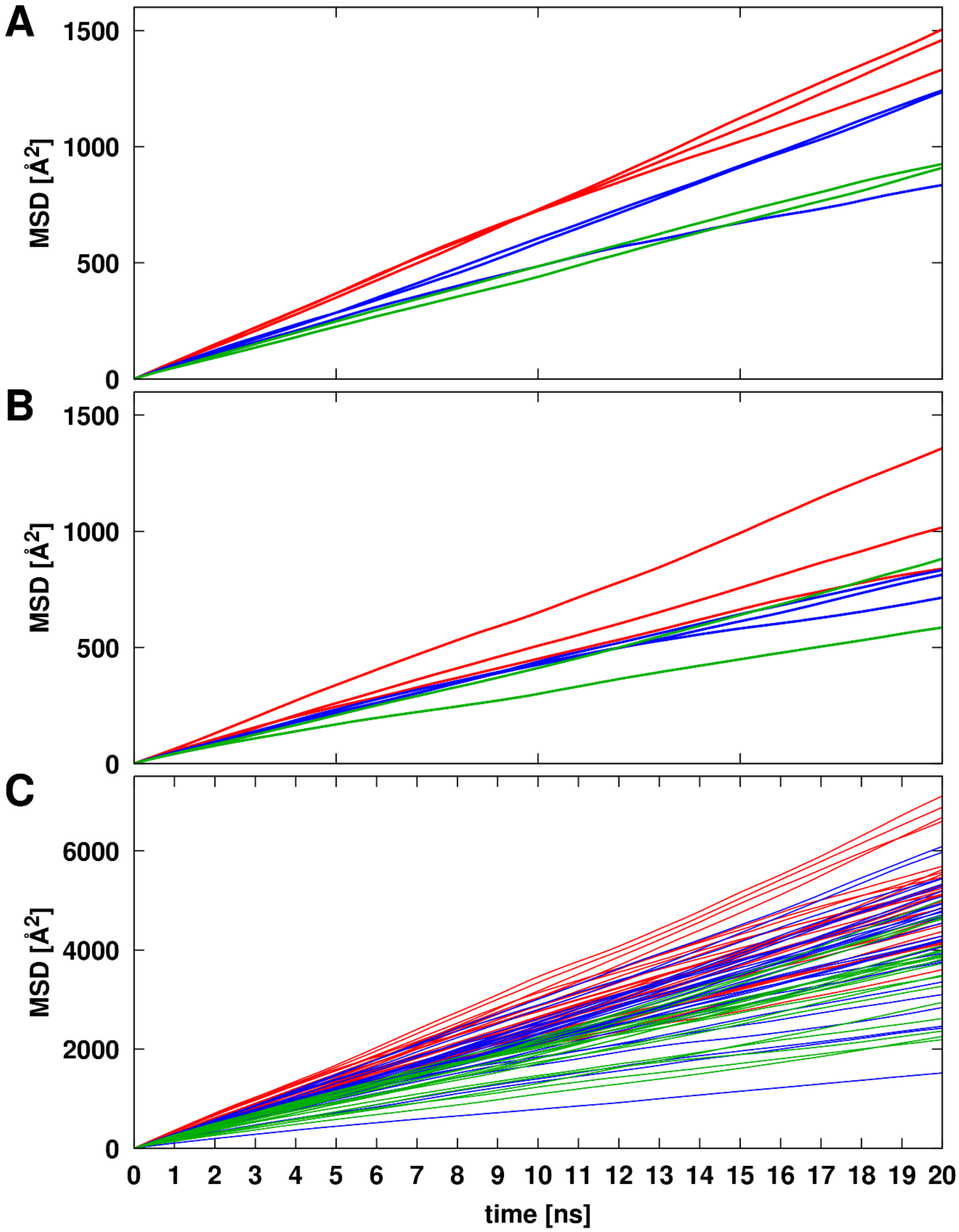


**Figure S14**. Mean square displacement of centers of mass vs. time used for determining translation diffusion coefficients for NS3/4A. Results are shown without substrates (A), with substrates (B), and for substrates (C) in simulations with water (red), in the presence of PEG (green), and in the presence of Ficoll (blue).

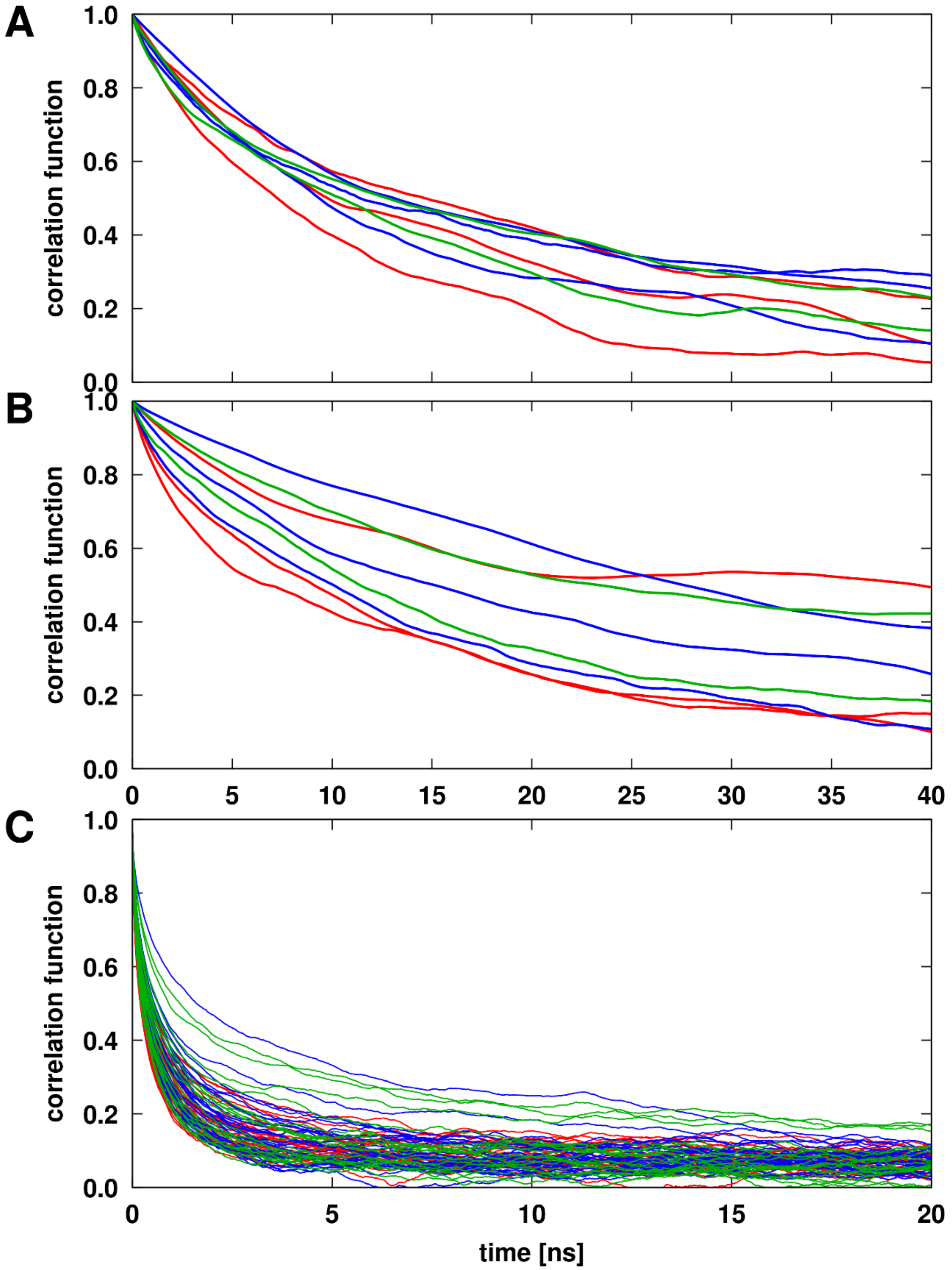


**Figure S15.** Rotational correlation functions used for determining rotational diffusion coefficients for NS3/4A. Results are shown without substrates (A), with substrates (B), and for substrates (C) in simulations with water (red), in the presence of PEG (green), and in the presence of Ficoll (blue).

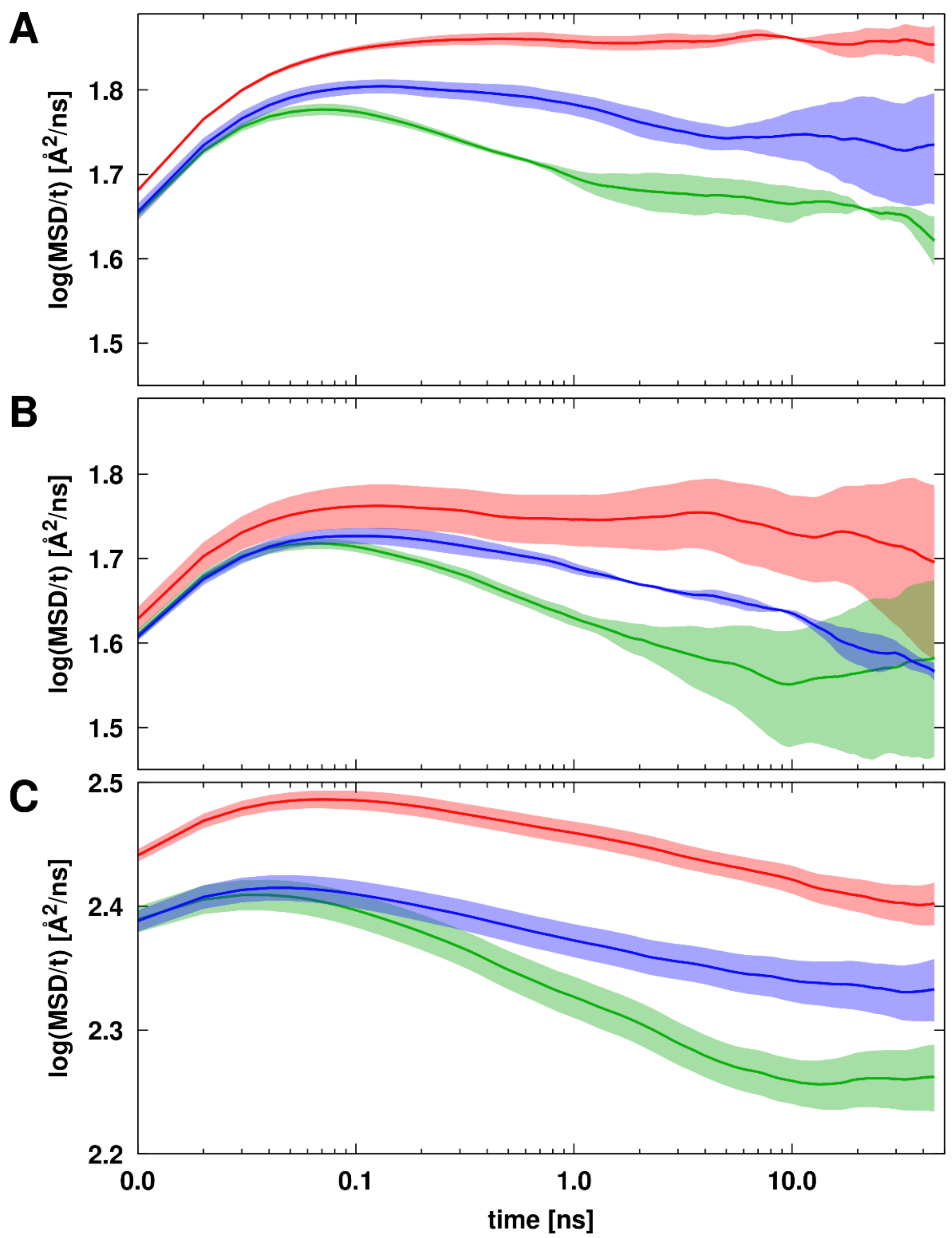


**Figure S16.** MSD/t vs. time on a log-log scale based on trajectory averaged mean square displacement curves. The underlying MSD curves are shown in **Figure S14**. Results are shown for NS3/4A without substrates (A), with substrates (B), and for substrates (C) in simulations with water (red), in the presence of PEG (green), and in the presence of Ficoll (blue). Shaded areas indicate standard errors of the mean.

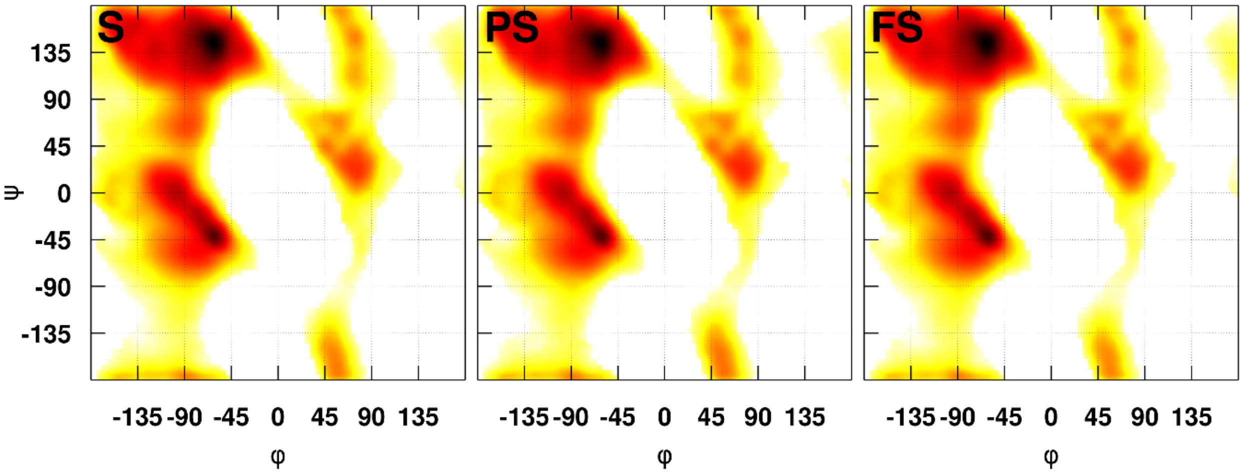


**Figure S17.** Ramachandran map of backbone torsion angles φ and ψ for substrates. Results from sampling substrates with NS3/4A in water (S), in the presence of PEG (PS), and in the presence of Ficoll (FS).

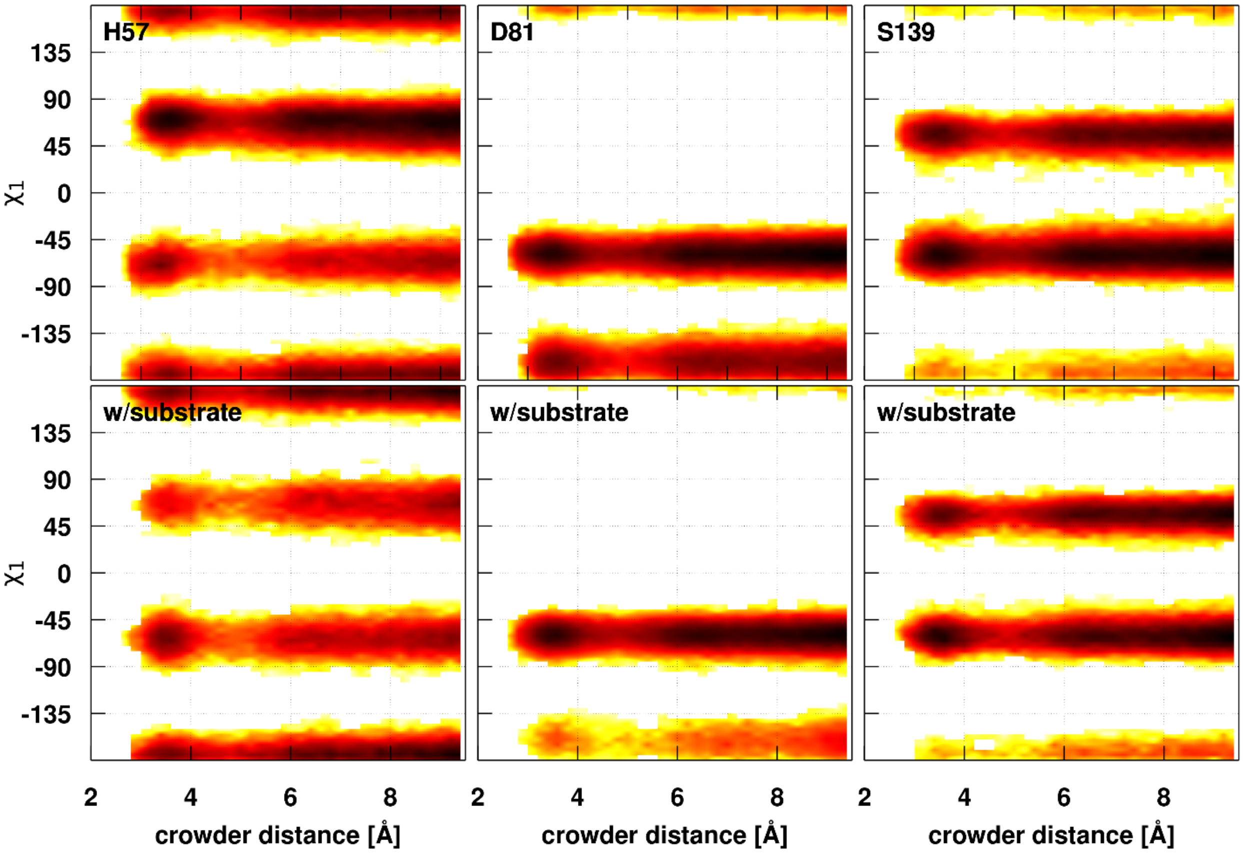


**Figure S18.** Side chain $\chi_1$ torsion angle sampling of active site residues as a function of closest PEG crowder distance. The distance was calculated to any of active site residues in simulations without (top row) and with (bottom row) substrates. In the crystal structure (4JMY), the values of the $\chi_1$ torsion are 76.6° for His57, -161.4° for Asp81, and -82.6° for Ser139.

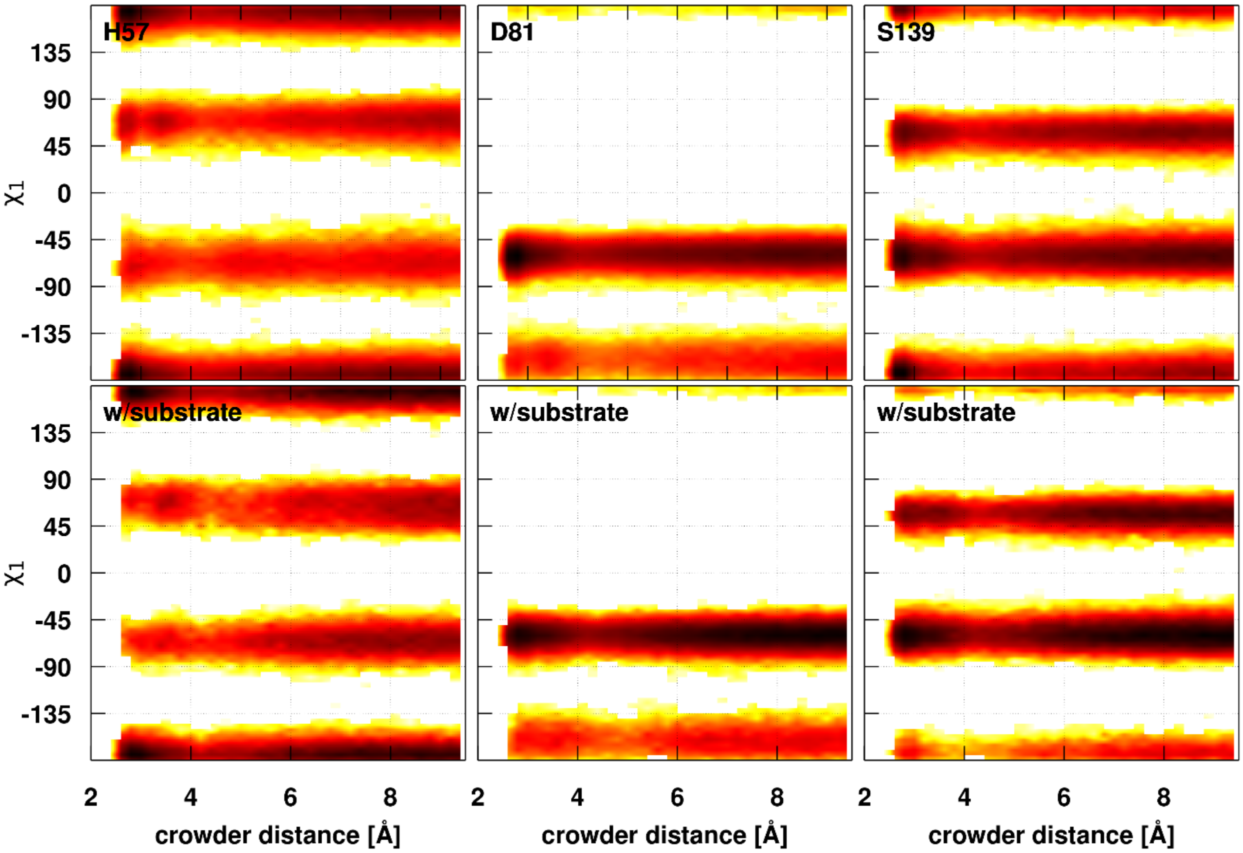


**Figure S19.** Side chain $\chi_1$ torsion angle sampling of active site residues as a function of Ficoll crowder distances. See **Figure S18** for additional details.

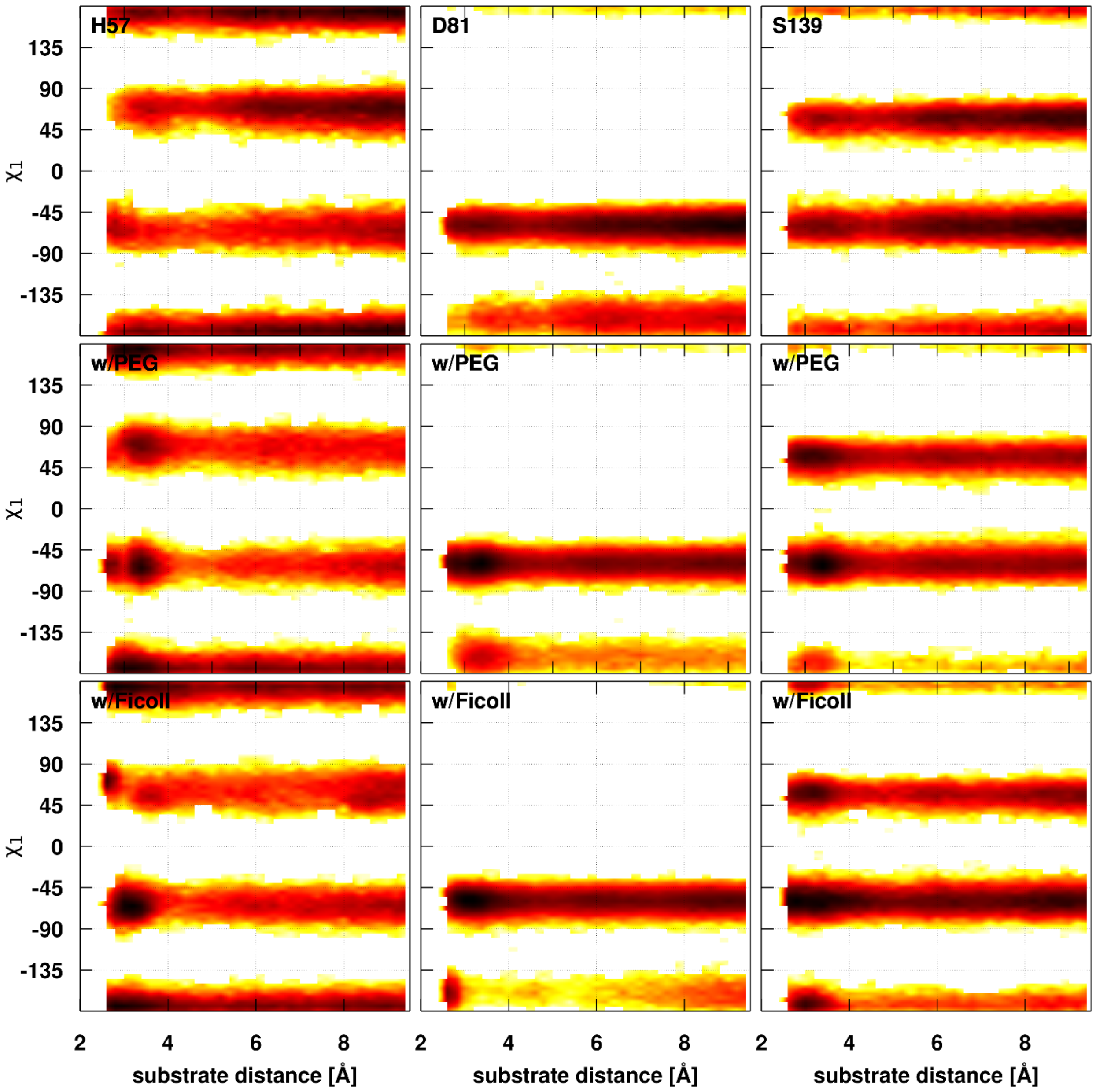


**Figure S20.** Side chain $\chi_1$ torsion angle sampling of active site residues as a function of substrate distances. Results are shown for simulations with only substrate (top row) and with PEG (middle row) or Ficoll (bottom row) crowders as in **Figure S18**.

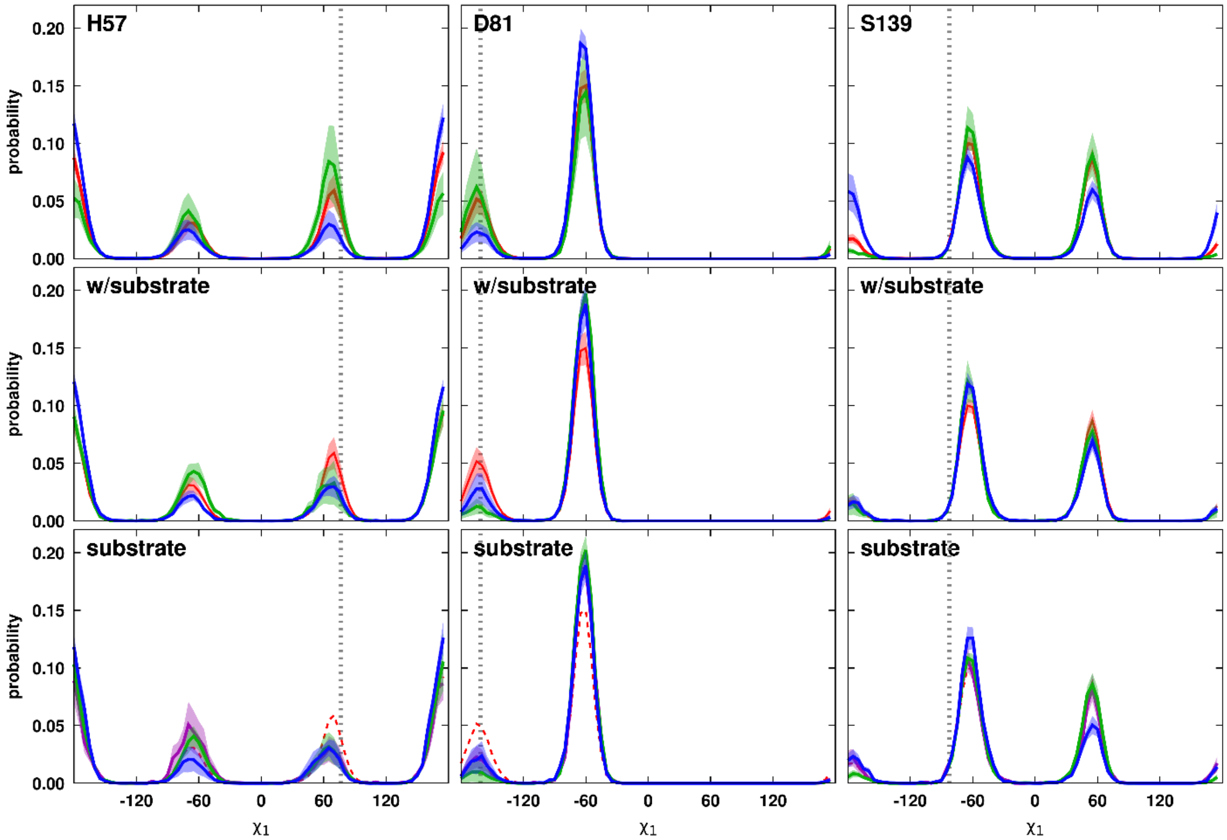


**Figure S21.** Sampling of side chain $\chi_1$ torsion angles for active site residues in water and when crowders or substrates are in close contact. Contacts were defined as a minimum distance of 5 Å. Results for simulations with water (red) or only crowders (PEG: green, Ficoll: blue) are shown in the top row. Results for simulations with substrate but based on close crowder contacts are shown in the middle row. Results based on substrates being in close contact are shown in the bottom row. In the bottom row, the results from the simulations with only substrates are shown in purple. The dilute water distributions are shown as a dashed line for reference. Shaded areas indicate standard errors.

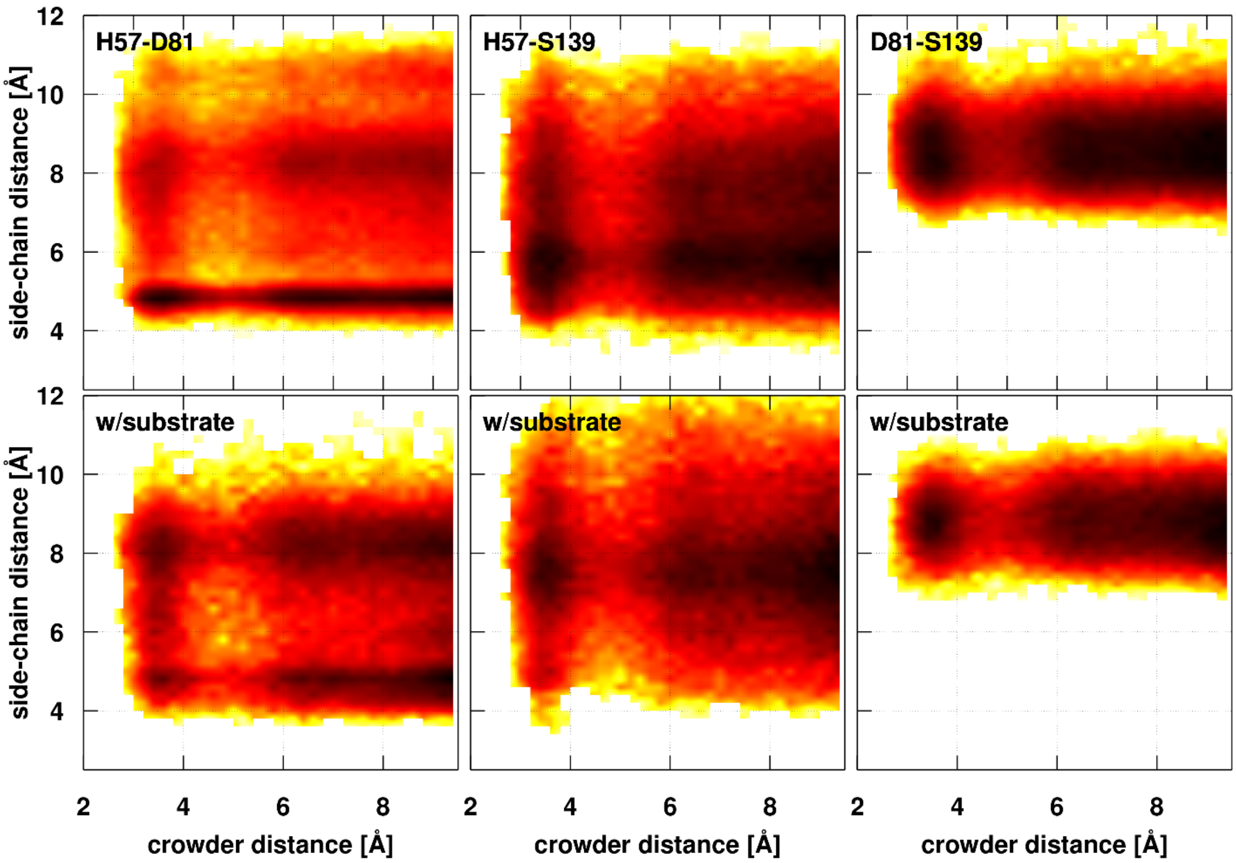


**Figure S22.** Side chain center distances between active site residues as a function of PEG crowder distances. See **Figure S18** for further details. In the crystal structure (PDB ID: 4JMY), the distances are 4.45 Å for His57-Asp81, 4.69 Å for His57-Ser139, and 7.37 Å for Asp81-Ser139.

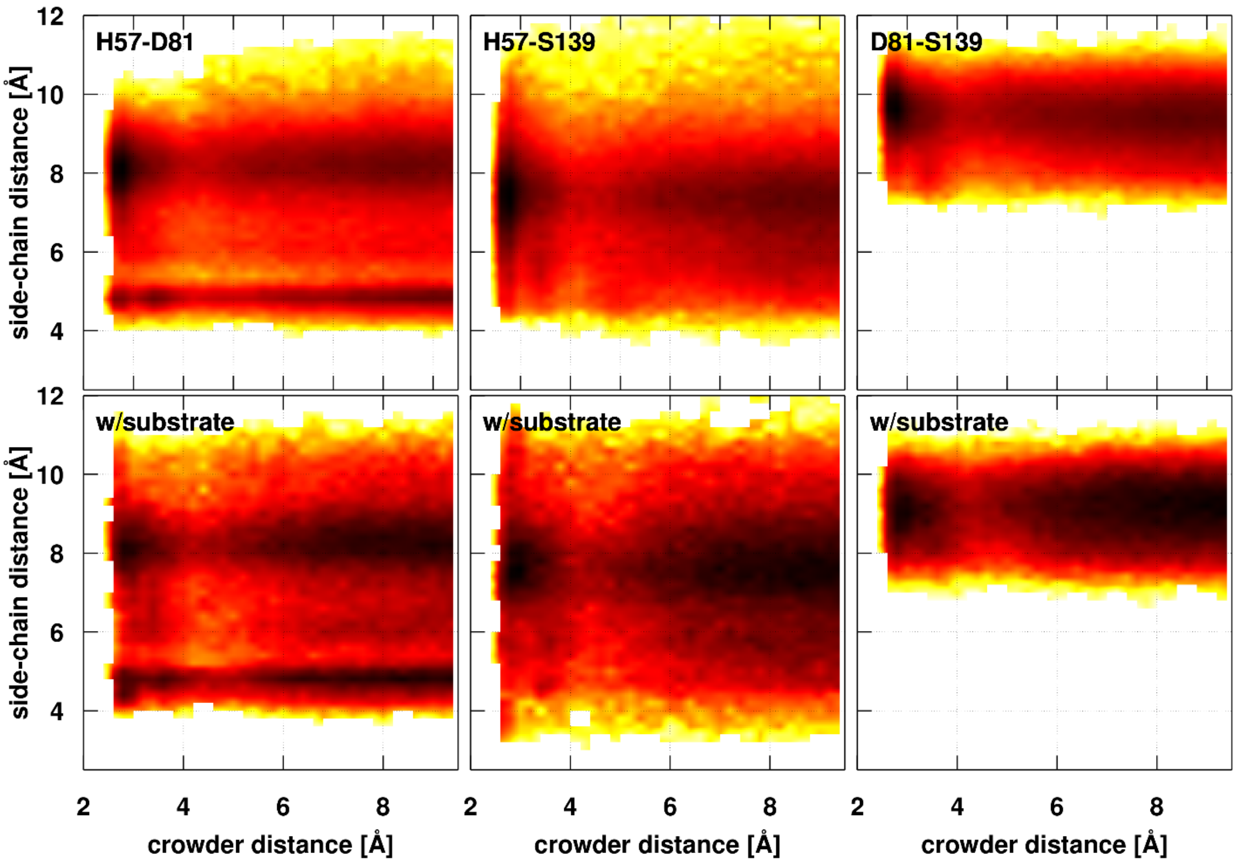


**Figure S23.** Side chain center distances between active site residues as a function of Ficoll crowder distances. See **Figures S18/S22** for further details.

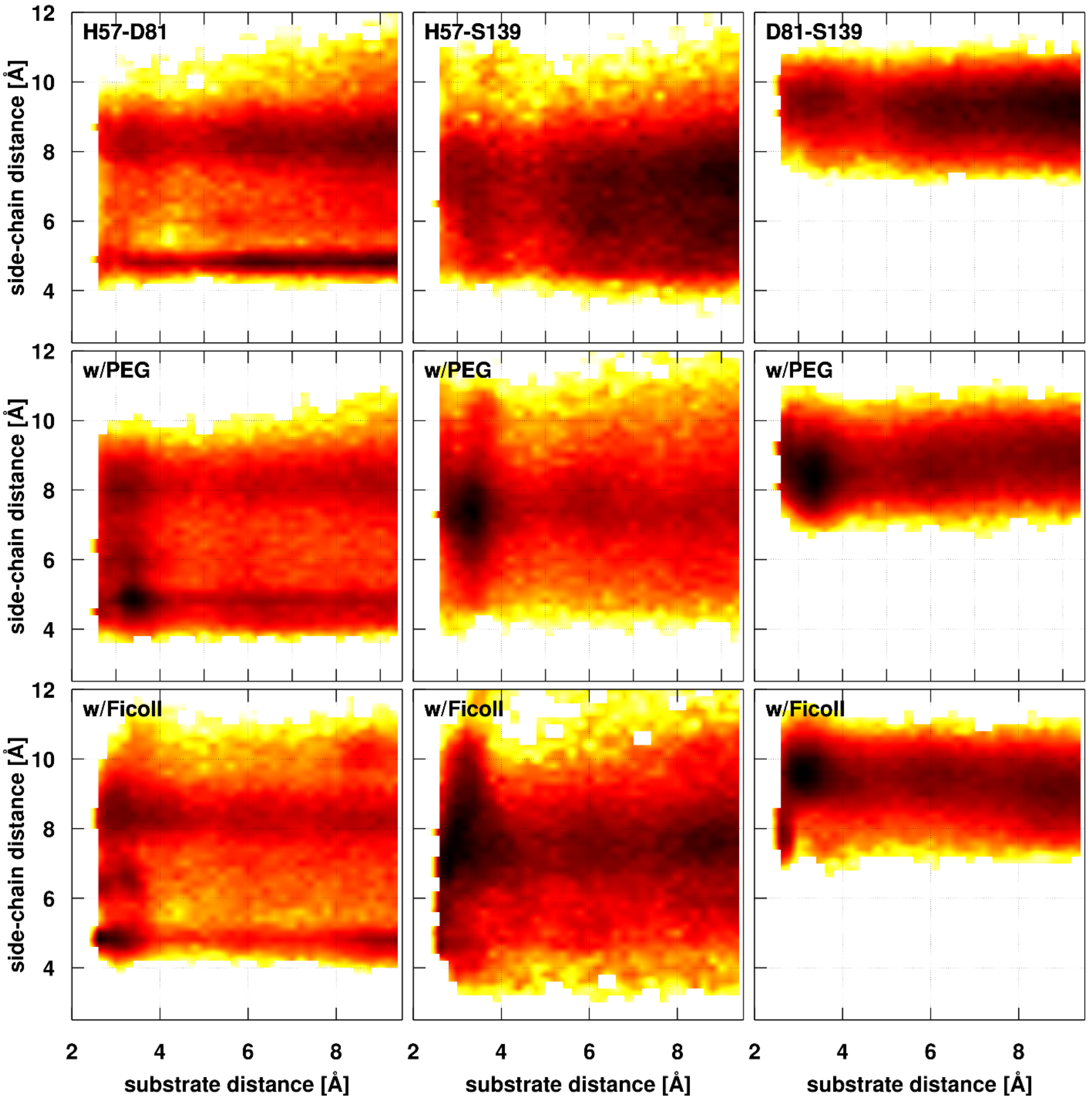


**Figure S24.** Side chain center distances between active site residues as a function of substrate distances. Results are shown for simulations with only substrates (top row) and with PEG (middle row) or Ficoll (bottom row) crowders as in **Figure S22**.

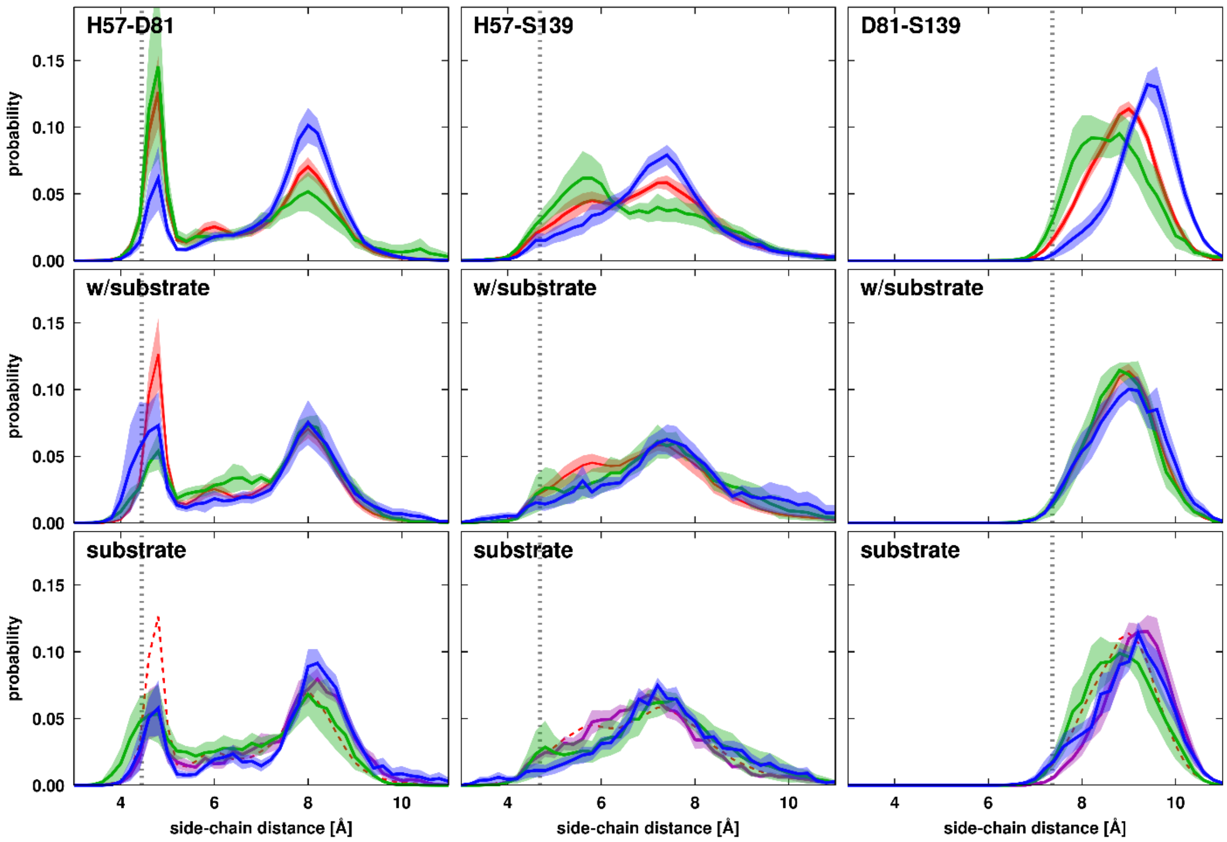


**Figure S25.** Sampling of side chain center distances between active site residues in water and when crowders or substrates are in close contact**.** See **Figure S21** for further details.

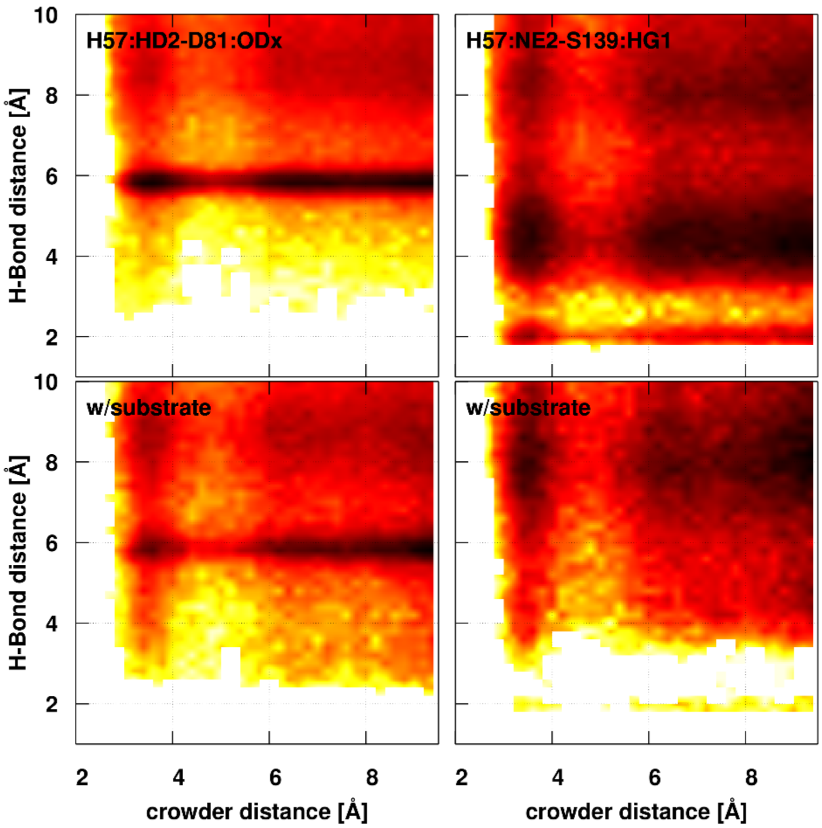


**Figure S26.** Catalysis-relevant distances between active site residues (His57:Nδ-H … Oδ1/2-Asp81 and His57-Nε … HOγ-Ser139) as a function of PEG crowder distances. Results are shown for simulations without (top row) and with (bottom row) substrates.

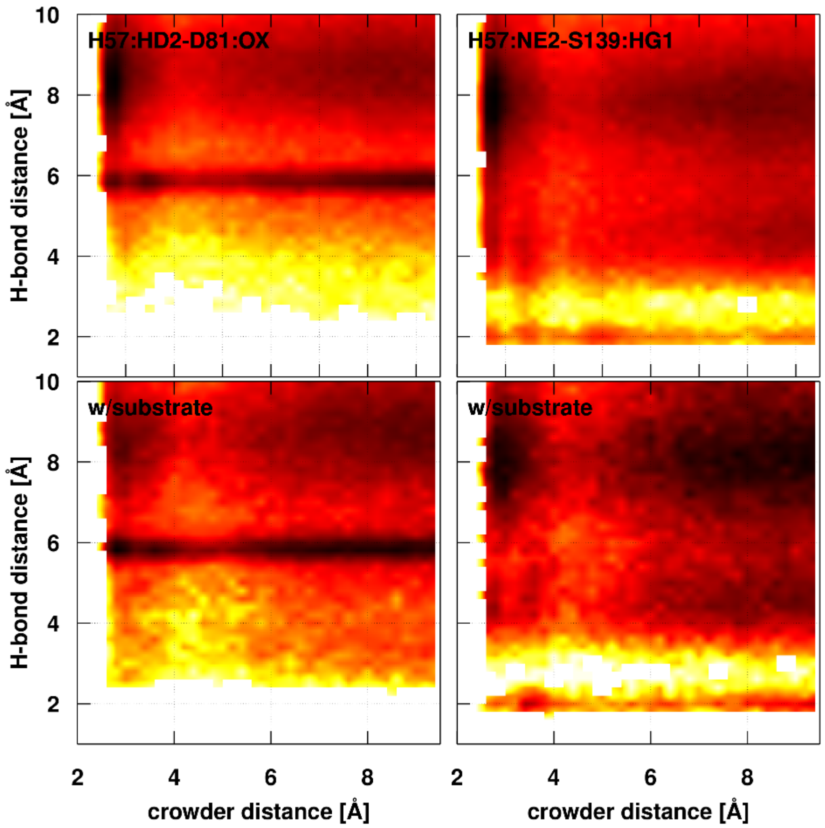


**Figure S27.** Catalysis-relevant distances between active site residues (His57:Nδ-H … Oδ1/2-Asp81 and His57-Nε … HOγ-Ser139) as a function of Ficoll crowder distances. See details in **Figure S26**.

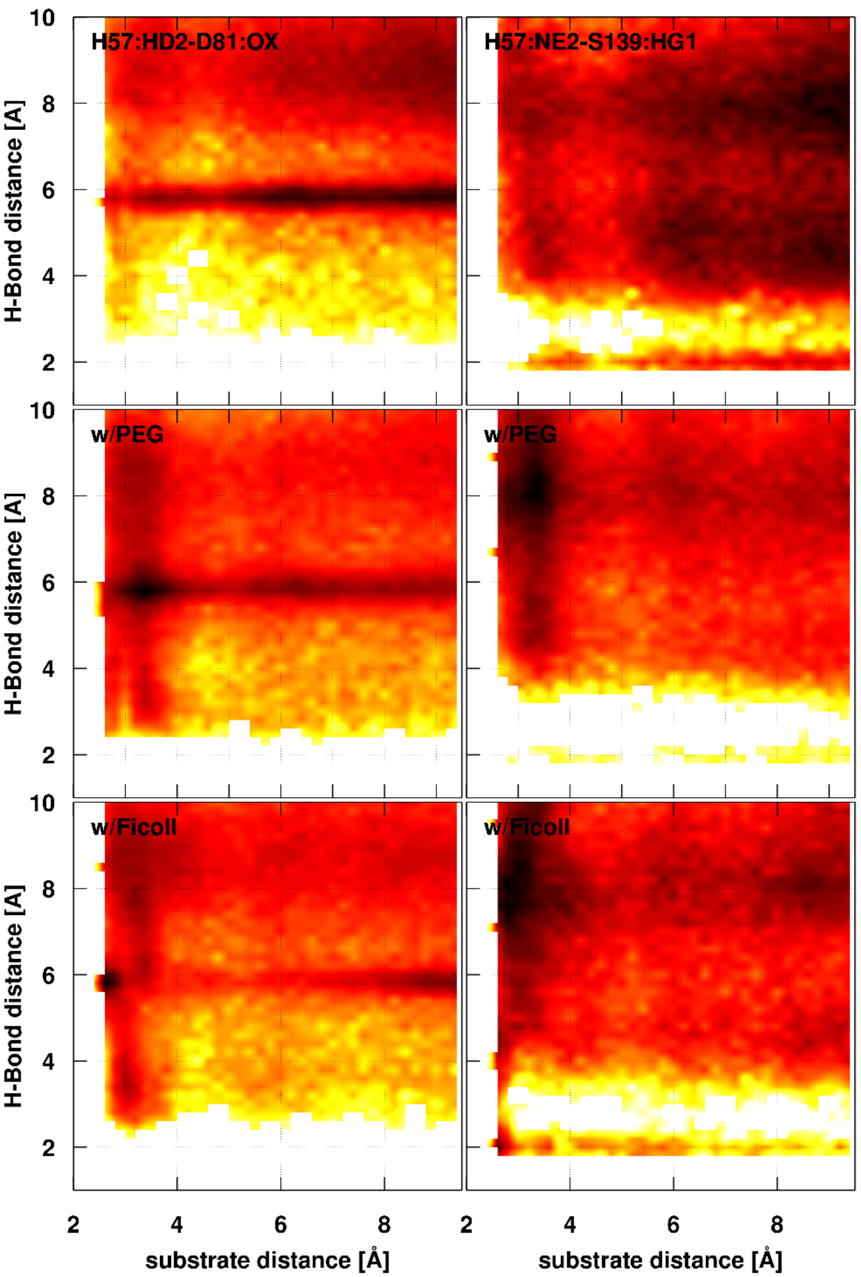


**Figure S28.** Catalysis-relevant distances between active site residues (His57:Nδ-H … Oδ1/2-Asp81 and His57-Nε … HOγ-Ser139) as a function of substrate distances. Results are shown for simulations with only substrate (top row) and with PEG (middle row) or Ficoll (bottom row) crowder as in **Figure S26**.

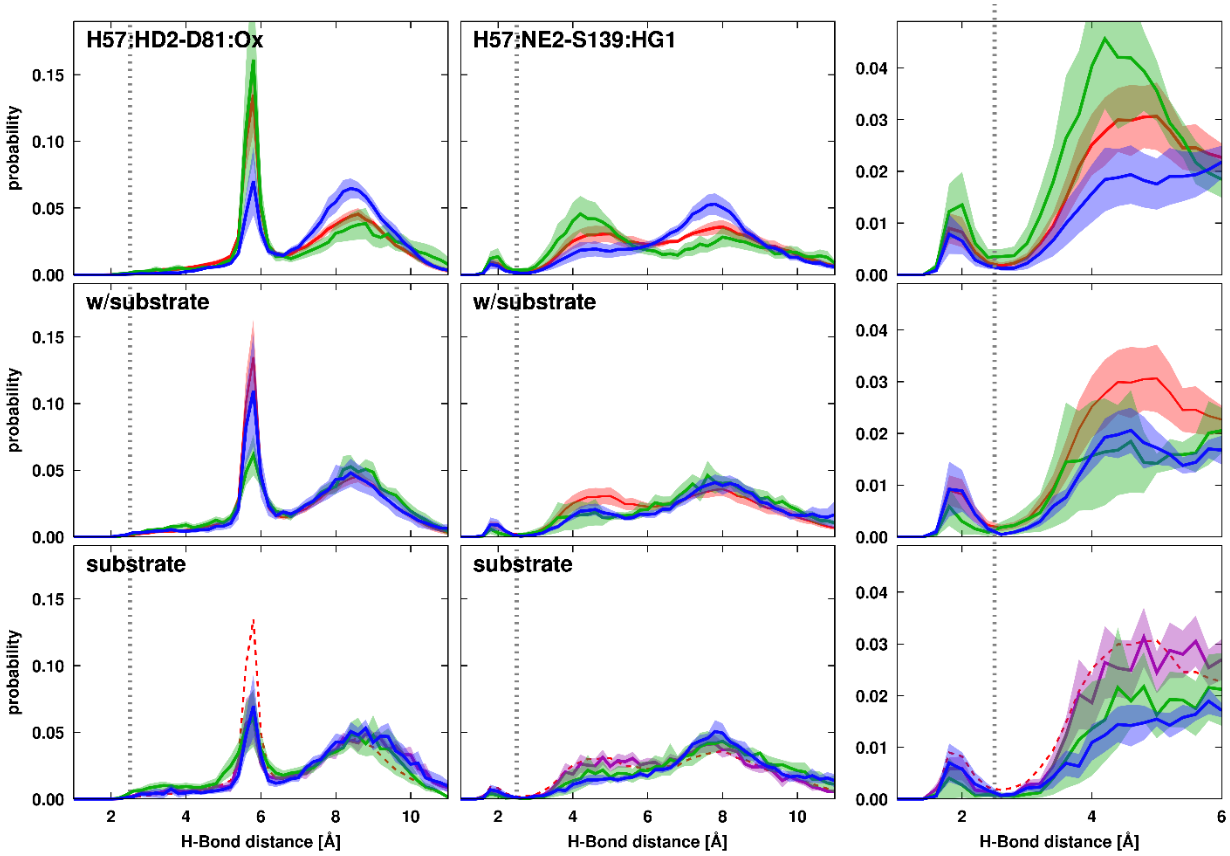


**Figure S29.** Sampling of catalytically relevant active site distances in water and when crowders or substrates are in close contact**.** See **Figure S21** for further details. The right-most column zooms in on the shorter-distance region of the data shown in the middle column.

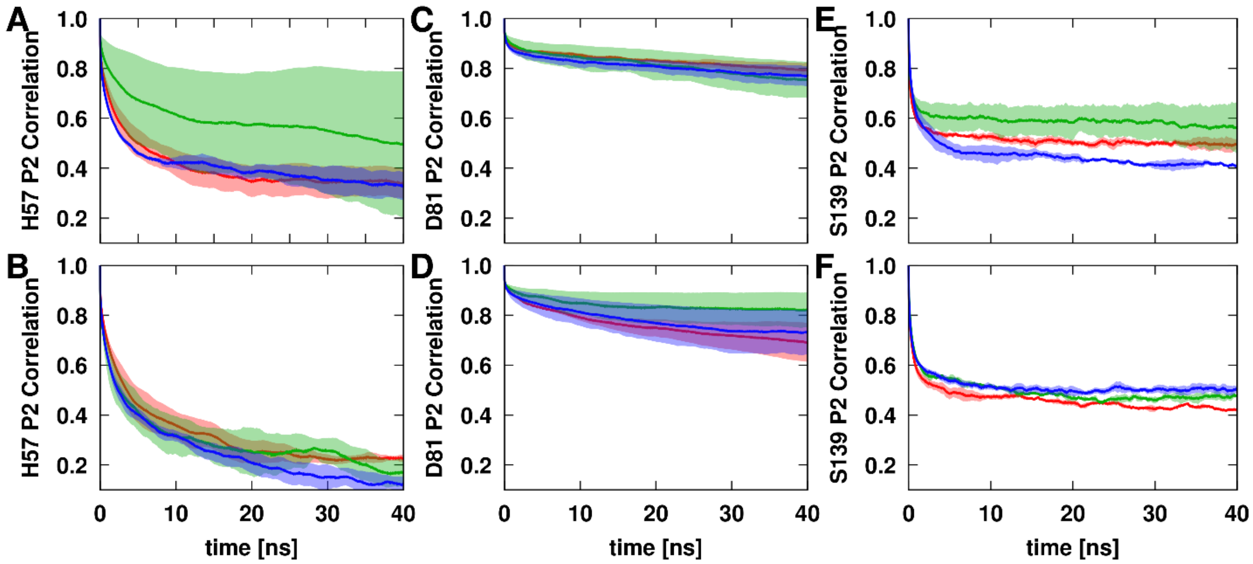


**Figure S30.** Orientational correlation function for active site side chains (His57, Asp81, Ser139) in NS3. Fluctuations were calculated based on the Cα-Nε vector (His57, A/B), the Cα-Cγ vector (Asp81, C/D), and the Cα-Oγ vector (Ser139, E/F) after superposition of the overall NS3 structure to a common reference in water (red) or in the presence of Ficoll (blue) or PEG (green) crowder without (A/C/E) and with (B/D/F) substrates. The correlation function was calculated as the second-order Legendre polynomial from the inner dot product of the orientational vector at different time points. Solid lines show simulation averages, standard errors are indicated by the shaded area. Results from double exponential fits to the correlation functions are given in **Table S4**. Only trajectories with the reduced friction were used in this analysis (**Table S1**).

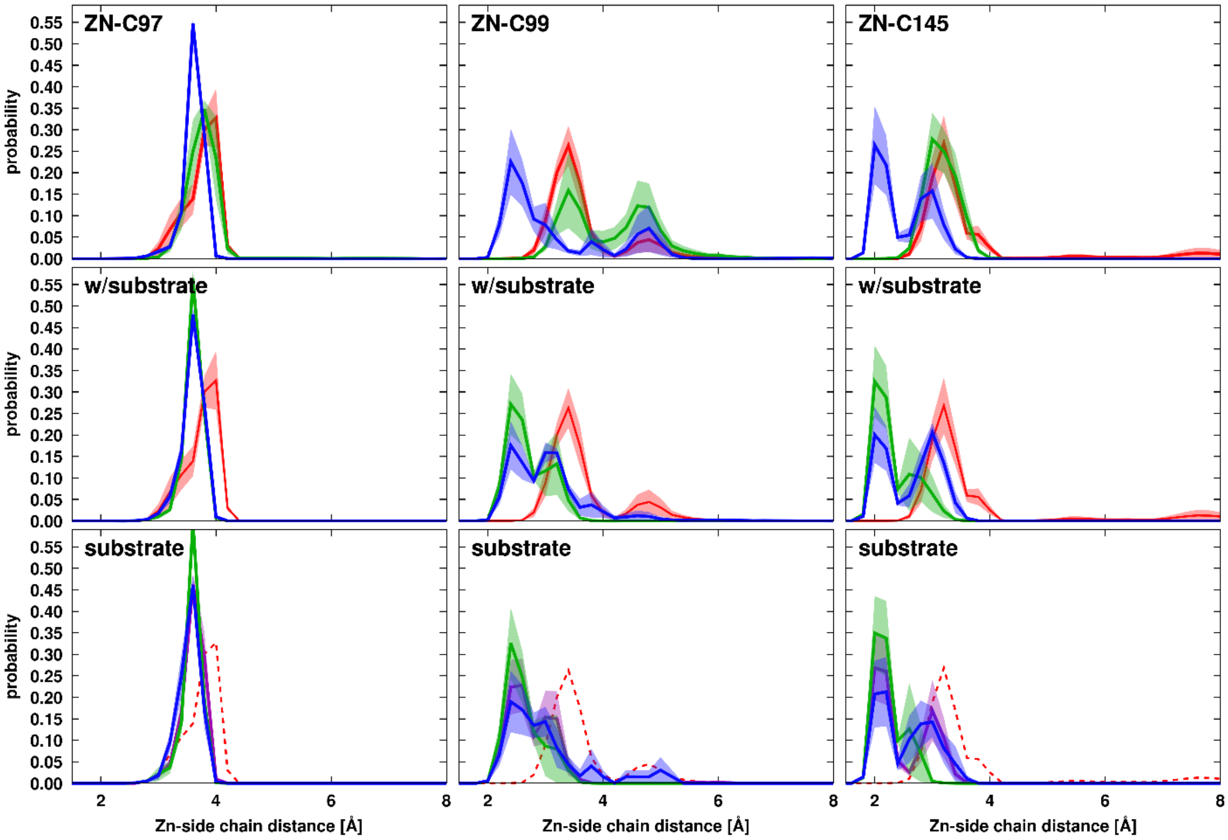


**Figure S31.** Sampling of distances between the $Zn^{2+}$ ion and the coordinating cysteine residues of NS3 in water and when crowders or substrates are in close contact**.** See **Figure S21** for further details.

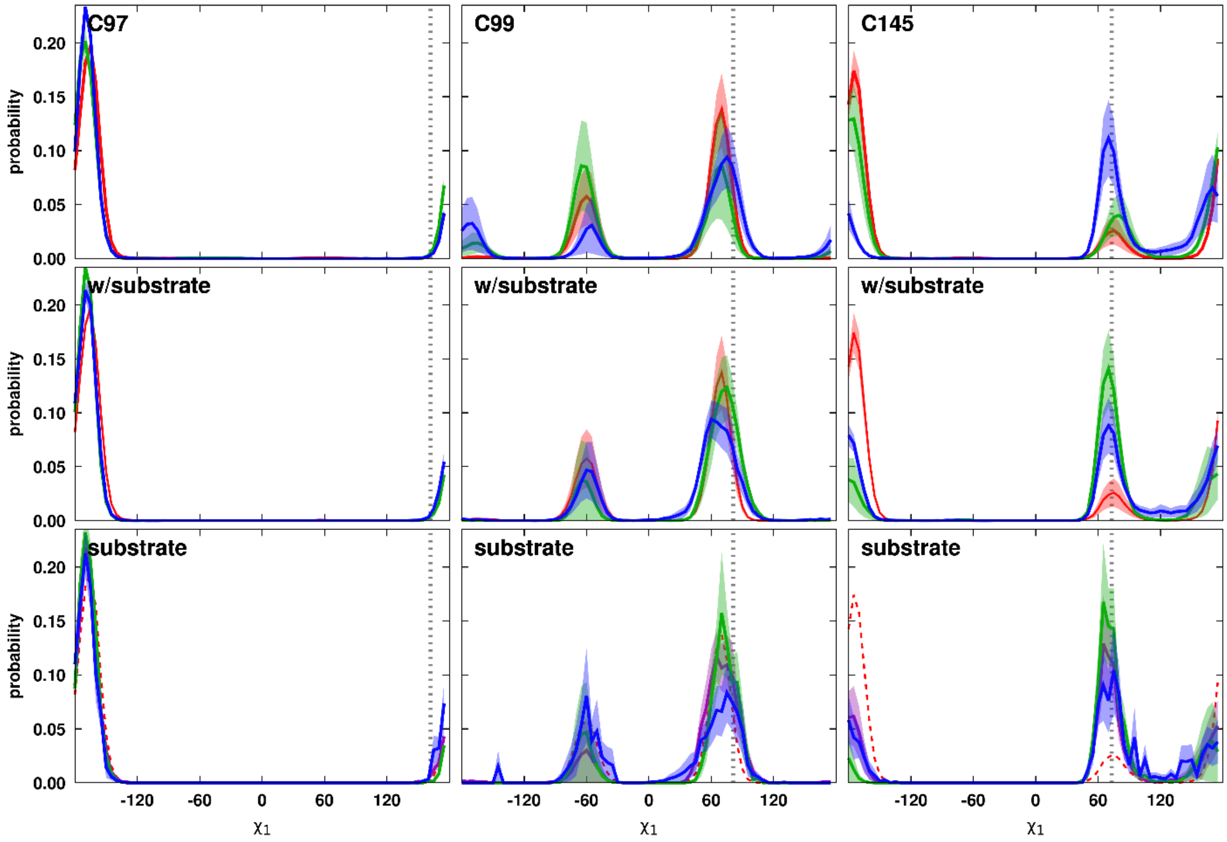


**Figure S32.** Sampling of side chain torsion $\chi_1$ angles for $Zn^{2+}$ coordinating cysteine residues of NS3 in water and when crowders or substrates are in close contact. Torsion angles present in the crystal structure (4JMY) are indicated by vertical dotted lines. See **Figure S21** for further details.

**Table S1**. Simulations of NS3/4A under different conditions analyzed in this study.

| System | Time[2] [ns] | Frames[3] | Atoms | PEG | Ficoll | Substr. | $H_2O$ | $Na^+$ | $Cl^-$ | Box[4] [$Å^3$] |
|---|---|---|---|---|---|---|---|---|---|---|
| **W1** | 555 | 11108 | 128726 | 0 | 0 | 0 | 41718 | 20 | 22 | 108.5 x 108.5 x 108.5 |
| **W2** | 576 | 11522 | 128726 | 0 | 0 | 0 | 41718 | 20 | 22 | 108.5 x 108.5 x 108.5 |
| **W3** | 538 | 10770 | 128726 | 0 | 0 | 0 | 41718 | 20 | 22 | 108.5 x 108.5 x 108.5 |
| **W1x[1]** | 502 | 50200 | 128726 | 0 | 0 | 0 | 41718 | 20 | 22 | 108.7 x 108.7 x 108.7 |
| **W2x[1]** | 502 | 50200 | 128726 | 0 | 0 | 0 | 41718 | 20 | 22 | 108.7 x 108.7 x 108.7 |
| **W3x[1]** | 502 | 50200 | 128726 | 0 | 0 | 0 | 41718 | 20 | 22 | 108.7 x 108.7 x 108.7 |
| **P1** | 567 | 11348 | 258455 | 130 | 0 | 0 | 76031 | 25 | 27 | 142.9 x 132.4 x 132.2 |
| **P2** | 500 | 50000 | 258455 | 130 | 0 | 0 | 76031 | 25 | 27 | 144.2 x 133.5 x 133.4 |
| **P3** | 500 | 50000 | 258455 | 130 | 0 | 0 | 76031 | 25 | 27 | 144.2 x 133.5 x 133.4 |
| **P1x[1]** | 502 | 50200 | 258455 | 130 | 0 | 0 | 76031 | 25 | 27 | 144.2 x 133.6 x 133.4 |
| **P2x[1]** | 502 | 50200 | 258455 | 130 | 0 | 0 | 76031 | 25 | 27 | 144.2 x 133.6 x 133.4 |
| **F1** | 502 | 50200 | 260298 | 0 | 110 | 0 | 78084 | 37 | 39 | 134.7 x 135.7 x 140.4 |
| **F2** | 502 | 50200 | 256218 | 0 | 110 | 0 | 76724 | 37 | 39 | 136.7 x 135.7 x 136.1 |
| **F2** | 502 | 50200 | 260891 | 0 | 110 | 0 | 78281 | 38 | 40 | 137.1 x 137.3 x 136.6 |
| **F1x[1]** | 502 | 50200 | 260298 | 0 | 110 | 0 | 78084 | 37 | 39 | 134.7 x 135.7 x 140.5 |
| **F2x[1]** | 502 | 50200 | 256218 | 0 | 110 | 0 | 76724 | 37 | 39 | 136.7 x 135.7 x 136.1 |
| **F3x[1]** | 502 | 50200 | 260891 | 0 | 110 | 0 | 78281 | 38 | 40 | 137.2 x 137.4 x 136.6 |
| **S1** | 528 | 10556 | 147865 | 0 | 0 | 10 | 47647 | 61 | 23 | 103.3 x 122.5 x 113.2 |
| **S2** | 666 | 13321 | 108274 | 0 | 0 | 10 | 34454 | 55 | 17 | 104.2 x 96.2 x 104.6 |
| **S3** | 548 | 10951 | 103337 | 0 | 0 | 10 | 32809 | 54 | 16 | 99.3 x 97.8 x 102.9 |
| **S1x[1]** | 502 | 50200 | 147865 | 0 | 0 | 10 | 47647 | 61 | 23 | 104.5 x 123.5 x 114.3 |
| **S2x[1]** | 502 | 50200 | 108274 | 0 | 0 | 10 | 34454 | 55 | 17 | 105.1 x 97.2 x 105.5 |
| **S3x[1]** | 502 | 50200 | 103337 | 0 | 0 | 10 | 32809 | 54 | 16 | 100.2 x 98.8 x 103.9 |
| **PS1** | 562 | 11232 | 246934 | 130 | 0 | 10 | 71736 | 72 | 34 | 143.1 x 132.7 x 125.6 |
| **PS2** | 505 | 10103 | 241244 | 130 | 0 | 10 | 69840 | 71 | 33 | 145.8 x 134.1 x 119.2 |
| **PS1x[1]** | 502 | 50200 | 246934 | 130 | 0 | 10 | 71736 | 72 | 34 | 144.4 x 133.9 x 126.8 |
| **PS2x[1]** | 502 | 50200 | 241244 | 130 | 0 | 10 | 69840 | 71 | 33 | 147.1 x 135.3 x 120.3 |
| **FS1** | 518 | 10364 | 244759 | 0 | 110 | 10 | 72457 | 73 | 35 | 131.7 x 132.5 x 137.2 |
| **FS2** | 549 | 10975 | 232827 | 0 | 110 | 10 | 68481 | 71 | 33 | 133.0 x 130.7 x 131.0 |
| **FS3** | 537 | 10740 | 235800 | 0 | 110 | 10 | 69472 | 71 | 33 | 132.6 x 131.8 x 132.0 |
| **FS1x[1]** | 502 | 50200 | 244759 | 0 | 110 | 10 | 72457 | 73 | 35 | 132.0 x 132.7 x 137.5 |
| **FS2x[1]** | 502 | 50200 | 232827 | 0 | 110 | 10 | 68481 | 71 | 33 | 133.2 x 130.9 x 131.2 |
| **FS3x[1]** | 502 | 50200 | 235800 | 0 | 110 | 10 | 69472 | 71 | 33 | 132.9 x 132.0 x 132.2 |

[1]Simulations with a reduced Langevin friction coefficient of 0.01 $ps^{-1}$

[2]Total simulation time.

[3]Total number of snapshots saved during simulation.

[4]Simulation-averaged

**Table S2**: Average Cα coordinate root mean square deviations of NS3/4A

| System | NS3/4A RMSD [Å] |
|---|---|
| **Water** | 1.573 (0.075) |
| **PEG** | 1.639 (0.050) |
| **Ficoll** | 1.647 (0.101) |
| **Substrate** | 1.578 (0.067) |
| **PEG/Substrate** | 1.697 (0.113) |
| **Ficoll/Substrate** | 1.480 (0.073) |

RMSD values were calculated after optimal superposition with respect to the experimental structure (PDB ID: 4JMY). Only the part of NS4A resolved in the PDB structure was considered. Averages based on all trajectories for a given system with standard errors of the mean in parentheses.

**Table S3**: Average radius of gyration for NS3, NS4A, substrate, and crowders from heavy atoms

| System | NS3<br>$R_g$ [Å] | NS4A<br>$R_g$ [Å] | Substrate<br>$R_g$ [Å] | Ficoll<br>$R_g$ [Å] | PEG<br>$R_g$ [Å] |
|---|---|---|---|---|---|
| **Water** | 16.22 (0.03) | 24.89 (1.28) | | | |
| **PEG** | 16.23 (0.03) | 25.98 (0.94) | | | 12.56 (0.03) |
| **Ficoll** | 16.27 (0.02) | 24.17 (1.15) | | 8.09 (0.001) | |
| **Substrate** | 16.25 (0.02) | 25.77 (0.53) | 7.81 (0.06) | | |
| **PEG/Substrate** | 16.21 (0.03) | 24.59 (1.82) | 7.68 (0.07) | | 12.57 (0.01) |
| **Ficoll/Substrate** | 16.21 (0.02) | 26.78 (1.27) | 7.64 (0.02) | 8.08 (0.002) | |

Averages based on all trajectories for a given system with standard errors of the mean in parentheses.

**Table S4**: Average secondary structure content for the NS4A and substrate calculated based on VMD Timeline results (the helix % includes both α and $3_{10}$ helix structures).

| System | NS4A | | Substrate | |
|---|---|---|---|---|
| | **helix [%]** | **β-sheet [%]** | **helix [%]** | **β-sheet [%]** |
| **Water** | 1.68 (0.76) | 2.24 (0.19) | | |
| **Ficoll** | 1.35 (0.95) | 2.51 (0.10) | | |
| **PEG** | 5.64 (1.31) | 2.03 (0.10) | | |
| **Substrate** | 1.75 (0.28) | 2.66 (0.29) | 3.45 (1.31) | 0 |
| **Ficoll/Substrate** | 5.58 (1.99) | 3.19 (1.00) | 5.47 (1.09) | 0 |
| **PEG/Substrate** | 8.30 (4.51) | 2.62 (0.33) | 3.04 (0.94) | 0 |

Averages based on all trajectories for a given system with standard errors of the mean in parentheses.

**Table S5**: Double-exponential fits to rotational correlation functions

| | NS3/4A | | | Substrate | | |
|---|---|---|---|---|---|---|
| | **$\tau_s$ [ns]** | **$\tau_f$ [ns]** | **$s_r$** | **$\tau_s$ [ns]** | **$\tau_f$ [ns]** | **$s_r$** |
| **Water** | 22.72 | 3.74 | 0.74 | | | |
| **PEG** | 28.22 | 3.43 | 0.72 | | | |
| **Ficoll** | 46.72 | 6.89 | 0.59 | | | |
| **Substrate** | 48.98 | 5.32 | 0.53 | 61.94 | 0.52 | 0.11 |
| **PEG/Substrate** | 4285.8 | 13.50 | 0.26 | 46.44 | 0.57 | 0.14 |
| **Ficoll/Substrate** | 34.42 | 4.78 | 0.79 | 52.23 | 0.60 | 0.12 |

Time decays and weight of slow-time scale component (see Methods) from double-exponential fits to combined rotational correlation functions.

**Table S6.** NS3-crowder contact life-times from two-exponential fits to contact survival decays

| | $\tau_1$ [ns] | $\tau_2$ [ns] | a | $\chi^2$ |
|---|---|---|---|---|
| **PEG** | 0.28 (0.09) | 6.71 (1.61) | 0.58 (0.045) | 0.851 |
| **PEG w/substrate** | 0.27 (0.03) | 6.37 (0.41) | 0.56 (0.002) | 0.727 |
| **Ficoll** | 0.23 (0.02) | 4.17 (0.23) | 0.52 (0.025) | 0.565 |
| **Ficoll w/substrate** | 0.24 (0.04) | 4.43 (0.41) | 0.50 (0.013) | 0.557 |
| **Substrate** | 1.93 (1.78) | 47.0 (28.4) | 0.42 (0.008) | 1.884 |
| **Substrate w/PEG** | 2.88 (0.93) | 94.3 (5.8) | 0.29 (0.050) | 1.406 |
| **Substrate w/Ficoll** | 1.48 (1.44) | 48.9 (35.1) | 0.43 (0.052) | 1.345 |

Averages over replicate trajectories with reduced friction coefficients. Standard errors are given in parentheses.

**Table S7.** Substrate-crowder contact life-times from two-exponential fits to contact survival decays

| | $\tau_1$ [ns] | $\tau_2$ [ns] | a | $\chi^2$ |
|---|---|---|---|---|
| **PEG** | 0.073 (0.002) | 1.50 (0.05) | 0.71 (0.000) | 0.255 |
| **Ficoll** | 0.106 (0.001) | 1.74 (0.03) | 0.60 (0.001) | 0.254 |

Averages over replicate trajectories with standard errors given in parentheses.

**Table S8**: Double-exponential fits to the combined orientational correlation functions of the His57, Asp81, and Ser139 active site side chains

| | **His57** | | | **Asp81** | | | **Ser139** | | |
|---|---|---|---|---|---|---|---|---|---|
| | **$\tau_1$ [ns]** | **$\tau_2$ [ns]** | **a** | **$\tau_1$ [ns]** | **$\tau_2$ [ns]** | **a** | **$\tau_1$ [ns]** | **$\tau_2$ [ns]** | **a** |
| **Water** | 129.5 | 1.07 | 0.44 | 403.7 | 0.257 | 0.88 | 455.7 | 0.319 | 0.54 |
| **PEG** | 141.8 | 1.53 | 0.67 | 247.1 | 0.331 | 0.88 | 509.4 | 0.269 | 0.61 |
| **Ficoll** | 108.4 | 0.99 | 0.47 | 303.8 | 0.363 | 0.85 | 214.4 | 0.717 | 0.49 |
| **Substrate** | 55.2 | 1.86 | 0.41 | 211.2 | 1.205 | 0.83 | 209.7 | 0.447 | 0.50 |
| **PEG/Substrate** | 46.6 | 1.22 | 0.41 | 783.3 | 0.911 | 0.86 | 296.7 | 0.549 | 0.53 |
| **Ficoll/Substrate** | 28.7 | 1.06 | 0.45 | 263.4 | 0.945 | 0.84 | 521.9 | 0.497 | 0.53 |
